# The T cell epitope landscape of SARS-CoV-2 variants of concern

**DOI:** 10.1101/2022.06.06.491344

**Authors:** Simen Tennøe, Marius Gheorghe, Richard Stratford, Trevor Clancy

**Affiliations:** NEC OncoImmunity AS, Oslo Cancer Cluster, Ullernchausseen 64/66, 0379 Oslo, Norway

## Abstract

During the COVID-19 pandemic, several SARS-CoV-2 variants of concern (VOC) emerged, bringing with them varying degrees of health and socioeconomic burdens. In particular, the Omicron VOC displayed distinct features of increased transmissibility accompanied by antigenic drift in the spike protein that partially circumvented the ability of pre-existing antibody responses in the global population to neutralize the virus. However, T cell immunity has remained robust throughout all the different VOC transmission waves and has emerged as a critically important correlate of protection against SARS-CoV-2 and it’s VOCs, in both vaccinated and infected individuals. Therefore, as SARS-CoV-2 VOCs continue to evolve, it is crucial that we characterize the correlates of protection and the potential for immune escape for both B cell and T cell human immunity in the population. Generating the insights necessary to understand T cell immunity, experimentally, for the global human population is at present critical but a time consuming, expensive, and laborious process. Further, it is not feasible to generate global or universal insights into T cell immunity in an actionable time frame for potential future emerging VOCs. However, using computational means we can expedite and provide early insights into the correlates of T cell protection. In this study, we generated and reveal insights on the T cell epitope landscape for the five main SARS-CoV-2 VOCs observed to date. We demonstrated here using a unique AI prediction platform, a strong concordance in global T cell protection across all mutated peptides for each VOC. This was modeled using the most frequent HLA alleles in the human population and covers the most common HLA haplotypes in the human population. The AI resource generated through this computational study and associated insights may guide the development of T cell vaccines and diagnostics that are even more robust against current and future VOCs, and their emerging subvariants.

## INTRODUCTION

During the COVID-19 pandemic ^1^, we experienced several SARS-CoV-2 variants that caused various degrees of altered infectivity, potential immune escape, or both, compared to the original wildtype Wuhan strain. Some of these variants had characteristic mutations that changed the epidemiology of the pandemic, and indeed some of these variants altered the clinical impact of COVID-19. There were five main lineages that were designated as SARS-CoV-2, so called, “variants of concern” (VOC). These lineages systematically ushered in new waves of transmission, changing the nature of the pandemic, and were accompanied by different health and socioeconomic challenges. These five VOCs (Alpha, Beta, Gamma, Delta, and Omicron) shared several mutations, as well as harboring several VOC-specific variants. The most recent VOC that emerged, the hyper-transmissible Omicron, was a presage to us all that VOCs will continue to emerge and pose a continuous health and socioeconomic threat, particularly now, as the SARS-CoV-2 virus now becomes endemic in the human population ^2^.

The VOCs were primarily characterized based on their increased infectivity through specific mutations within the receptor binding domain (RBD) of the spike protein with key properties that often rendered the mutated RBDs with either; 1) increased potential to enter host cells through increased binding affinity for its cognate receptor, or 2) evasion of neutralizing antibody protection through a diminished capacity of hosts antibodies to bind to the mutated RBD and inhibit its ability to bind to the host receptor ^3^. A more deadly variant that confers both increased infectivity and evasion of vaccine or infection induced immune protection will create very difficult challenges ^2, 4^. Therefore, developing broadly protective vaccines with a focus on emerging virulent VOCs will be important to ensure the human population maintains protective immunity against future emerging SARS-CoV-2 VOCs ^5^.

In the quest to develop broadly protective vaccines against emerging VOCs, researchers have increasingly looked beyond the traditional antibody-centric approaches and begun to focus on characterizing the requisite T cell responses that correlate with robust and lasting protection against COVID-19 ^6^. Although neutralizing antibody protection from infection or vaccination is the current gold standard, there was widespread escape from pre-existing vaccine induced, or naturally acquired, neutralizing antibody responses during the Omicron transmission wave ^7^. Interestingly, many infected SARS-CoV-2 individuals during the COVID-19 pandemic demonstrated virus-specific T cell responses in the absence of measurable virus specific antibodies ^8, 9^. Numerous studies during the pandemic demonstrated that T cell immunity played a critical role in protection against the virus ^10, 11^. For example, a systems immunology approach that analyzed high dimensional molecular data from 139 COVID-19 patients representing various degrees of disease severity, revealed that CD8+ T cells were associated with improved clinical outcome ^12^. Additionally, common T cell epitopes were identified between SARS-CoV-2 and human coronaviruses (HCoV) and T cell responses against these cross-reactive epitopes were important for the observed clinical protection against COVID-19 ^13, 14, 15, 16^. Although individual viral mutations do have potential to diminish or destroy effector T cell responses at the level of the individual by destroying important HLA-restricted epitopes, it is highly unlikely that these same mutations will confer a selective advantage (via immune escape) in other individuals with different HLA haplotypes, or at the level of a population, which has a highly diverse HLA haplotype landscape. This rational is further strengthened when we consider the size of the SARS-CoV-2 RNA genome which is approximately 30kb encoding 14 open reading frames (some of which are overlapping) and offers ample protein “real estate” for HLA restricted T cell immunity. Consequently, T-cells can exploit the whole viral proteome, and are not limited to the real estate offered by the spike protein, particularly the RBD domain, unlike the humoral response. Furthermore, T cell protection is sourced not only from the broad spectrum of the SARS-CoV-2 viral proteome, but also from across the entire betacoronavirus family. For example, T cell immunity reported during the Middle Eastern respiratory virus (MERS) outbreak was demonstrated to be more robust compared to the B cell antibody response ^17^.

These findings and observations strongly suggest that T cell immunity is a key correlate of protection against beta-coronavirus infections including SARS-CoV-2 and it’s VOCs, and that adopting vaccine strategies that can drive broader and more potent T-cell responses will be the key to developing more efficacious and broadly protective vaccines in the future. Whilst neutralizing antibody responses are essential to prevent transmission and provide sterile immunity, they are short-lived, as has been observed for SARS-CoV-1, SARS-CoV-2 and the seasonal coronaviruses ^18^. For example, while antibody responses were shown to be short-lived in patients infected with SARS-CoV ^19, 20^, the corresponding T cell responses in the same patients were detected 17 years after infection ^19^. Interestingly, the current mRNA-based spike-centric vaccines against SARS-CoV-2 have been shown to induce broad T cell responses that recognize several SARS-CoV-2 variants and HCoVs ^21^ in addition to driving neutralizing antibody responses. T cell responses were measured and detected in most vaccine studies during the COVID-19 pandemic ^22^. In fact, protective clinical benefits of vaccination were seen as early as 11 days after first vaccination coinciding with a robust T cell response ^23^. Even before the emergence of the most recent VOC, Omicron, T cell responses induced by vaccines demonstrated strong cross-protection against different VOC ^24^.

In terms of the most recent hyper-transmissible Omicron VOC, its increased infectivity cannot be explained by a higher viral load alone ^25, 26^, pointing to immune evasion as an explanation ^27^. Further, it was demonstrated very early in Omicron’s rapid spread that it evades antibody neutralization from both vaccinated and convalescent individuals ^28, 29, 30, 31, 32^. In addition, the large number of new mutations in the Omicron VOC (32 mutations in the spike protein alone) also raised fears that these changes might enable the virus to circumvent pre-existing T-cell immunity induced by the vaccine or by natural infection with earlier VOCs. However, despite Omicron’s extensive number of mutations and its ability to escape from neutralizing antibodies in vaccinated and convalescent individuals, T cell responses induced by both vaccination and infection remained robust and were able to eliminate Omicron infected cells ^33, 34^. This observation led to the speculation that well-preserved T cell immunity to Omicron contributes to protection from severe COVID-19 disease ^34, 35^. Interestingly, T cell immunity induced by SARS-CoV-2 vaccines was demonstrated to be highly cross-reactive against the Omicron and Delta variant ^36^, which suggests that the current vaccines may actually be providing some protection against severe disease through T cell immunogenicity, by stimulating T-cell responses against the HLA restricted T cell epitopes on the spike protein.

However, if the current vaccines do indeed mediate some of their protection by stimulating the cellular arm of the immune system, such protection is clearly limited to the HLA restricted epitopes that exist in the spike protein and it would be clearly beneficial to consider additional epitope rich regions that exist across the entire proteome of the virus. This would be critically important if we envisage designing future T-cell centric vaccines that can provide universal protection across the global spectrum of HLA haplotypes in the human population. Consequently, a better mapping of the T-cell epitope landscape across multiple different HLA’s will empower the vaccine community with the necessary knowledge and cellular immunity toolkit to guide future vaccine design and development and facilitate the development of vaccines that are better equipped to combat future VOCs.

In this study, we map a predicted T cell epitope landscape for the five main SARS-CoV-2 VOCs. Through the exploitation of state-of-the-art *in silico* methods that incorporate advanced artificial intelligence predictors of universal T cell immunogenicity, we profiled the Class I HLA immunogenicity for the five VOCs against 156 of the most frequent HLA alleles in the human population. We demonstrate conclusively in this extensive analysis that immunogenic T cell epitopes are robust across all VOC, for almost all HLA alleles in the human population, and fully protective when we incorporate the most frequent HLA haplotypes. The experimental accumulation of this evidence using *in-vitro* or *in-vivo* approaches would require extensive laborious, expensive, and time-consuming efforts. The far more comprehensive *in-silico* analysis described in this study and the subsequent community resource it generated corroborates the existing recent experimental evidence of robust T cell protection against VOCs (whereby the experimental evidence is supported by only a very limited number of HLA alleles). Taken together, the resource generated through this computational study may guide the development of T cell vaccines and diagnostics that are more robust against current and future VOCs, and their subvariants. In addition, the statistical and AI predictive metrics applied here may be deployed to quickly gauge the potential alteration of cellular immunity against future SARS-CoV-2 lineages that may characterize emerging VOCs, as we show for the Omicron-BA2 strain.

## RESULTS

### A resource of mutated epitopes in VOCs from the perspective of their antigen presentation potential

We performed a pairwise comparative analysis of the predicted antigen presentation (AP) scores between each of the VOCs and the original SARS-CoV-2 strain identified in Wuhan. The AP scores were computed using a state-of-the-art AI engine that predicts the potential of HLA-epitope complexes to be presented on the surface of host infected cells by Class I HLA alleles, and therefore recognizable by cognate T cells (see Methods). The peptides originating from the Wuhan strain are referred to here as the “*wildtype”*, while the mutated peptides emerging from any of the VOC are referred to as “*mutant”*. In this study, we limited the scope of our analysis to an examination of the differences in AP potential among either the mutated or wildtype peptides that were predicted to have a high likelihood of being bound to HLA and presented to the cell surface. Therefore, in the following analyses, the AP scores were filtered such that only peptides with an AP score > 0.5 (in a scoring system that ranges from 0 to 1) in either the mutant or the corresponding wildtype peptide were considered. Consequently, the analyses presented here only considers potential T cell epitopes that either have a high AP score in both wildtype and associated mutant peptide or were presented with a gain or loss of AP potential as an outcome of the mutation. The selection of 0.5 as an AP score threshold was motivated by i) the need for sufficiently high AP potential, such that we ignore peptides with a negligible or low chance of being presented on the cell surface, and ii) the need to have enough viable epitopes for the comparison of AP score distributions between mutant and wildtype (i.e., to ensure statistical power). However, the exact choice of threshold is not critical since we only compare differences between variants. The threshold can be tuned in an analysis, and we provide the entire landscape of AP predictions for the mutated peptides and their corresponding wildtypes across all VOCs (i.e., irrespective of their AP score) across 156 of the most frequent Class I HLA alleles in the human population in Supplementary File 1, as a resource to the community to further explore the data.

### The AP profile (a proxy of T-cell immunity) is robust across all SARS-CoV-2 VOCs

Non-synonymous mutations in the proteins of SARS-CoV-2 had almost no effect on their potential to be presented by the most frequent HLA alleles in the human population, and thereby recognized by cognate HLA restricted CD8+ T cells from a global perspective (the 156 most frequent Class I HLA alleles across all mutated peptides in all VOCs). We observed highly similar AP score distributions between the mutant and corresponding wildtype peptides. It should be noted that we were not, in this analysis, interested in the difference in AP scores between mutant and wildtype at the individual peptide level, as the loss in AP score in one peptide might be offset by gain in AP score in another peptide. Instead, we focus on the distribution of the AP scores which are illustrated in Figure 1A for each VOC. The variation in peptide counts we observed was due to a different number of mutated peptides for each VOC that pass the 0.5 AP threshold. The number of mutated peptides was simply a function of the number of mutations for a given strain (*i*.*e*., more mutations translated to more mutated peptides). When investigating these distributions pooled across all VOCs, we observed that the shapes of these distributions were very similar, as depicted in Figure 1A, subfigure titled “Comparison”. Here, the AP score distributions were layered on top of each other, using percentages for each bin instead of the raw count. The reason why no sharp cutoff can be observed around an AP of 0.5 is because a peptide was considered for analysis if it has an AP score > 0.5 in either the mutant or wildtype. Figure 1B shows the same AP score distributions with a violin plot representation to further facilitate the interpretation of these pairwise comparisons. Again, we observed that the difference in AP score distributions between mutant and wildtype was negligible.

**Figure 1:**
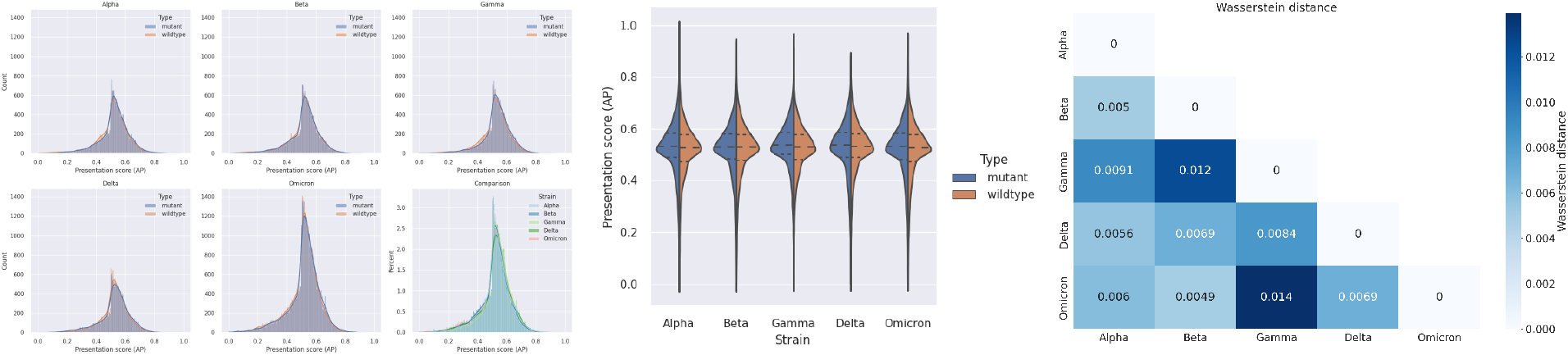
Comparisons of the distribution of antigen presentation (AP) scores for each SARS-COV-2 variant of concern (VOC) for the mutated and corresponding wildtype peptides. The AP scores have been filtered such that only peptides with AP scores > 0.5 in either the mutant or corresponding wildtype are shown. **A**: Histogram with a kernel density estimate of the distribution of AP scores for the mutant and wildtype peptides for each VOC. The subfigure “Comparison” shows the percentage of AP scores in each bin for all VOCs. **B**: Violin plots of the AP score distribution between mutant and wildtype peptides for each VOC. The dashed lines within each violin plot represent quartiles. **C**: The pairwise Wasserstein distance between the distribution of AP scores for the mutated peptides in the VOCs.

The same trend was observed when we analyzed the relatively minor shifts in AP score distributions between VOCs. This is confirmed by Figure 1C were the Wasserstein distance ^37^, also commonly called “the earth mover” distance, was calculated and plotted for the pairwise distance between each VOC. The intuitive explanation for the Wasserstein distance is how much “work” it takes to transform one distribution into another. In other words, it reflects how much of one distribution we need to move, multiplied by the distance it is moved. Note that the unit of the Wasserstein distance is the same as for the AP score. An advantage of investigating the Wasserstein distance between these distributions is that it takes the specific shape of each distribution into account. As Figure 1C shows, only small Wasserstein distances were observed between the different VOCs.

Table 1 summarizes the differences in AP score distributions between mutant and wildtype peptides within each VOC. The average difference in AP scores, column one in Table 1, is an intuitive metric to assess the impact of the collective set of mutations occurring within a VOC. It can quickly show us in a global manner if there was a general gain or drop in the AP scores in a VOC compared to the wildtype strain. However, a drawback of the mean difference, is that it only considers the mean of the distributions and does not capture the shape of the AP score distributions.

**Table 1:**
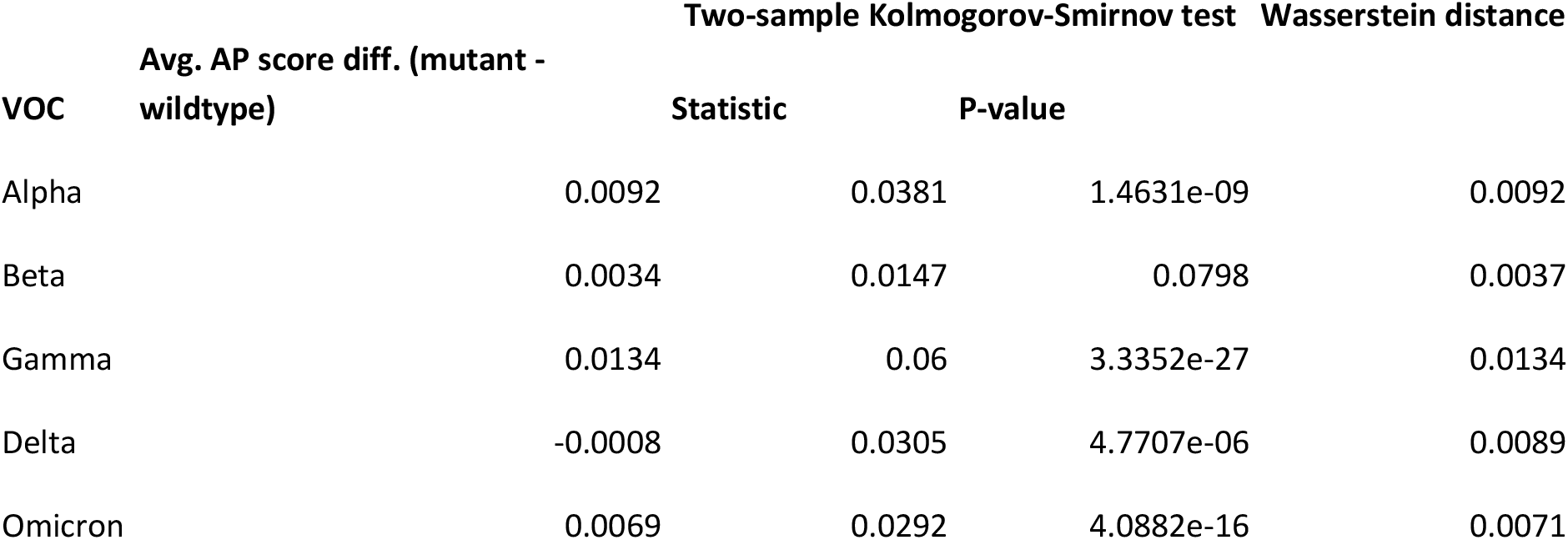
Different metrics summarizing the difference in AP scores between mutant and wildtype peptides within a VOC. Three different methods were used to compare the distributions: i) the aver-age difference in AP scores, ii) the two-sample Kolmogorov-Smirnov (KS) test, iii) and the Wasserstein distance.

| VOC | Avg. AP score diff. (mutant - wildtype) | Two-sample Kolmogorov-Smirnov test |  | Wasserstein distance |
| --- | --- | --- | --- | --- |
|  |  | Statistic | P-value |  |
| Alpha | 0.0092 | 0.0381 | 1.4631e-09 | 0.0092 |
| Beta | 0.0034 | 0.0147 | 0.0798 | 0.0037 |
| Gamma | 0.0134 | 0.06 | 3.3352e-27 | 0.0134 |
| Delta | -0.0008 | 0.0305 | 4.7707e-06 | 0.0089 |
| Omicron | 0.0069 | 0.0292 | 4.0882e-16 | 0.0071 |

The two-sided Kolmogorov-Smirnov (KS) test statistic, second column in Table 1, shows the largest absolute difference between the two empirical distribution functions, for both mutant and wildtype, and lies in the interval [0, 1]. Like the Wasserstein distance, the two-sided KS test statistic considers the shape of the distributions. However, the maximum distance is reached as soon as the two distributions no longer overlap. This is a crucial difference with respect to the Wasserstein distance, which continues to increase the further the two distributions diverge from each other. It should be noted that while the p-value for the KS test is significant (< 0.05) (i.e., the AP scores come from statistically different distributions) for all but the Beta variant, the p-value does not inform us on how different the distributions are from each other.

Overall, Table 1 shows us that the different metrics are in good agreement when considering the difference in AP score distributions between the mutant and wildtype, for each VOC. All three metrics are very low, considering that the range of AP scores and KS statistic is [0, 1]. Moreover, ranking the VOC from largest to smallest difference between mutant and wildtype based on the KS statistic or Wasserstein distance results in the same order.

However, this analysis compared only the subset of peptides containing mutations to their corresponding wildtype counterpart. As Table 2 shows, this subset of mutated peptides represents only a small fraction out of the total number of peptides in the original SARS-CoV-2 Wuhan strain. In this table, *Wildtype* refers to the number of peptides with an AP score > 0.5 in the complete, original Wuhan strain. As column three shows, the number of peptides with an AP score > 0.5 in either the mutant or its corresponding wildtype counterpart, for each of the VOC, is relatively small when compared to the number of wildtype peptides. Even for Omicron, the VOC with the highest number of mutations (59), we observe that only ∼5% of the peptides with a high AP score are mutated when compared to the Wuhan wildtype. In other words, Table 2 helps to put the results presented here in perspective, as they reflect only the small fraction of peptides that underwent nonsynonymous mutation events. Therefore, with the comparison of the AP score distributions in mind, one must consider the fact that ∼95% or more of the peptides with an AP > 0.5 in each VOC are of course identical.

**Table 2:** Number of peptides with AP score > 0.5 in either the mutant or the corresponding wildtype for each of the VOCs and the fraction they represent out of the Wuhan Wildtype. “Wuhan Wildtype” is the number of peptides with antigen presentation score > 0.5 in the complete original Wuhan strain.

| VOC | Nb. of non synonymous mutations | Nb. of mutated peptides | Fraction of all wildtype peptides in the original Wuhan strain |
| --- | --- | --- | --- |
| Alpha | 21 | 14485 | 0.0173 |
| Beta | 22 | 14890 | 0.0178 |
| Gamma | 25 | 17106 | 0.0204 |
| Delta | 22 | 13952 | 0.0167 |
| Omicron | 59 | 42356 | 0.0506 |
| Wildtype (Wuhan) | NA | 837211 | 1 |

As previously mentioned, the Supplementary File 2 (S2-Figures 2, 4, 6, and 10 and the raw data table) also offers, as a resource, the complete set of AP score predictions for all mutated peptides and all HLA alleles considered for each of the VOCs without any filtering, allowing to study the distributions of the entire set of AP scores.

**Figure 2:**
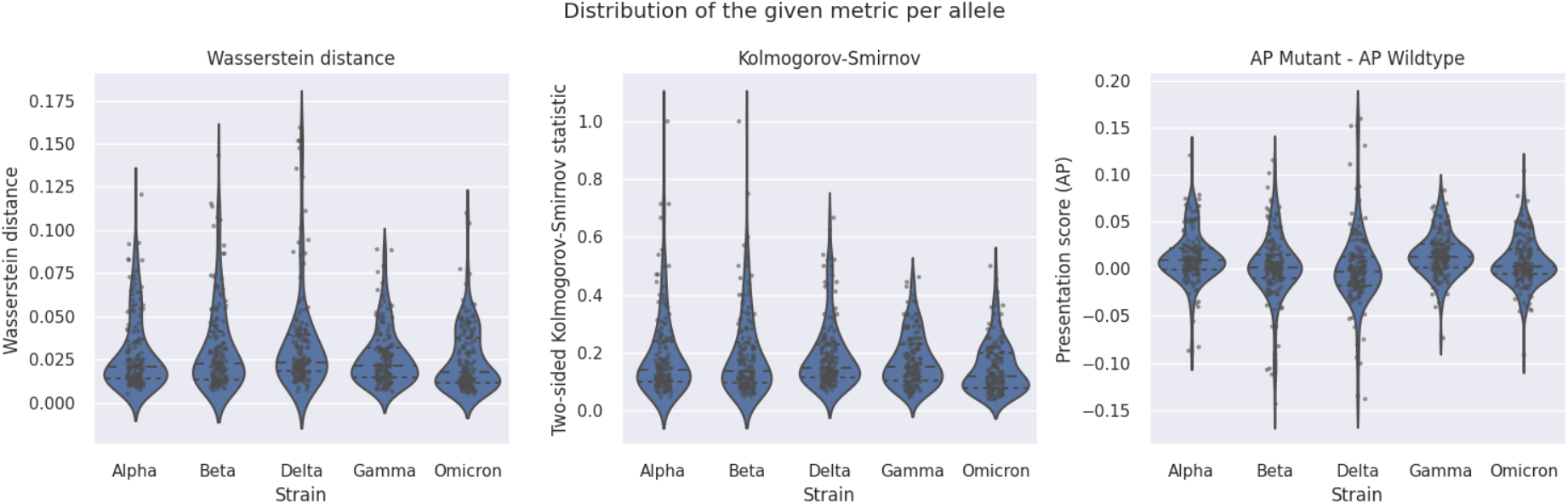
Violin plots of the distributions of the Wasserstein distance, two-sided Kolmogorov-Smirnov statistic, and average difference in AP score between mutant and wildtype epitopes across the HLA alleles, for each VOC. The dashed lines within each vioslin represent the quartiles and the dots show the score for each metric for each epitope.

We also examined the recently emerged more transmissible Omicron sub-lineage BA.2 as a use case, the plots including this variant can be found in Supplementary File 2 (S2-Figures 1, 3, 5, 7, and 9). In brief Omicron BA.2 diverged slightly further from the wildtype than Omicron and presents an increase in average difference of the AP score when compared to Omicron, suggesting that it may be more immunogenic from the perspective of T-cell immunity. However, the differences remain small. Overall, the biggest difference in AP scores due to mutations was found in Gamma (Wasserstein: 0.0134, avg. difference in AP score: 0.0069) and Omicron-BA.2 (Wasserstein: 0.0108, avg. difference in AP score: 0.0108).

**Figure 3:**
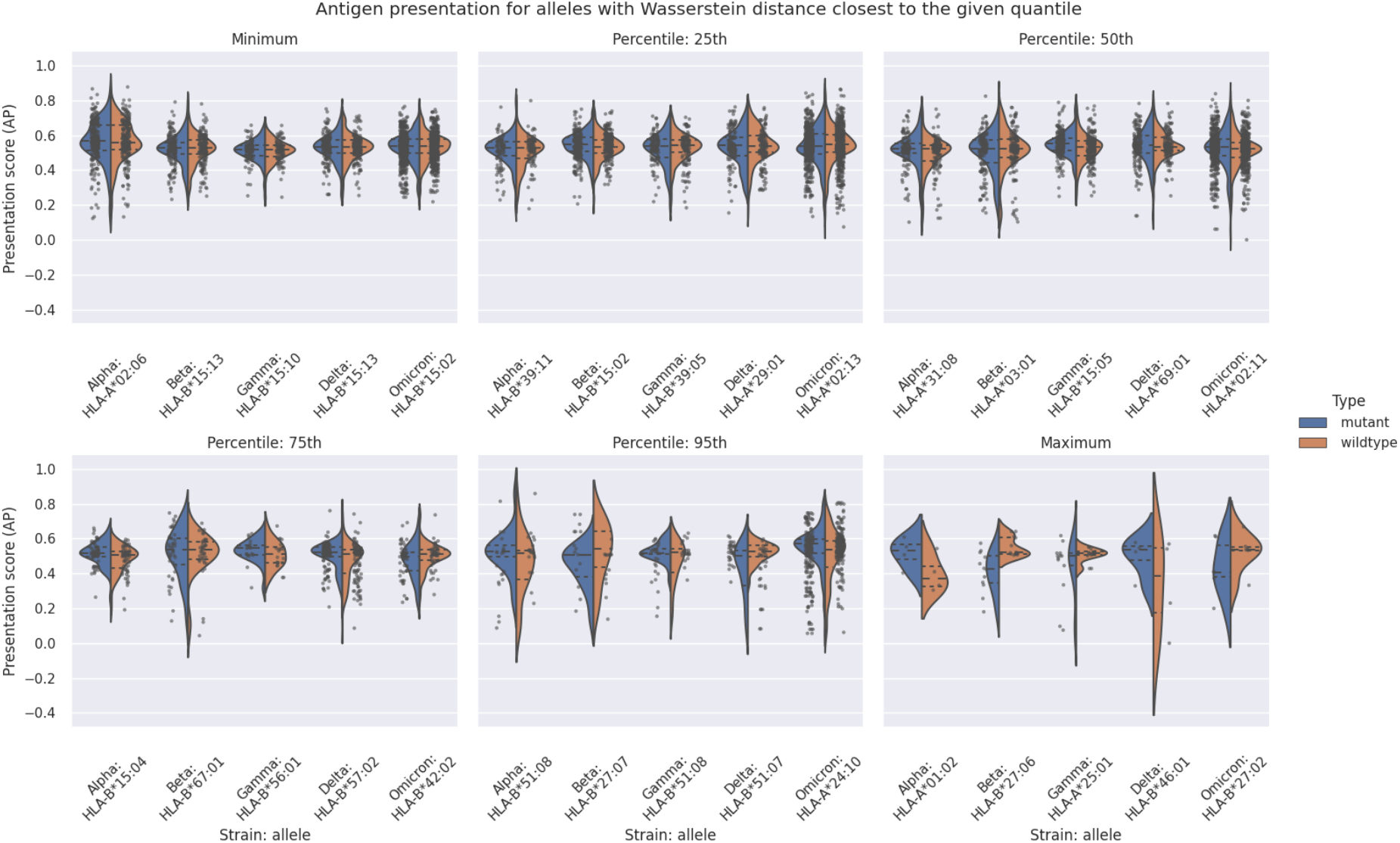
Violin plots of the AP score distribution for the mutant and wildtype peptides for the HLA alleles with a Wasserstein distance closest to the minimum, maximum, or given percentile of all the Wasserstein distances for that VOC. The dashed lines within each violin represent the quartiles and the dots show the AP for each peptide.

### The antigen presentation potential across different VOCs does not significantly differ for most of the HLA alleles considered in the analysis

We next examined the distribution of AP scores for the mutant and the corresponding wildtype peptides on an HLA allele specific basis, to assess if there are specific HLA alleles in the human population that are more susceptible to immune escape by the emerging VOCs. Table 3 shows the mean and the 5^th^ percentile (for the average AP score difference only), in addition to 25^th^, 75^th^, and 95^th^ percentiles of the Wasserstein distance, two-sided KS test statistic, and of the average AP score difference between mutant and wildtype epitopes for each VOC. The mean and percentiles were calculated based on the set of individual values for each metric that were inferred for each of the available HLA alleles per VOC. Since the Wasserstein distance and the two-sided KS test statistic have a minimum value of 0, we are only concerned with the larger percentiles. For the average AP score difference on the other hand, which can range from -1 to 1, we looked at the mean, 25^th^ and 75^th^ percentiles, in addition to the 5^th^ and 95^th^ percentiles to capture the outliers.

**Table 3:** Mean, 5th (for the average AP score difference only), 25th 75th, and 95th percentiles of the Wasserstein distance, of the two-sided Kolmogorov-Smirnov statistic, and of the average difference in AP score between mutant and wildtype epitopes across the HLA alleles for each VOC.

| VOC | Avg. AP score diff. (mutant - wildtype) |  |  |  |  | Two-sample Kolmogorov-Smirnov test statistic |  |  |  | Wasserstein distance |  |  |  |
| --- | --- | --- | --- | --- | --- | --- | --- | --- | --- | --- | --- | --- | --- |
|  | Percentile |  |  |  |  | Mean | Percentile |  |  | Mean | Percentile |  |  |
|  | Mean | 5th | 25th | 75th | 95th |  | 25th | 75th | 95th |  | 25th | 75th | 95th |
| Alpha | 0.0127 | -0.0224 | -0.0006 | 0.0217 | 0.065 | 0.1898 | 0.1011 | 0.2391 | 0.4779 | 0.0285 | 0.0138 | 0.0355 | 0.0729 |
| Beta | 0.0028 | -0.0478 | -0.0094 | 0.0152 | 0.0596 | 0.1935 | 0.0956 | 0.2362 | 0.475 | 0.0316 | 0.0136 | 0.0412 | 0.0863 |
| Gamma | 0.0151 | -0.0522 | 0.0012 | 0.0266 | 0.0622 | 0.1738 | 0.1042 | 0.2301 | 0.475 | 0.0268 | 0.0148 | 0.0318 | 0.0959 |
| Delta | 0.0006 | -0.017 | -0.018 | 0.0129 | 0.0564 | 0.1919 | 0.114 | 0.2287 | 0.3333 | 0.0343 | 0.0181 | 0.0393 | 0.0595 |
| Omicron | 0.0074 | -0.0302 | -0.0054 | 0.0204 | 0.05 | 0.1473 | 0.0781 | 0.2 | 0.3333 | 0.0251 | 0.0115 | 0.0377 | 0.0556 |

From Table 3, we observed that the mean of the average difference in AP score and the mean of the Wasserstein distance are both very small. The mean of the two-sided KS test statistic is slightly larger, as expected, but can still be considered small in this context. The absolute values of the 25^th^ and 75^th^ percentiles of the average difference in AP scores were also small, therefore we examined the 5^th^ and 95^th^ percentiles to see the larger differences. However, no extreme shifts in the average difference in AP scores were observed. For the Wasserstein distance, we observed a similar distribution at the 75^th^ percentile whereby the differences in distributions between the mutant and wildtype were relatively small, while marginally larger difference was first observed only at the 95^th^ percentile.

In Figure 2 we plotted the distributions of these three metrics across the entire set of HLA alleles for each VOC. We observed that out of the 156 HLA alleles, the vast majority do not show substantial differences between the two AP score distributions, as also shown in Table 3. This suggests that, at a first glance, there is no apparent HLA (sub)population at more risk than other HLA (sub)populations due to lowered T-cell responses to a given VOC. Plots of the AP score distribution for each allele, can be found in Supplementary File 2 (S2-Figure 5).

Figure 3 shows the distribution of AP scores for mutant and wildtype epitopes for a selected group of HLA alleles. Each plot in the figure corresponds to either the minimum, maximum, or a specific percentile. For each VOC, we selected HLA alleles such that their Wasserstein distance, between mutant and wildtype is the closest to the minimum, maximum, or the given percentile. For instance, for “Percentile: 25^th^”, the HLA alleles shown are the ones for which the Wasserstein distance is the closest to where 25% of the data lives with respect to the Wasserstein distance metric. This analysis serves as a guide to establish a connection between the AP score distributions and the Wasserstein distance by showing representative samples for the AP score distributions between mutant and wildtype epitopes. A more detailed figure with all HLA alleles can be found in Supplementary File 2(S2-Figure 3).

From Figure 3 we observed that up to and including 50th percentile. (i.e., the median) the differences in AP score distribution between mutant and wildtype are relatively small. Even at 75^th^ percentile (i.e., third quartile) the AP score distributions are very similar.

As such, the consensus conclusion from Table 2, Figure 2, and Figure 3 is that only ∼5% of the 156 HLA alleles analyzed demonstrated larger differences in AP score distributions between mutant and wildtype. Consequently, most HLA alleles in the human population do not demonstrate an altered propensity to present mutated SARS-CoV-2 peptides across the different VOCs.

Even if the vast majority of HLAs present a similar AP score distribution between mutant and wildtype, as Figure 3 shows, a very small subset of HLA alleles presented a considerable gain or loss in AP score between mutant and wildtype (see Figure 3, “Percentile: 95th or “Maximum”). These correspond to the HLA alleles with the largest Wasserstein distance between mutant and wildtype, for each VOC. Based on these findings, we then assessed which haplotypes are potentially the most affected. We took the three HLA alleles per VOC that showed the greatest Wasserstein distance in AP score distributions between mutant and wildtype and queried them against The Allele Frequency Net Database (AFND) ^38^. This analysis did not highlight any specific ethnicity or region of the world to be systematically affected by a given VOC, although the Australian population seems more affected by three of the VOCs as Table 4 shows.

**Table 4:**
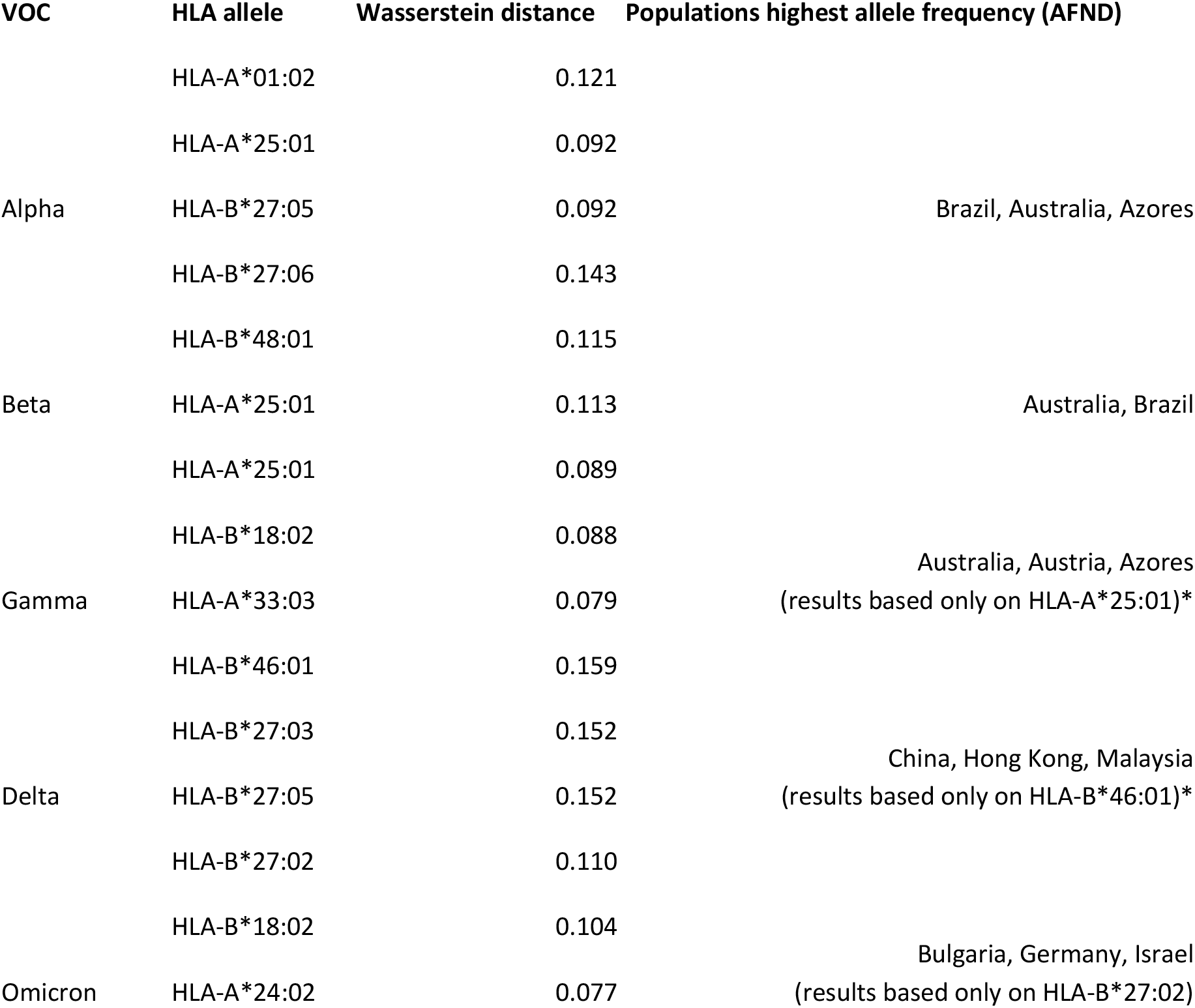
Top three HLA alleles that present the biggest Wasserstein distance for each of the VOC and their corresponding haplotype according to The Allele Frequency Net Database (AFND). The last column shows in which populations the set of HLA alleles is presented with the highest frequency. For the cells denoted by an * it means that the AFND database only provided results for one of the top three most affected HLA alleles.

Another interesting observation, summarized in Table 5, was that the HLA-A alleles often had a Wasserstein distance that was an order of magnitude higher than the HLA-B alleles. Additionally, the HLA-A alleles have a higher average AP score difference than the HLA-B alleles, for all VOCs but Beta.

**Table 5.**
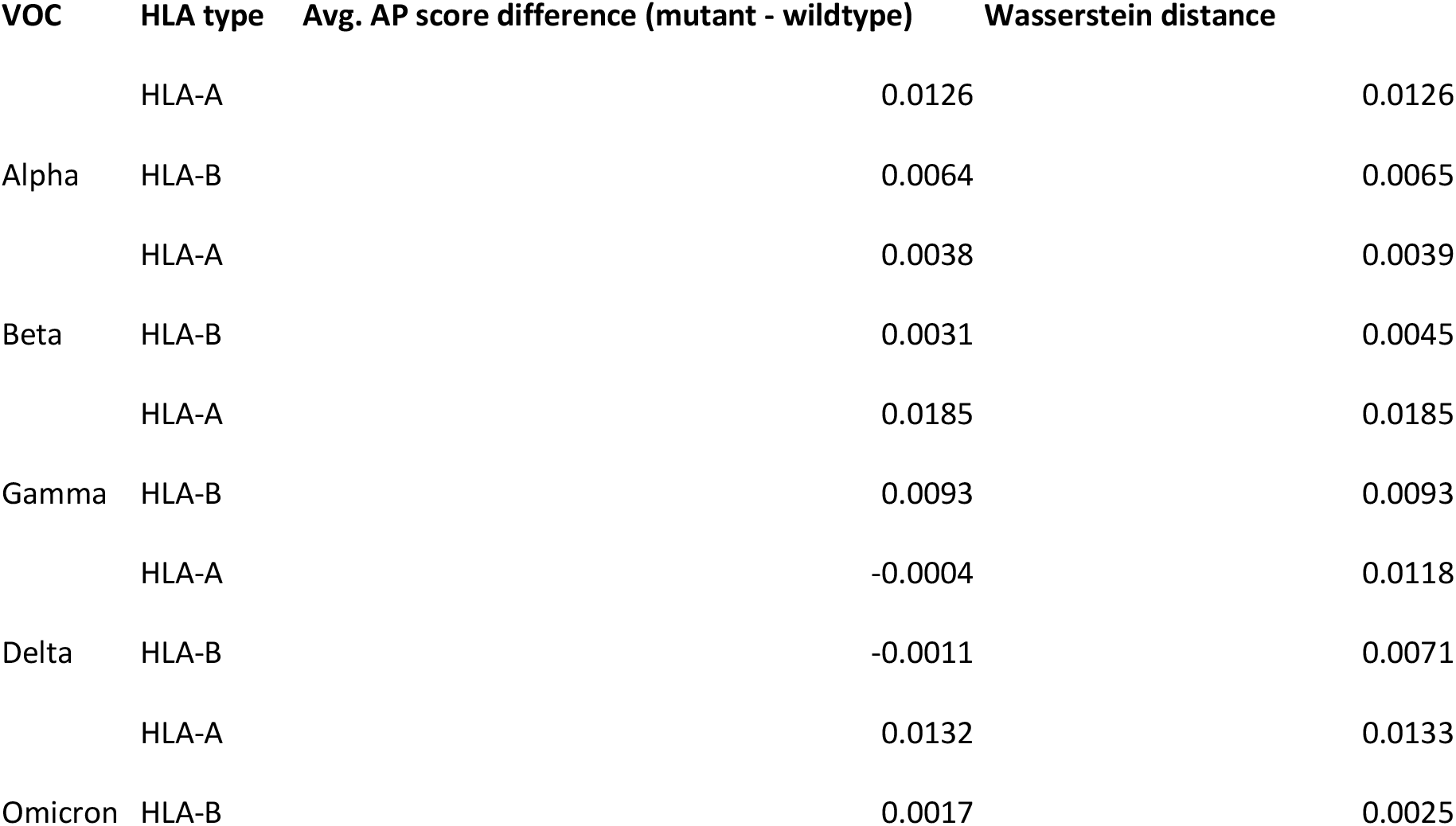
Wasserstein distance and average difference between the distribution of AP scores for the mutant and corresponding wildtype peptides for all HLA-A and HLA-B alleles.

| VOC | HLA type | Avg. AP score difference (mutant - wildtype) | Wasserstein distance |
| --- | --- | --- | --- |
| Alpha | HLA-A | 0.0126 | 0.0126 |
|  | HLA-B | 0.0064 | 0.0065 |
| Beta | HLA-A | 0.0038 | 0.0039 |
|  | HLA-B | 0.0031 | 0.0045 |
| Gamma | HLA-A | 0.0185 | 0.0185 |
|  | HLA-B | 0.0093 | 0.0093 |
| Delta | HLA-A | -0.0004 | 0.0118 |
|  | HLA-B | -0.0011 | 0.0071 |
| Omicron | HLA-A | 0.0132 | 0.0133 |
|  | HLA-B | 0.0017 | 0.0025 |

### A mutation centric perspective of the T cell epitope landscape of VOCs

To assess the effect of a specific non-synonymous mutation on the potential of the subsequent mutated peptides to be presented as an HLA bound epitope to T-cells, we examined the average difference in AP score between mutant and wildtype epitopes for each nonsynonymous mutation in each VOC. The results are illustrated in Figure 4, stratified by each of the different SARS-CoV-2 proteins. We chose the average difference in AP score since we were specifically interested in the direction of the change around each mutated peptide. A positive value means that the AP score on average is increased for a given mutation, while a negative score means that the AP score decreases for that mutation in the variant. From Figure 4, we see that while most mutations do not seem to have a substantial impact with respect to the AP score, there are still a few outliers.

**Figure 4:**
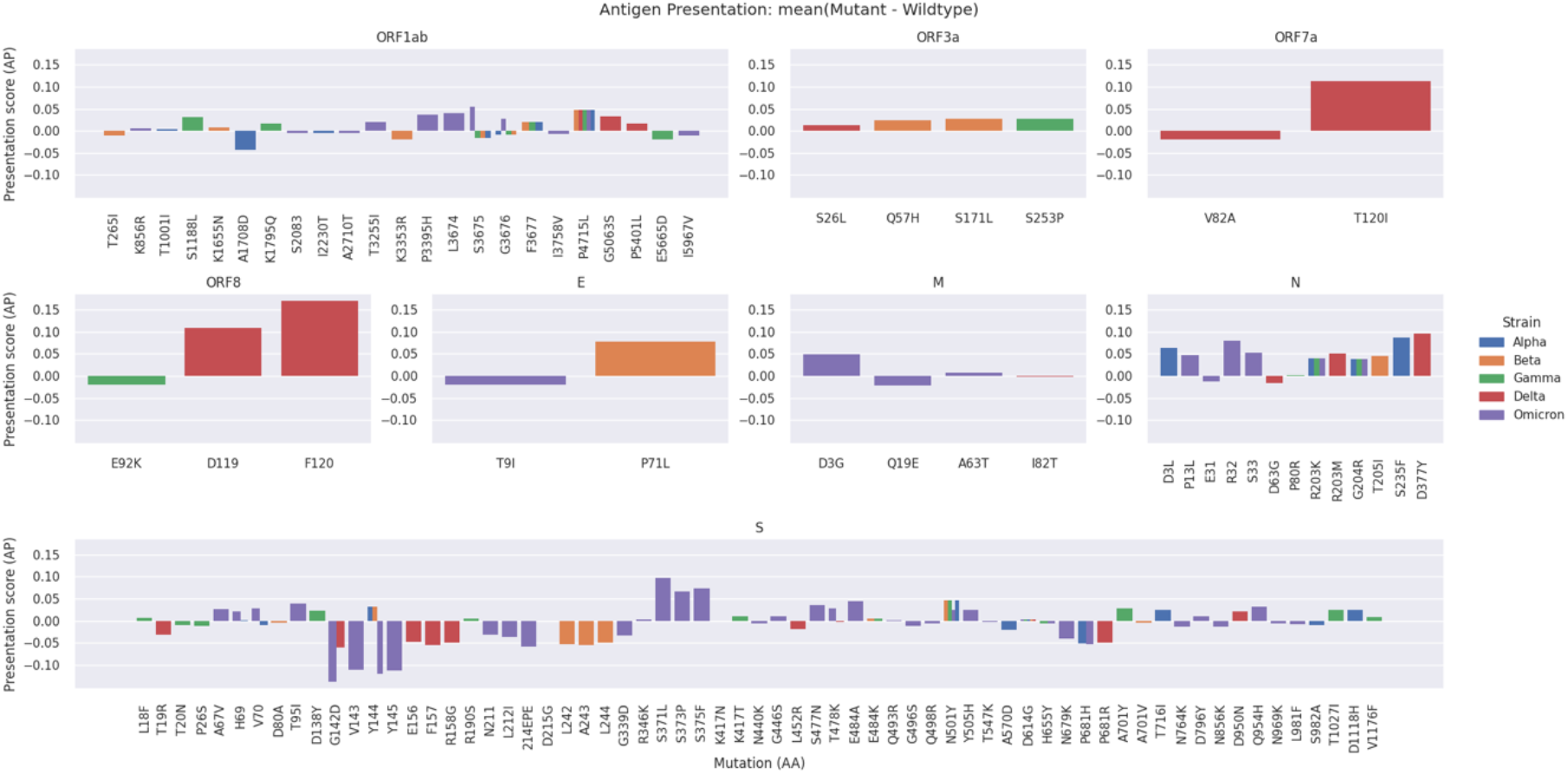
Average difference in AP score (mutant – wildtype) for each mutation in the VOC, denoted by the different colors. Each plot corresponds to a protein of the virus in which mutations occurred.

Distinct set of mutations that were found to have the largest impact in AP scores, were generally co-occurring on the same candidate epitope. As depicted in Figure 5, co-occurring mutations, for example S: S371P, S: S373F, and S: K375N in the Omicron VOC, were more likely to increase the difference in AP potential of the candidate epitopes where they lie. In turn, this makes these specific amino acid variations more impactful in in terms of resulting T cell immunity. In this specific case, there was a gain in the AP scores for all three of the co-occurring mutations.

**Figure 5:**
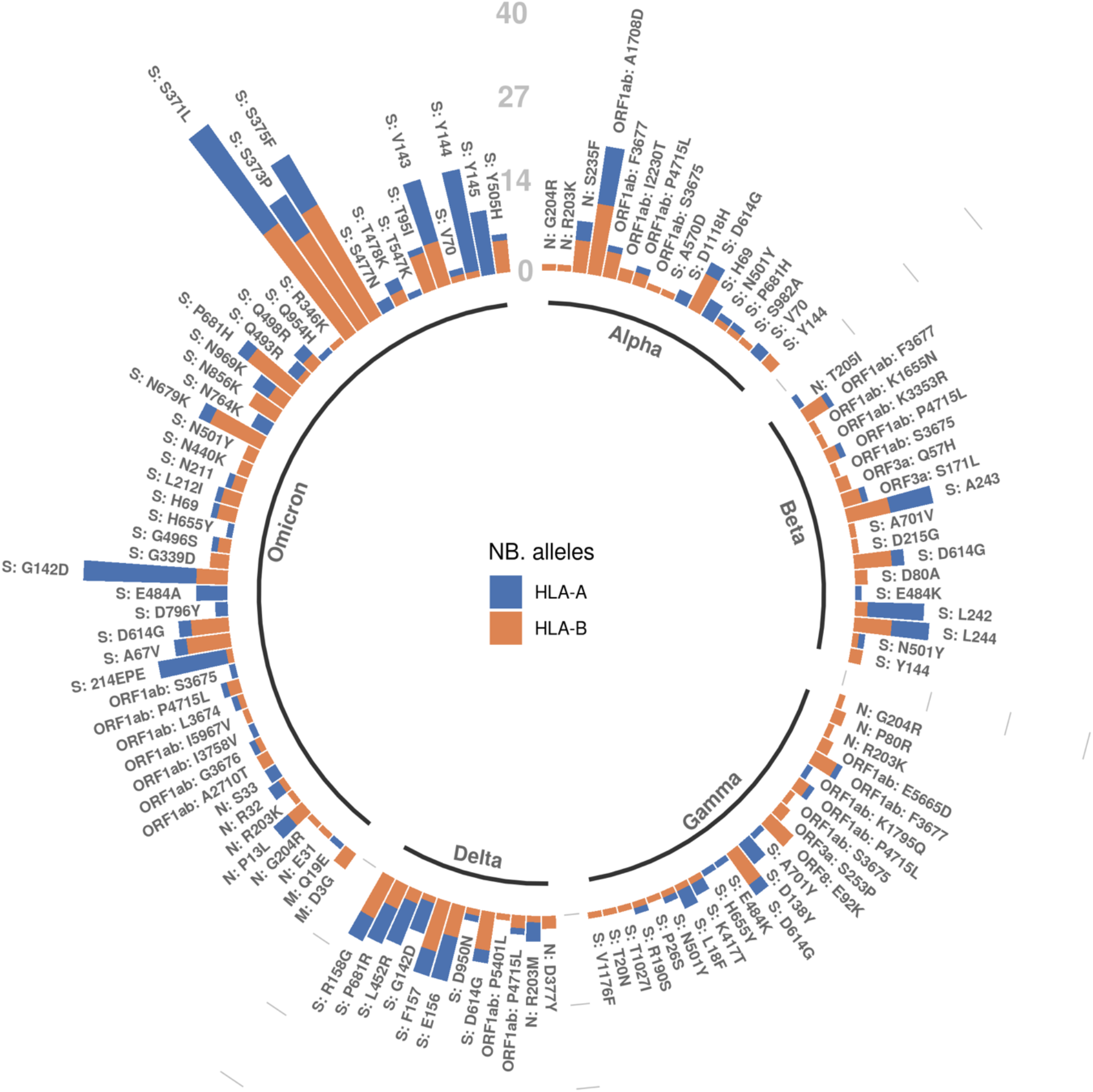
The number of HLA alleles per mutation for which there was a significant difference (p-value < 0.05) using the two-sided Kolmogorov-Smirnov test on the mutant and the corresponding wildtype AP score distribution across VOCs. The label of each bar contains the protein name and the mutation details separated by colon. The number of HLA-A alleles and HLA-B alleles is displayed separately for each mutation.

Figure 5 summarizes the number of HLA alleles per mutation that present a significantly different AP score (p-value < 0.05 for the two-sided KS test) as compared to the corresponding wildtype peptides, we also provide in Supplementary File 2 similar plots with a summary of the number of peptides with an AP score > 0.5 in either the mutant or wildtype which served as input for the AP score difference significance test (Supplementary File 2, S2-Figure 9). Notably, as also observed in Table 5, there seems to be a general trend for HLA-B alleles to be more affected than HLA-A alleles with respect to the AP score.

## METHODS

### Shared mutation profile of VOCs

The list of mutations for each VOCs, as labeled by the World Health Organization (WHO), was compiled from https://www.who.int/en/activities/tracking-SARS-CoV-2-variants. For each mutation, all overlapping peptides of length 9 and 10 that overlap a given mutation were considered as candidate HLA restricted epitopes.

### Predicted probability of SARS-CoV-2 mutated peptides being HLA-presented on the surface of host infected cells

The important determinants of antigen presentation (AP) were assessed for each mutated peptide and their wildtype counterpart in the VOCs for their potential to be efficiently presented. These determinants consisted of; (1) the predicted binding affinity between the candidate peptide and 156 of the most frequent HLA molecules in the human population, (2) the predicted potential of the candidate peptide to be efficiently processed by the antigen processing machinery of the host infected cell and (3) the predicted probability of the candidate mutated peptide to be presented on the host infected cell surface, which, among other factors, takes binding and processing into account. The NEC Immune Profiler (NIP) was used as a source to provide the AI predictions for these key determinants, such as the AP scores ^39^. The AP score is in the range between 0 and 1, with 1 being the maximum of the likelihood that a specific candidate mutated peptide was presented on the host infected cell surface.

The AP scores were calculated based on a set of 156 most frequent Class I HLA alleles (A and B) for all the possible combinations between an HLA allele and each epitope overlapping a given mutation, for each VOC. The complete set of raw AP score predictions is provided in the raw data table in the Supplementary File 2.

### Statistical analysis

The statistical analyses were performed in the Python programming language (version 3.6.12) using the *SciPy* (version 1.3.1) package which implements functions for the Kolmogorov-Smirnov test and Wasserstein distance. For validation and complementarity purposes, the R programming language (version 4.1.1) was also used to generate a separate set of statistics on the same input data. Package *stats* in base R was used for the Kolmogorov-Smirnov test and package *transport* (version 0.12-2) to calculate the Wasserstein distance.

## DISCUSSION

The health and socio-economic burden of current and future potential emerging SARS-CoV-2 VOCs, remains entirely unknown. The most recent Omicron VOC that emerged, fortunately produced a milder COVID-19 disease, especially in highly vaccinated or infected populations ^7^. The milder manifestation of the disease was in the backdrop of far higher transmissibility of Omicron compared to Delta, and indeed all other VOCs. However, Omicron and the recently emerged Omircron-BA2 subvariant, continues to circulate with hyper-transmissible rates ^27^ accompanied by high immune escape, as measured through antibody serological responses ^40^.

The continuous rapid circulation of Omicron places critical importance in predicting the various clinical, epidemiological, and immune-escape parameters of VOCs in the human population, as Omicron and its subvariants transitions toward an endemic threat ^41^. However, as SARS-CoV-2 progresses on its path toward becoming endemic ^42^, this transition may not necessarily equate to less virulent or milder disease ^4^. There is strong reasoning that Omicron’s manifestation as a milder COVID-19 disease, in the backdrop of its rapid antigenic evolution, may be a coincidence, and more virulent and dangerous escape variants may emerge in the future ^43^. The distinct antibody immune escape observed during the COVID-19 pandemic was not only limited to Omicron ^3^, as several other VOCs also demonstrated a reduced susceptibility to being neutralized by vaccine induced antibody ^44^. Accordingly, as the human population acquires increasing levels of immunity through vaccination and/or infection; the evolutionary trajectory of SARS-CoV-2 toward increasing infectivity (through optimized host receptor binding), alone, will not satisfy its natural evolutionary drive toward increasing transmission rates. To combat the increasing immunity in the human population, SARS-CoV-2 is predicted to also evolve by escaping natural or vaccine induced immunity and gain the ability to infect previously protected individuals ^43^. Thus, as SARS-CoV-2’s future evolution drives ever increasing levels of antigenic drift it is important that we characterize the correlates of protection and potential immune escape for both B cell and T cell human immunity, to guide future vaccine and diagnostic design.

As discussed in the introduction to this study, the T cell immunity that correlates with protection against SARS-CoV-2 has been shown to be more durable and cross protective ^45^ compared to the relatively transient and narrow (strain specific), protection afforded by antibody responses as witnessed by the recent Omicron VOC ^21^. Although there have been some efforts to profile T cell responses against SARS-CoV-2 VOCs in vitro with a limited number of HLA alleles ^11, 46^; an all-encompassing screen that covers the global human population would be far too time-consuming, laborious, and expensive, to generate using wet lab approaches, necessitating the need for AI-based *in silico* profiling approaches.

Here we provide an AI generated resource of mutated epitopes from all current VOCs, as well as the Omicron BA2 subvariant, from the perspective of their antigen presentation potential, which can be used as a proxy for immunogenicity. The insights gained from analyzing this resource mirrors the early empirical findings from wet-lab-based studies that demonstrated that the T cell responses induced by vaccination and natural infection (from other VOC) remain cross reactive against the Omicron VOC. The *in-silico* evidence we present here highlights this trend more comprehensively, across all possible mutated epitopes in all VOCs, and across the most frequent HLA alleles in the human population.

This finding not only informs how we can track the correlates of immune protection across different vaccines and vaccine modalities, but also advocates the advancement of T cell-centric vaccine approaches to combat emerging VOCs ^47^. Relying on the current spike protein vaccines for broad protection against VOCs is arguably not viable long-term ; not only because the T cell epitope cargo is limited to a single viral protein that is subject to a high mutational rate, but also because it has been clearly demonstrated that T cell responses to the spike protein were reduced by approximately 50%, in 20% of naturally infected or vaccinated individuals ^43^.

This resource and analysis provided us the opportunity to automatically assess the difference in immunogenic potential in candidate epitopes between a VOC lineage a mutant strain and the originating SARS-CoV-2 lineage identified in Wuhan by examining the potential T cell antigen drift of the current and emerging VOCs. We demonstrated the robustness of the majority of identified T-cell epitopes across all VOCs using several distance metrics, however the Wasserstein distance and the average difference in AP score were the most informative.

Using both the Wasserstein distance and the average difference in AP score, we were able to examine different properties of the VOC AP score distributions. The average difference in AP score was easy to interpret and gave us a direction for the potential antigenic drift, however it did not consider the shape of the two AP score distributions. The Wasserstein distance did not give a direction for the antigenic drift however it considered the shape of the two distributions aiding their comparison. Based on the observations presented, one can argue that the Wasserstein distance and the average difference in AP score are very similar, so it might be sufficient to just examine the average difference in AP score. However, we still recommend examining both to cover a wider array of patterns in the distributions. Moreover, it seemed like no specific HLA populations are more affected than others, and generally the differences between mutant and wildtype per allele were very small.

The results in this study, demonstrate comprehensively, that significant antigenic drift resulting in escape against pre-existing (natural or vaccine induced) T-cell response is unlikely to emerge in the SARS-CoV-2 emerging VOCs. The analysis we outlined here, such as delta calculations of AP potential between a VOC mutated epitope and its corresponding wildtype, serves as an index of T cell antigenic drift or T-cell immunogenicity for emerging lineages of SARS-CoV-2.

## REFERENCES

1. Organization, W.H. WHO Director-General’s opening remarks at the media briefing on COVID-19 - 11 March 2020.

2. Callaway, E. Beyond Omicron: what’s next for COVID’s viral evolution. Nature 600, 204–207 (2021).

3. Dejnirattisai, W. et al. SARS-CoV-2 Omicron-B.1.1.529 leads to widespread escape from neutralizing antibody responses. Cell 185, 467–484 e415 (2022).

4. Katzourakis, A. COVID-19: endemic doesn’t mean harmless. Nature 601, 485 (2022).

5. Morens, D.M., Taubenberger, J.K. & Fauci, A.S. Universal Coronavirus Vaccines - An Urgent Need. N Engl J Med 386, 297–299 (2022).

6. Tarke, A. et al. Comprehensive analysis of T cell immunodominance and immunoprevalence of SARS-CoV-2 epitopes in COVID-19 cases. Cell Rep Med 2, 100204 (2021).

7. Sigal, A. Milder disease with Omicron: is it the virus or the pre-existing immunity? Nat Rev Immunol 22, 69–71 (2022).

8. Nelde, A. et al. SARS-CoV-2-derived peptides define heterologous and COVID-19-induced T cell recognition. Nat Immunol 22, 74–85 (2021).

9. da Silva Antunes, R. et al. Differential T-Cell Reactivity to Endemic Coronaviruses and SARS-CoV-2 in Community and Health Care Workers. J Infect Dis 224, 70–80 (2021).

10. Moss, P. The T cell immune response against SARS-CoV-2. Nat Immunol 23, 186–193 (2022).

11. Grifoni, A. et al. SARS-CoV-2 human T cell epitopes: Adaptive immune response against COVID-19. Cell Host Microbe 29, 1076–1092 (2021).

12. Su, Y. et al. Multi-Omics Resolves a Sharp Disease-State Shift between Mild and Moderate COVID-19. Cell 183, 1479–1495 e1420 (2020).

13. Stoddard, C.I. et al. Epitope profiling reveals binding signatures of SARS-CoV-2 immune response in natural infection and cross-reactivity with endemic human CoVs. Cell Rep 35, 109164 (2021).

14. Sette, A. & Crotty, S. Pre-existing immunity to SARS-CoV-2: the knowns and unknowns. Nat Rev Immunol 20, 457–458 (2020).

15. Ogbe, A. et al. T cell assays differentiate clinical and subclinical SARS-CoV-2 infections from cross-reactive antiviral responses. Nat Commun 12, 2055 (2021).

16. Lineburg, K.E. et al. CD8(+) T cells specific for an immunodominant SARS-CoV-2 nucleocapsid epitope cross-react with selective seasonal coronaviruses. Immunity 54, 1055–1065 e1055 (2021).

17. Zhao, J. et al. Recovery from the Middle East respiratory syndrome is associated with antibody and T-cell responses. Sci Immunol 2 (2017).

18. Edridge, A.W.D. et al. Seasonal coronavirus protective immunity is short-lasting. Nat Med 26, 1691–1693 (2020).

19. Tang, F. et al. Lack of peripheral memory B cell responses in recovered patients with severe acute respiratory syndrome: a six-year follow-up study. J Immunol 186, 7264–7268 (2011).

20. Wu, L.P. et al. Duration of antibody responses after severe acute respiratory syndrome. Emerg Infect Dis 13, 1562–1564 (2007).

21. Le Bert, N. et al. SARS-CoV-2-specific T cell immunity in cases of COVID-19 and SARS, and uninfected controls. Nature 584, 457–462 (2020).

22. Tregoning, J.S., Flight, K.E., Higham, S.L., Wang, Z. & Pierce, B.F. Progress of the COVID-19 vaccine effort: viruses, vaccines and variants versus efficacy, effectiveness and escape. Nat Rev Immunol 21, 626–636 (2021).

23. Oberhardt, V. et al. Rapid and stable mobilization of CD8(+) T cells by SARS-CoV-2 mRNA vaccine. Nature 597, 268–273 (2021).

24. Skelly, D.T. et al. Two doses of SARS-CoV-2 vaccination induce robust immune responses to emerging SARS-CoV-2 variants of concern. Nat Commun 12, 5061 (2021).

25. Hay, J.A. et al. Viral dynamics and duration of PCR positivity of the SARS-CoV-2 Omicron variant. medRxiv (2022).

26. Puhach, O. et al. Infectious viral load in unvaccinated and vaccinated patients infected with SARS-CoV-2 WT, Delta and Omicron. medRxiv (2022).

27. Kozlov, M. How does Omicron spread so fast? A high viral load isn’t the answer. Nature (2022).

28. Carreno, J.M. et al. Activity of convalescent and vaccine serum against SARS-CoV-2 Omicron. Nature 602, 682–688 (2022).

29. VanBlargan, L.A. et al. An infectious SARS-CoV-2 B.1.1.529 Omicron virus escapes neutralization by therapeutic monoclonal antibodies. Nat Med (2022).

30. Cele, S. et al. Omicron extensively but incompletely escapes Pfizer BNT162b2 neutralization. Nature 602, 654–656 (2022).

31. Collie, S., Champion, J., Moultrie, H., Bekker, L.G. & Gray, G. Effectiveness of BNT162b2 Vaccine against Omicron Variant in South Africa. N Engl J Med 386, 494–496 (2022).

32. Planas, D. et al. Considerable escape of SARS-CoV-2 Omicron to antibody neutralization. Nature 602, 671–675 (2022).

33. Ahmed, S.F., Quadeer, A.A. & McKay, M.R. SARS-CoV-2 T Cell Responses Elicited by COVID-19 Vaccines or Infection Are Expected to Remain Robust against Omicron. Viruses 14 (2022).

34. Keeton, R. et al. T cell responses to SARS-CoV-2 spike cross-recognize Omicron. Nature (2022).

35. GeurtsvanKessel, C.H. et al. Divergent SARS CoV-2 Omicron-reactive T-and B cell responses in COVID-19 vaccine recipients. Sci Immunol, eabo2202 (2022).

36. Liu, J. et al. Vaccines Elicit Highly Cross-Reactive Cellular Immunity to the SARS-CoV-2 Omicron Variant. medRxiv (2022).

37. Vaserstein, L. Markov Processes over Denumerable Products of Spaces, Describing Large Systems of Automata, vol. 5. Problems Inform. Transmission, 1969.

38. Gonzalez-Galarza, F.F. et al. Allele frequency net database (AFND) 2020 update: gold-standard data classification, open access genotype data and new query tools. Nucleic Acids Res 48, D783–D788 (2020).

39. Malone, B. et al. Artificial intelligence predicts the immunogenic landscape of SARS-CoV-2 leading to universal blueprints for vaccine designs. Sci Rep 10, 22375 (2020).

40. Wilks, S.H. et al. Mapping SARS-CoV-2 antigenic relationships and serological responses. bioRxiv (2022).

41. Lavine, J.S., Bjornstad, O.N. & Antia, R. Immunological characteristics govern the transition of COVID-19 to endemicity. Science 371, 741–745 (2021).

42. Shaman, J. & Galanti, M. Will SARS-CoV-2 become endemic? Science 370, 527–529 (2020).

43. Markov, P.V., Katzourakis, A. & Stilianakis, N.I. Antigenic evolution will lead to new SARS-CoV-2 variants with unpredictable severity. Nat Rev Microbiol (2022).

44. Harvey, W.T. et al. SARS-CoV-2 variants, spike mutations and immune escape. Nat Rev Microbiol 19, 409–424 (2021).

45. Tarke, A. et al. SARS-CoV-2 vaccination induces immunological T cell memory able to cross-recognize variants from Alpha to Omicron. Cell 185, 847–859 e811 (2022).

46. Naranbhai, V. et al. T cell reactivity to the SARS-CoV-2 Omicron variant is preserved in most but not all individuals. Cell 185, 1041–1051 e1046 (2022).

47. Dolgin, E. T-cell vaccines could top up immunity to COVID, as variants loom large. Nat Biotechnol 40, 3–4 (2022).

