## Supplementary File 2 for "The T cell epitope landscape of SARS-CoV-2 variants of concern"

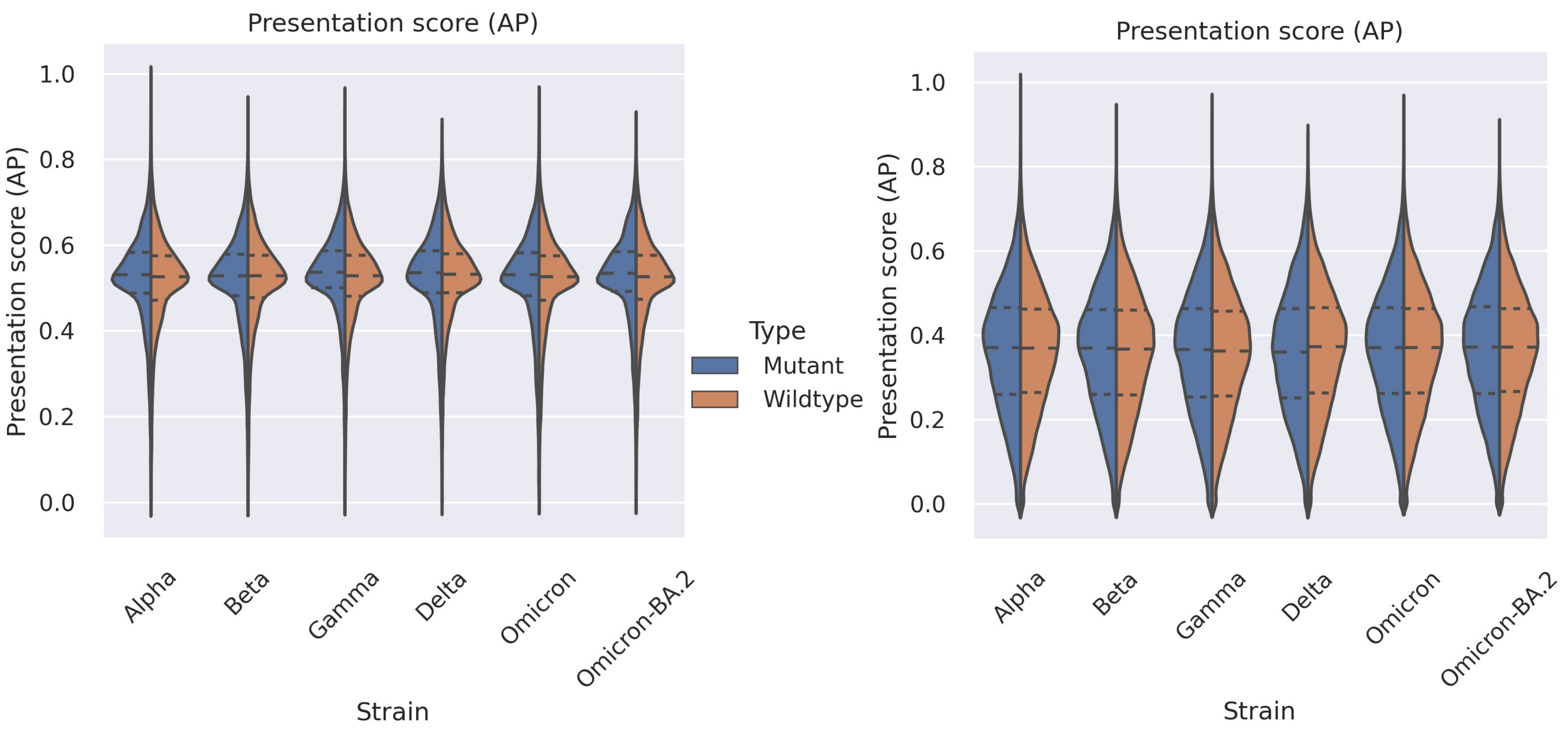


**Supplementary Figures 1-2:** Violin plots of the distribution of antigen presentation (AP) scores for the SARS-CoV2 variants (mutant) and the original Wuhan lineage (wildtype) epitopes. The Omicron BA.2 subvariant was also included along with the variants of concern. The dashed lines within each violin represent the quartiles. For the left plots, only entries with an AP score above 0.5 in either the mutant or wildtype were considered, while for the right plots, all AP score entries (i.e., no filtering) were considered.

Supplementary Figure 3: Violin plots of the distribution of presentation scores (AP) for mutant and wildtype epitopes per allele for each VOC and Omicron-BA.2. The dashed lines within each violin represent the quartiles. Only entries with an AP score above 0.5 in either the mutant or wildtype were considered for this plot.

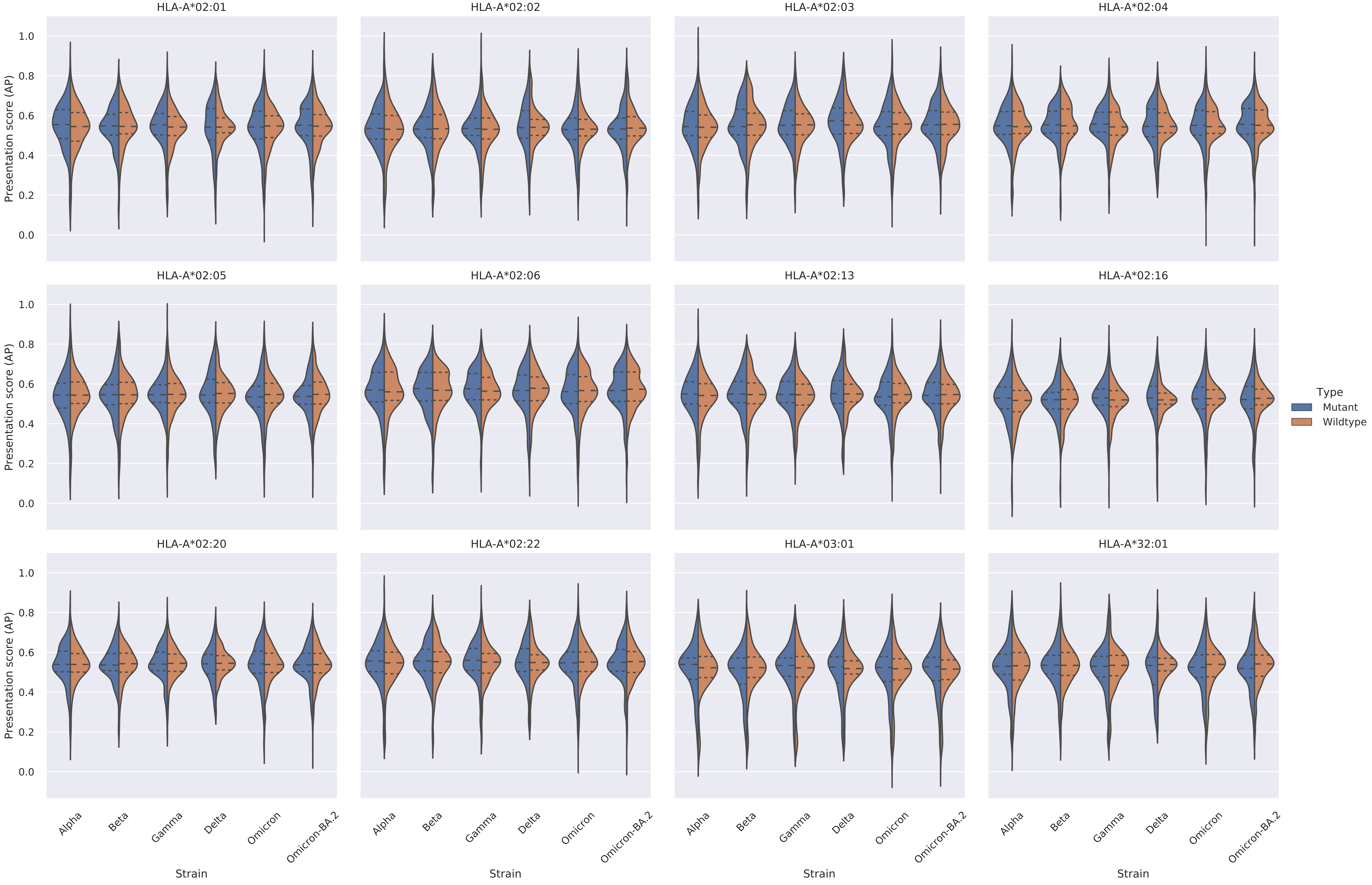

Supplementary Figure 3: Violin plots of the distribution of presentation scores (AP) for mutant and wildtype epitopes per allele for each VOC and Omicron-BA.2. The dashed lines within each violin represent the quartiles. Only entries with an AP score above 0.5 in either the mutant or wildtype were considered for this plot.

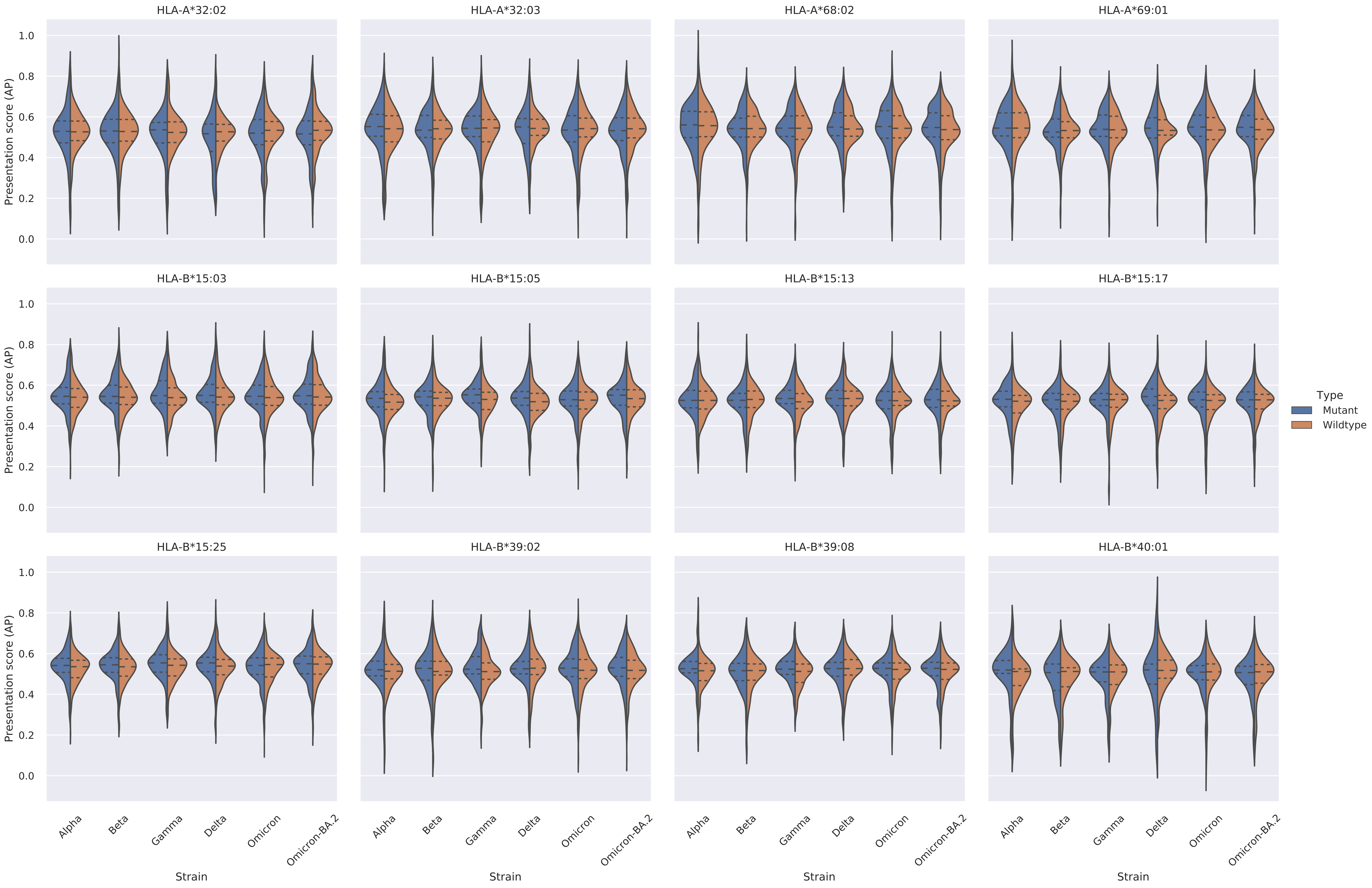

Supplementary Figure 3: Violin plots of the distribution of presentation scores (AP) for mutant and wildtype epitopes per allele for each VOC and Omicron-BA.2. The dashed lines within each violin represent the quartiles. Only entries with an AP score above 0.5 in either the mutant or wildtype were considered for this plot.

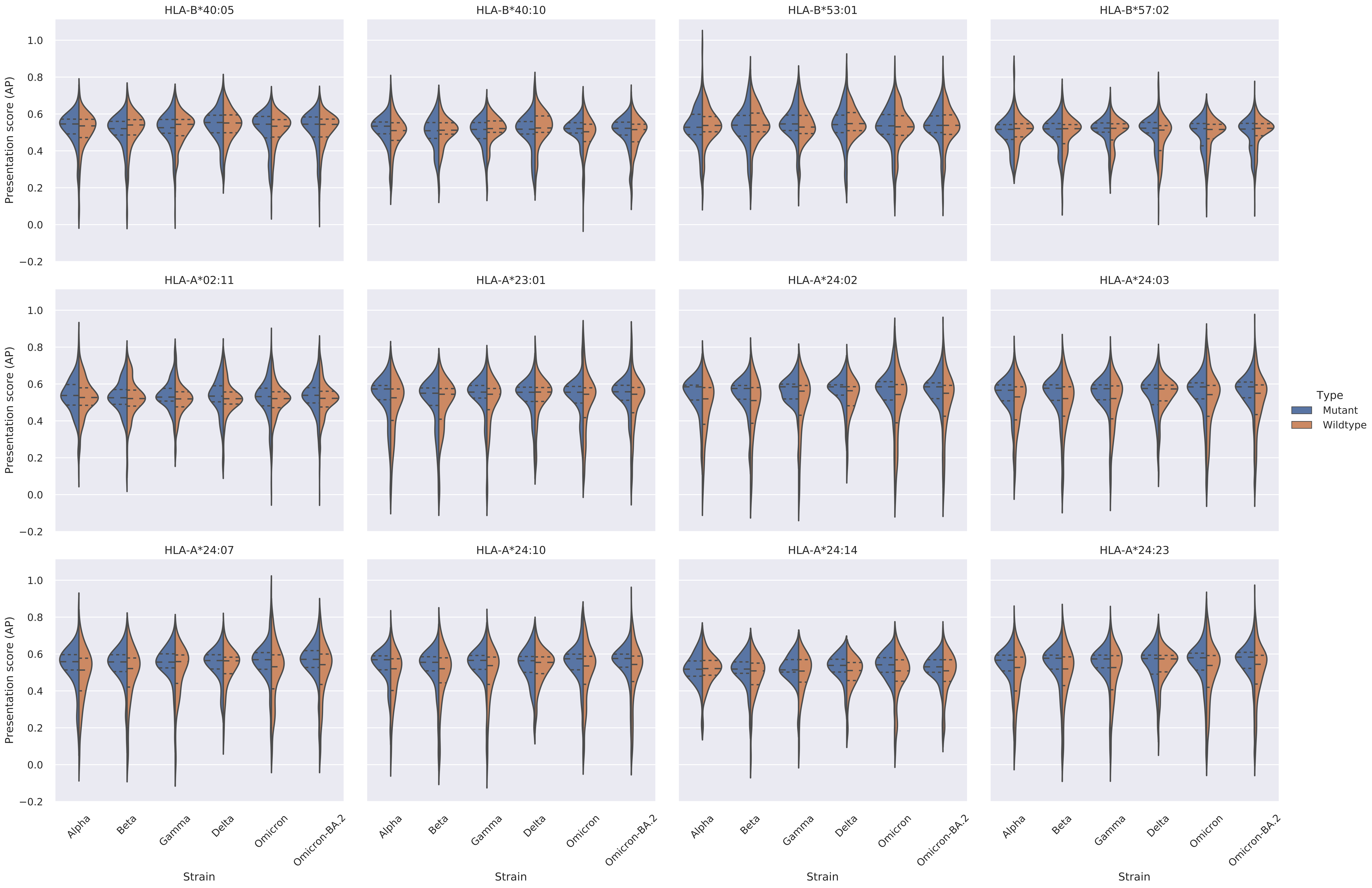

Supplementary Figure 3: Violin plots of the distribution of presentation scores (AP) for mutant and wildtype epitopes per allele for each VOC and Omicron-BA.2. The dashed lines within each violin represent the quartiles. Only entries with an AP score above 0.5 in either the mutant or wildtype were considered for this plot.

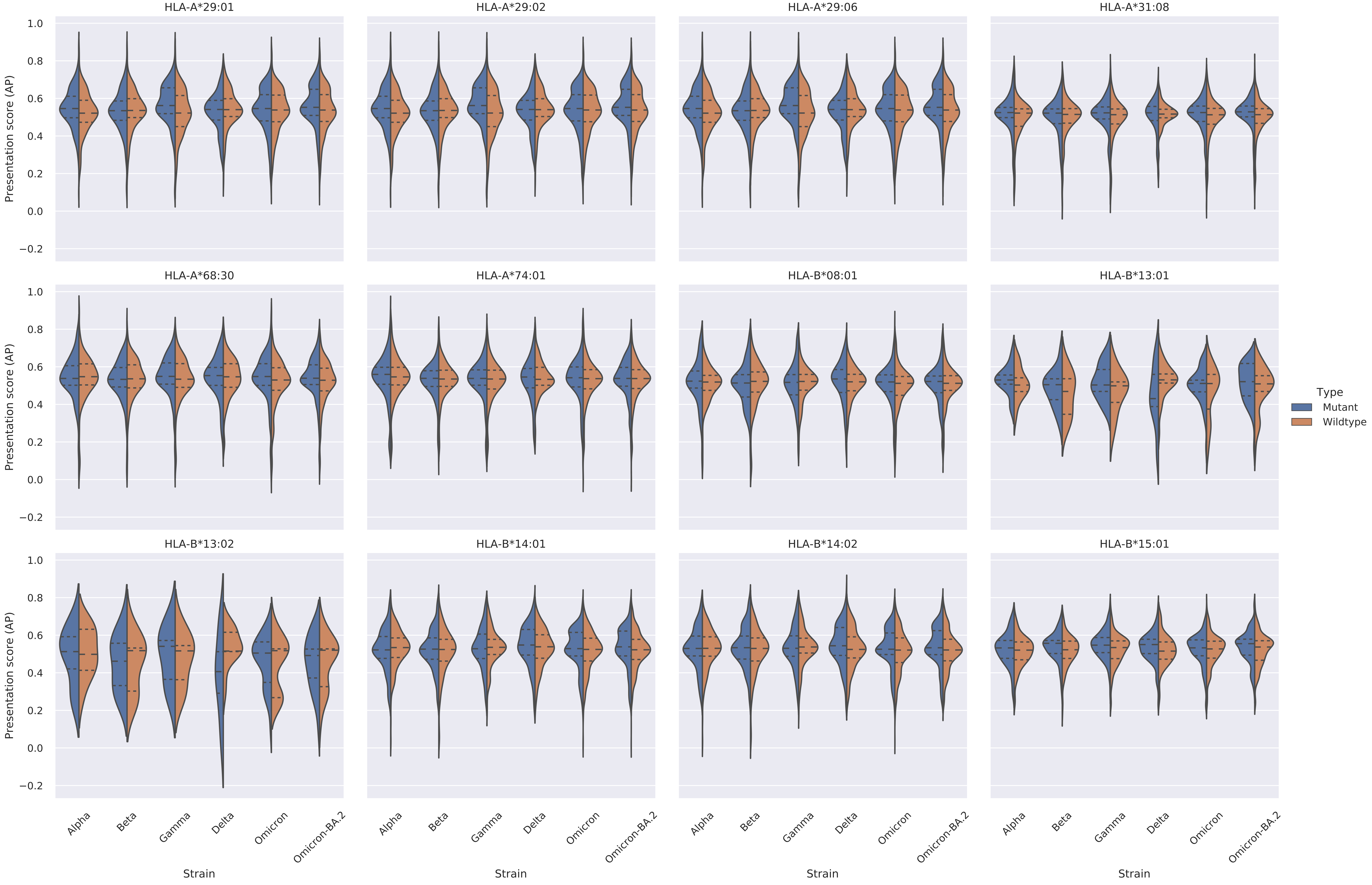

Supplementary Figure 3: Violin plots of the distribution of presentation scores (AP) for mutant and wildtype epitopes per allele for each VOC and Omicron-BA.2. The dashed lines within each violin represent the quartiles. Only entries with an AP score above 0.5 in either the mutant or wildtype were considered for this plot.

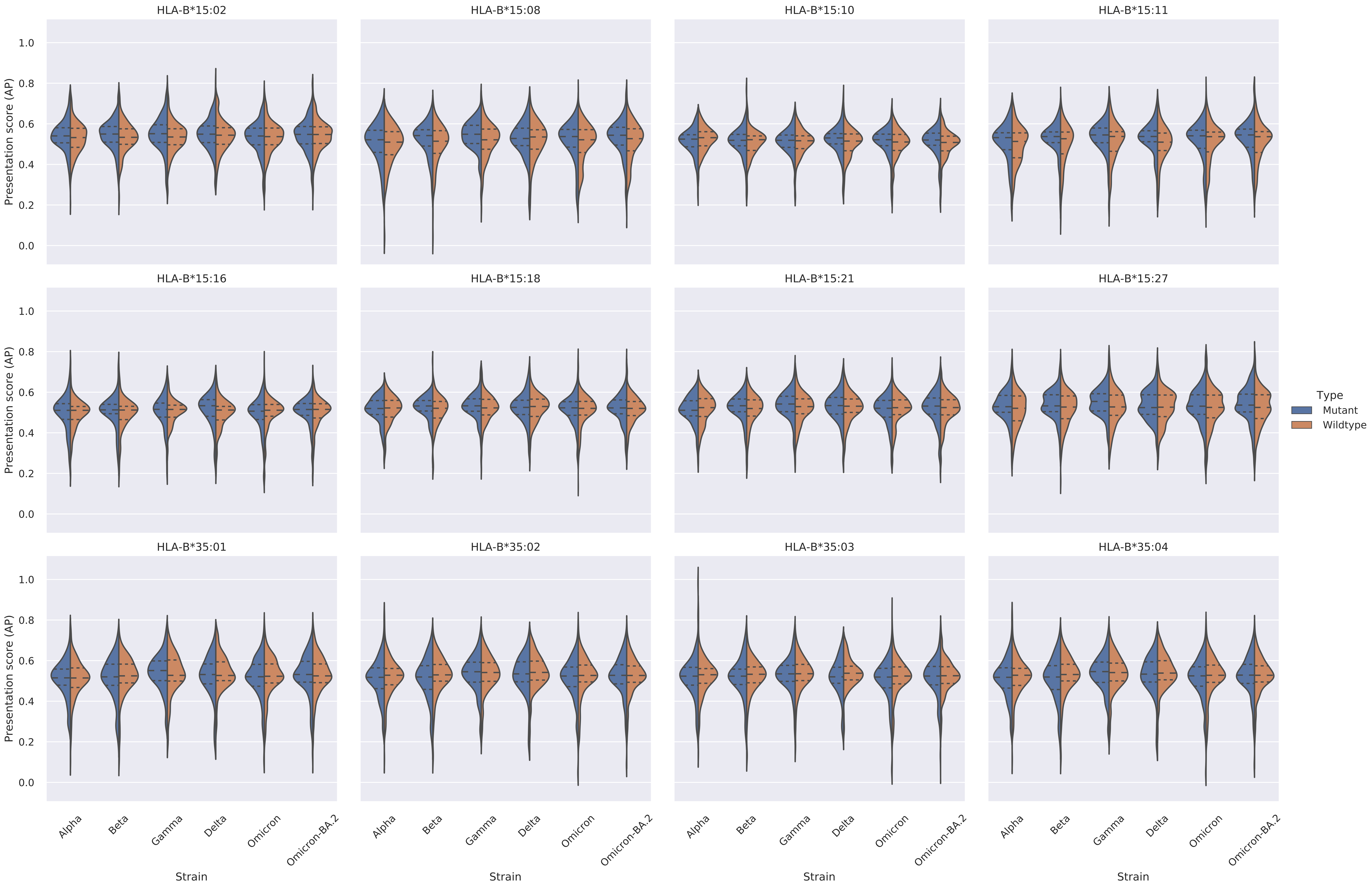

Supplementary Figure 3: Violin plots of the distribution of presentation scores (AP) for mutant and wildtype epitopes per allele for each VOC and Omicron-BA.2. The dashed lines within each violin represent the quartiles. Only entries with an AP score above 0.5 in either the mutant or wildtype were considered for this plot.

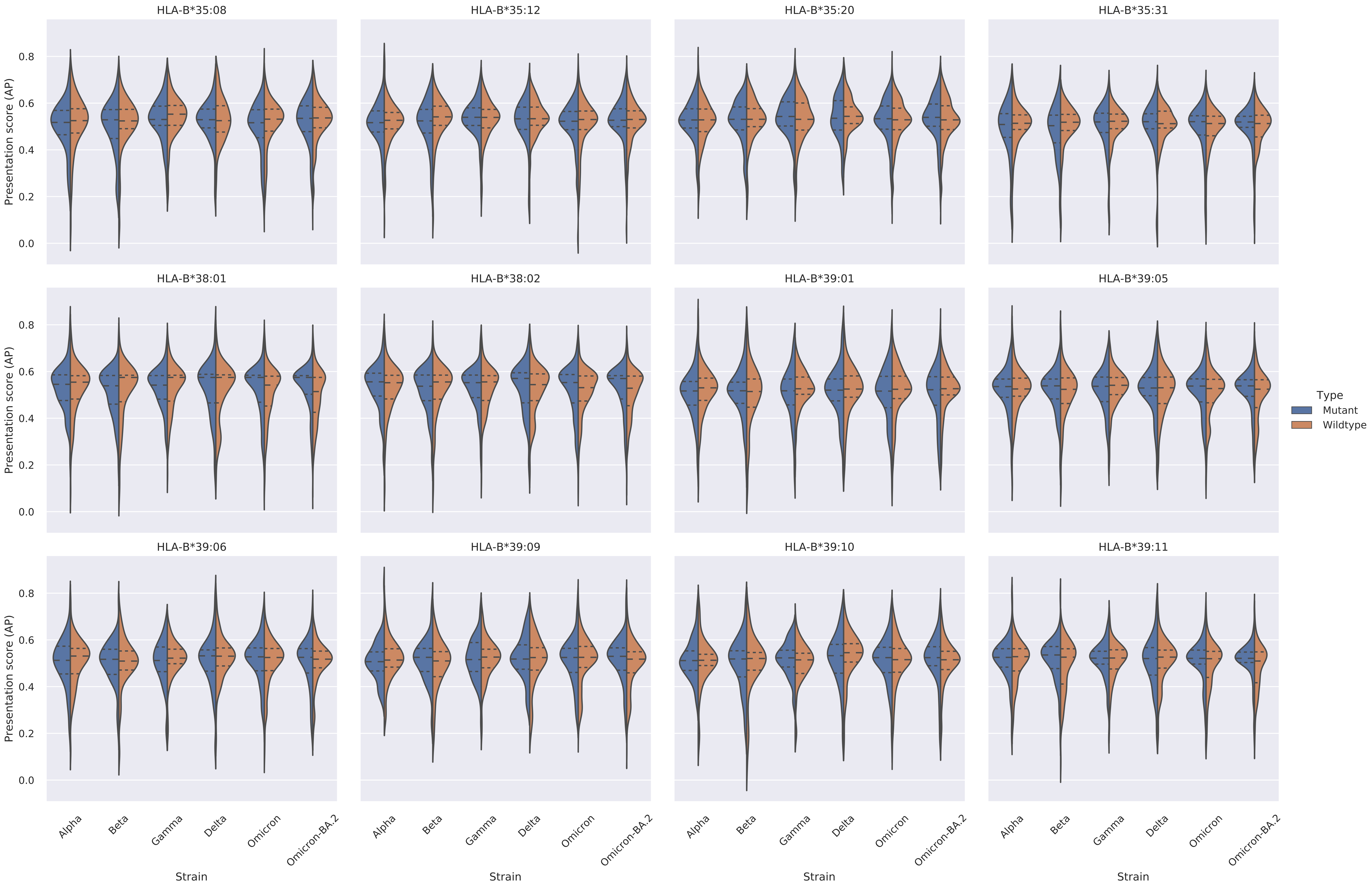

Supplementary Figure 3: Violin plots of the distribution of presentation scores (AP) for mutant and wildtype epitopes per allele for each VOC and Omicron-BA.2. The dashed lines within each violin represent the quartiles. Only entries with an AP score above 0.5 in either the mutant or wildtype were considered for this plot.

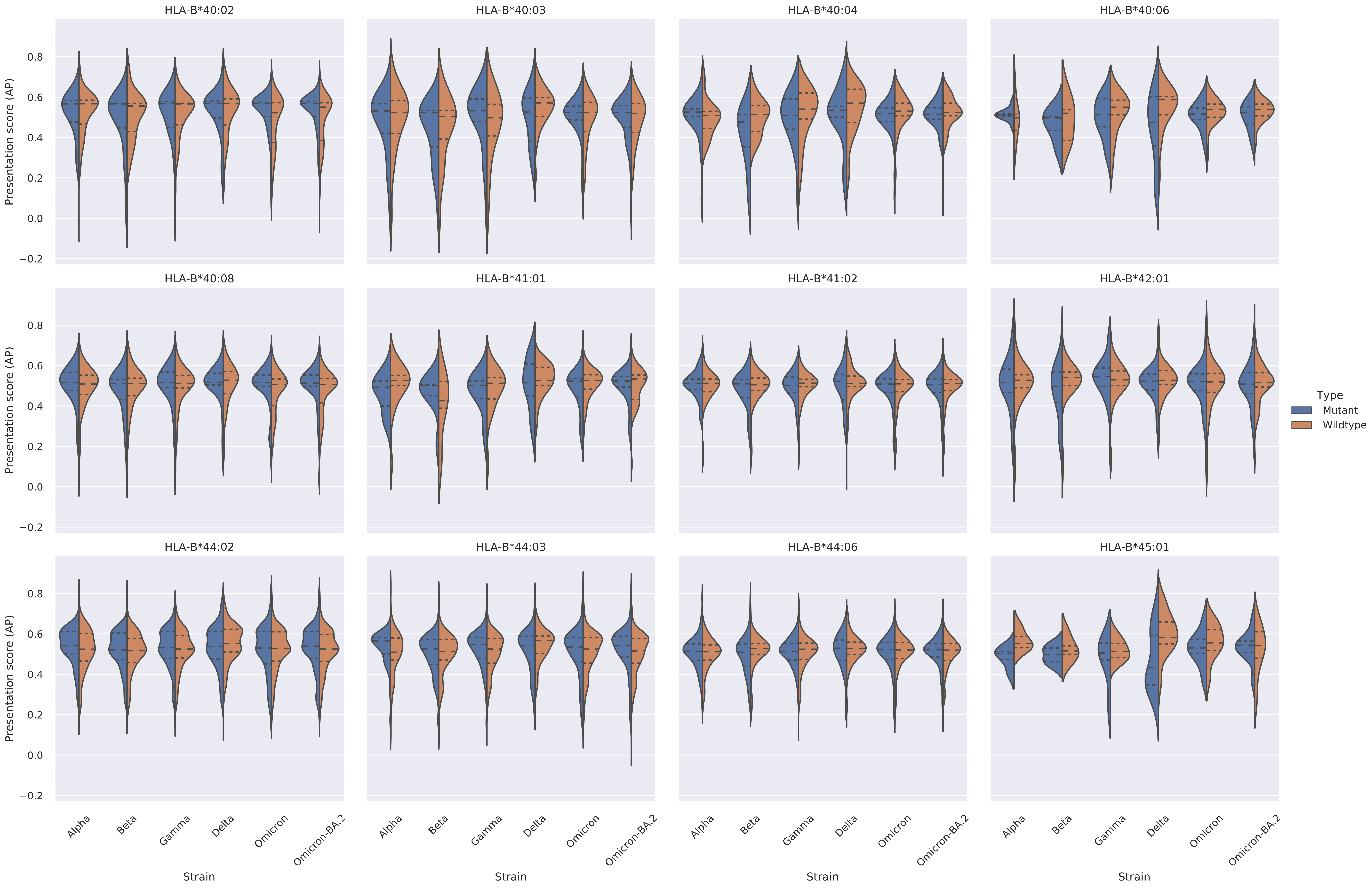

Supplementary Figure 3: Violin plots of the distribution of presentation scores (AP) for mutant and wildtype epitopes per allele for each VOC and Omicron-BA.2. The dashed lines within each violin represent the quartiles. Only entries with an AP score above 0.5 in either the mutant or wildtype were considered for this plot.

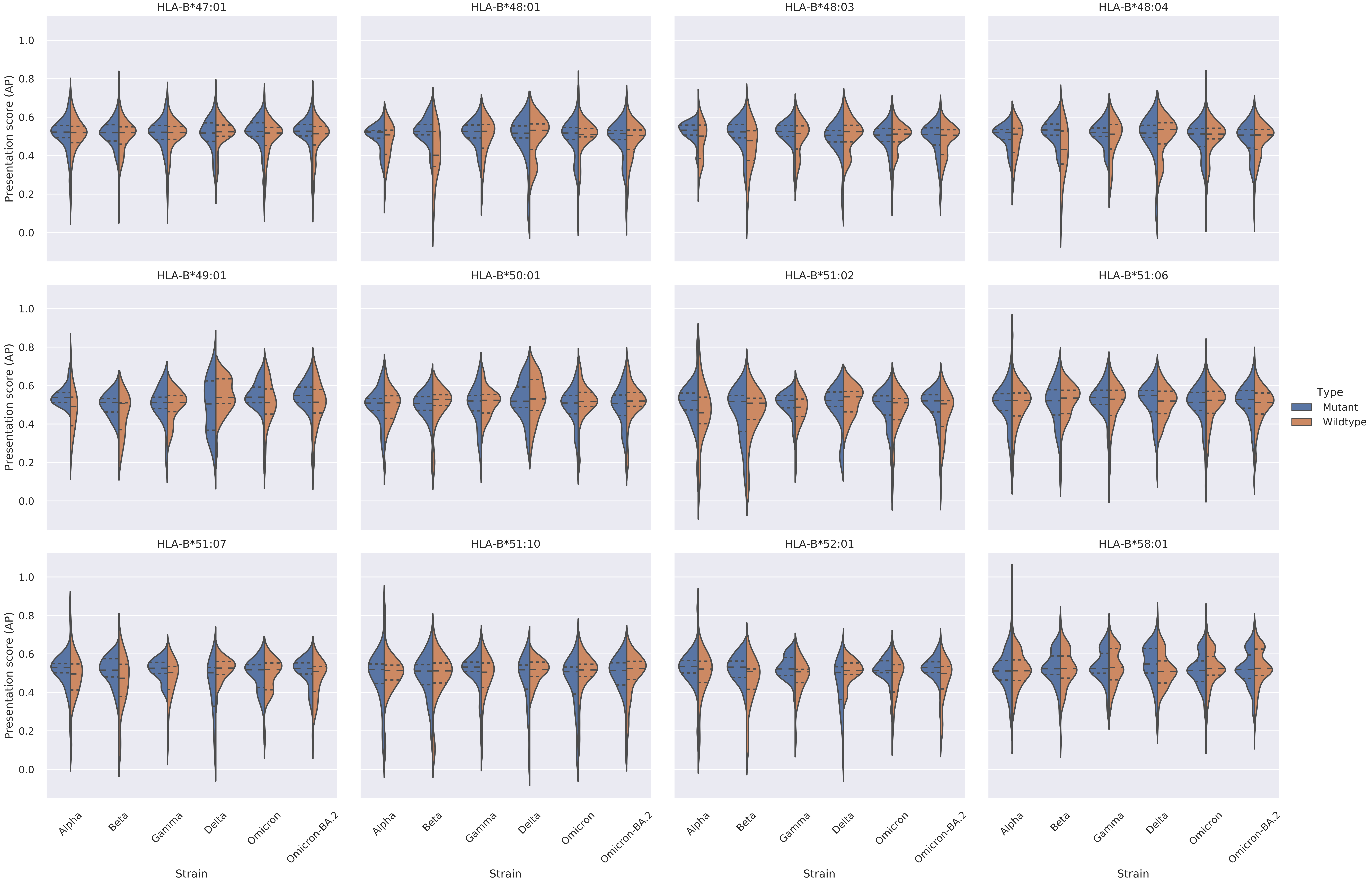

Supplementary Figure 3: Violin plots of the distribution of presentation scores (AP) for mutant and wildtype epitopes per allele for each VOC and Omicron-BA.2. The dashed lines within each violin represent the quartiles. Only entries with an AP score above 0.5 in either the mutant or wildtype were considered for this plot.

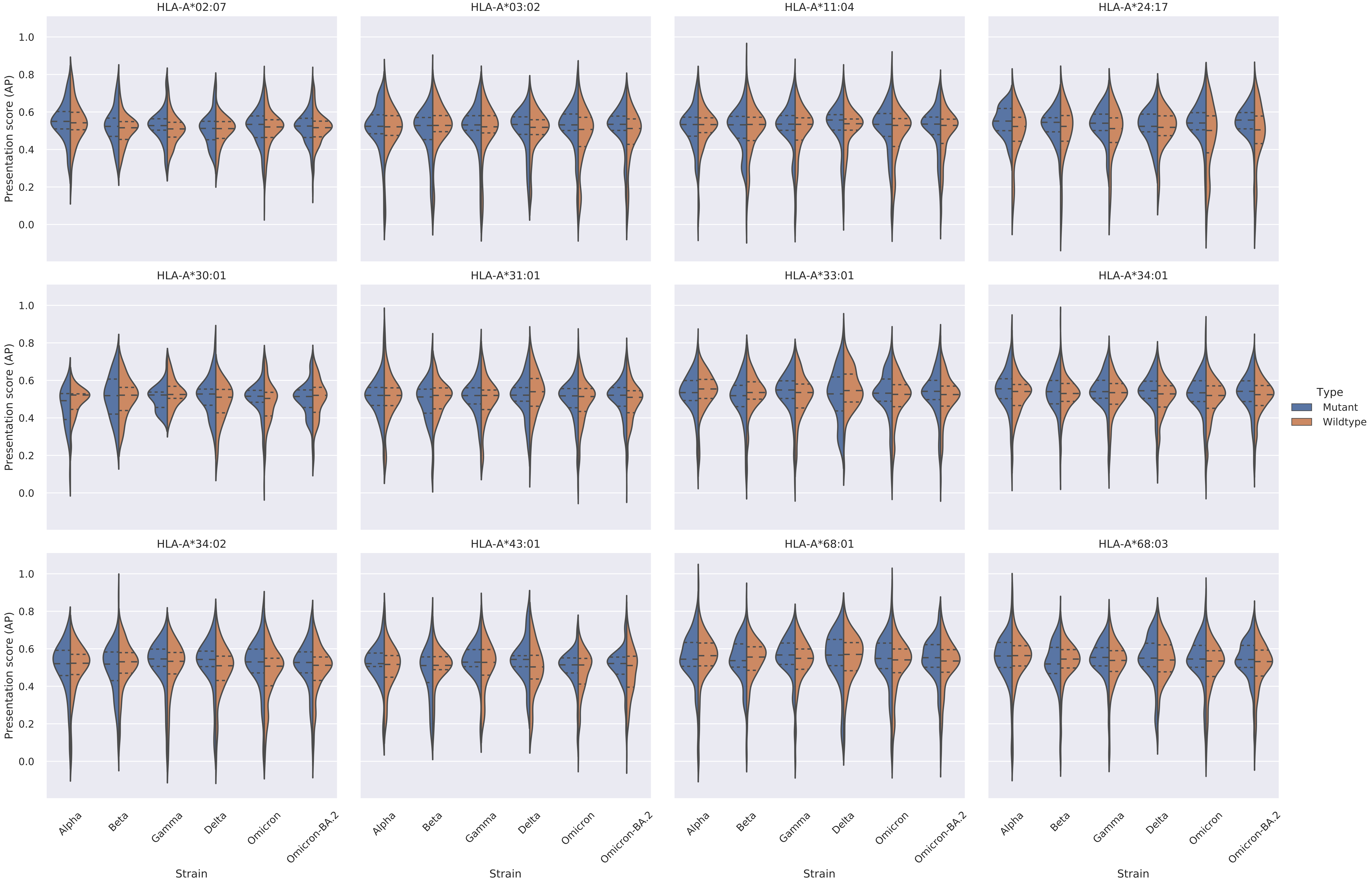

Supplementary Figure 3: Violin plots of the distribution of presentation scores (AP) for mutant and wildtype epitopes per allele for each VOC and Omicron-BA.2. The dashed lines within each violin represent the quartiles. Only entries with an AP score above 0.5 in either the mutant or wildtype were considered for this plot.

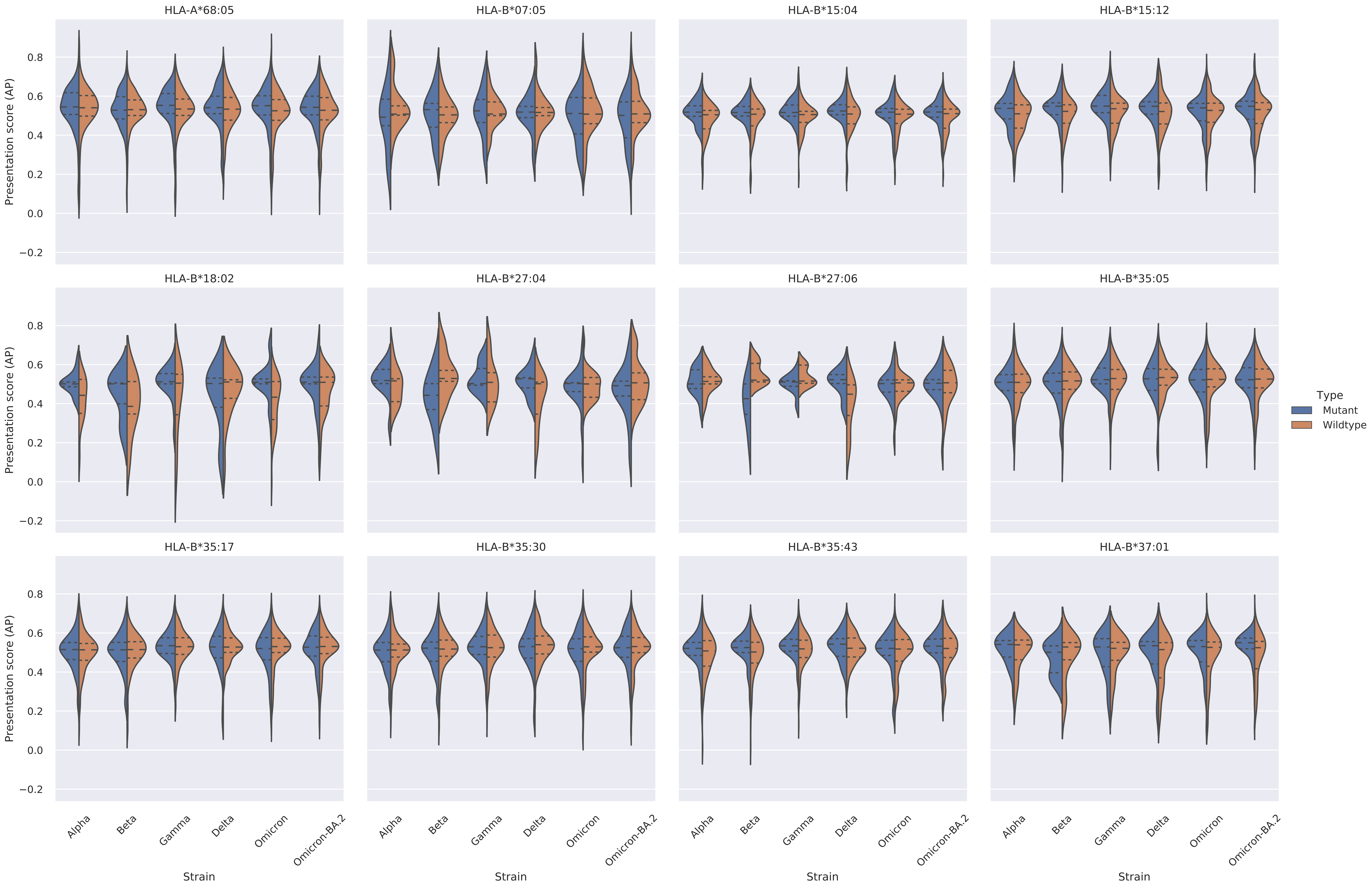

Supplementary Figure 3: Violin plots of the distribution of presentation scores (AP) for mutant and wildtype epitopes per allele for each VOC and Omicron-BA.2. The dashed lines within each violin represent the quartiles. Only entries with an AP score above 0.5 in either the mutant or wildtype were considered for this plot.

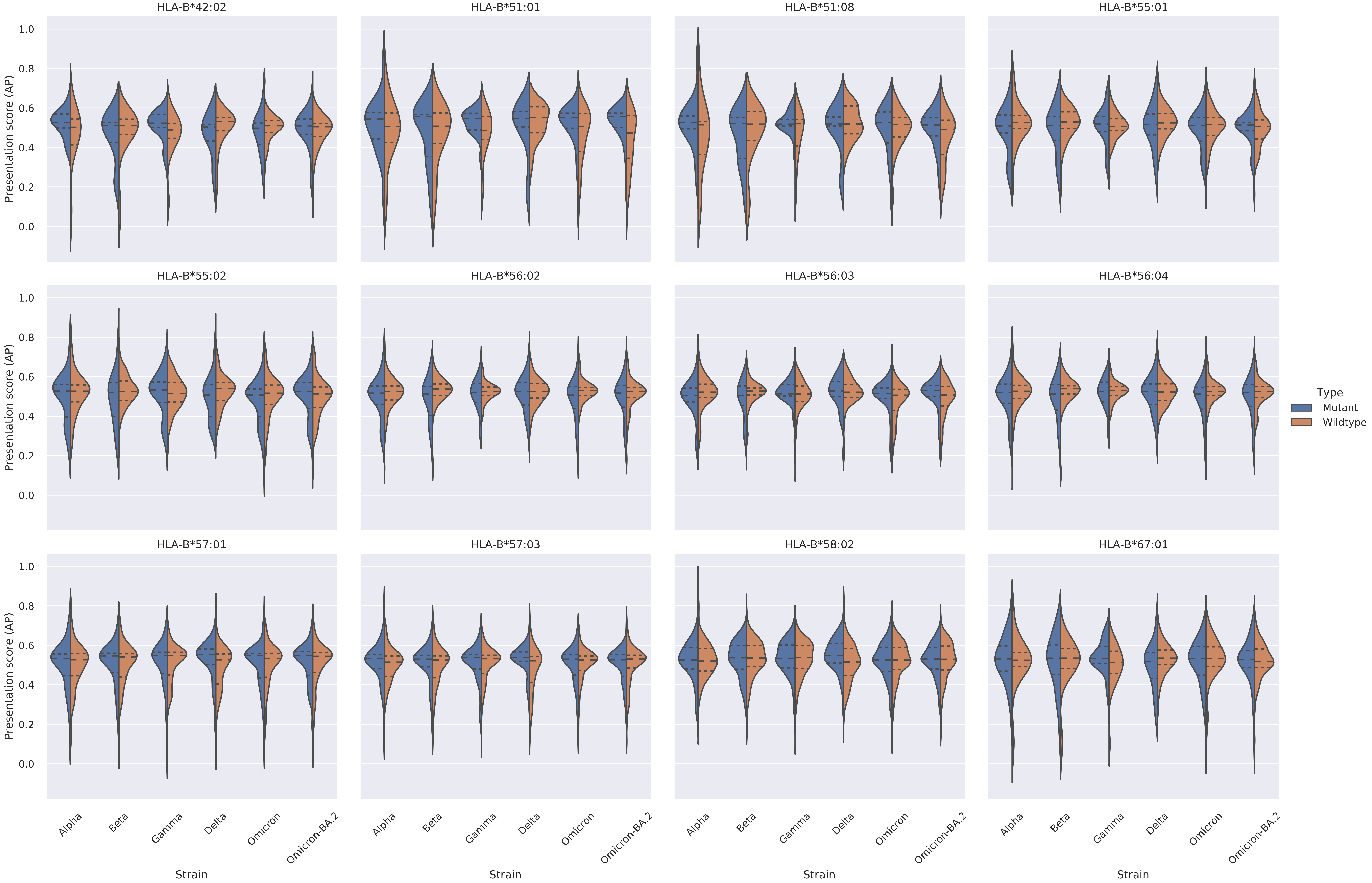

Supplementary Figure 3: Violin plots of the distribution of presentation scores (AP) for mutant and wildtype epitopes per allele for each VOC and Omicron-BA.2. The dashed lines within each violin represent the quartiles. Only entries with an AP score above 0.5 in either the mutant or wildtype were considered for this plot.

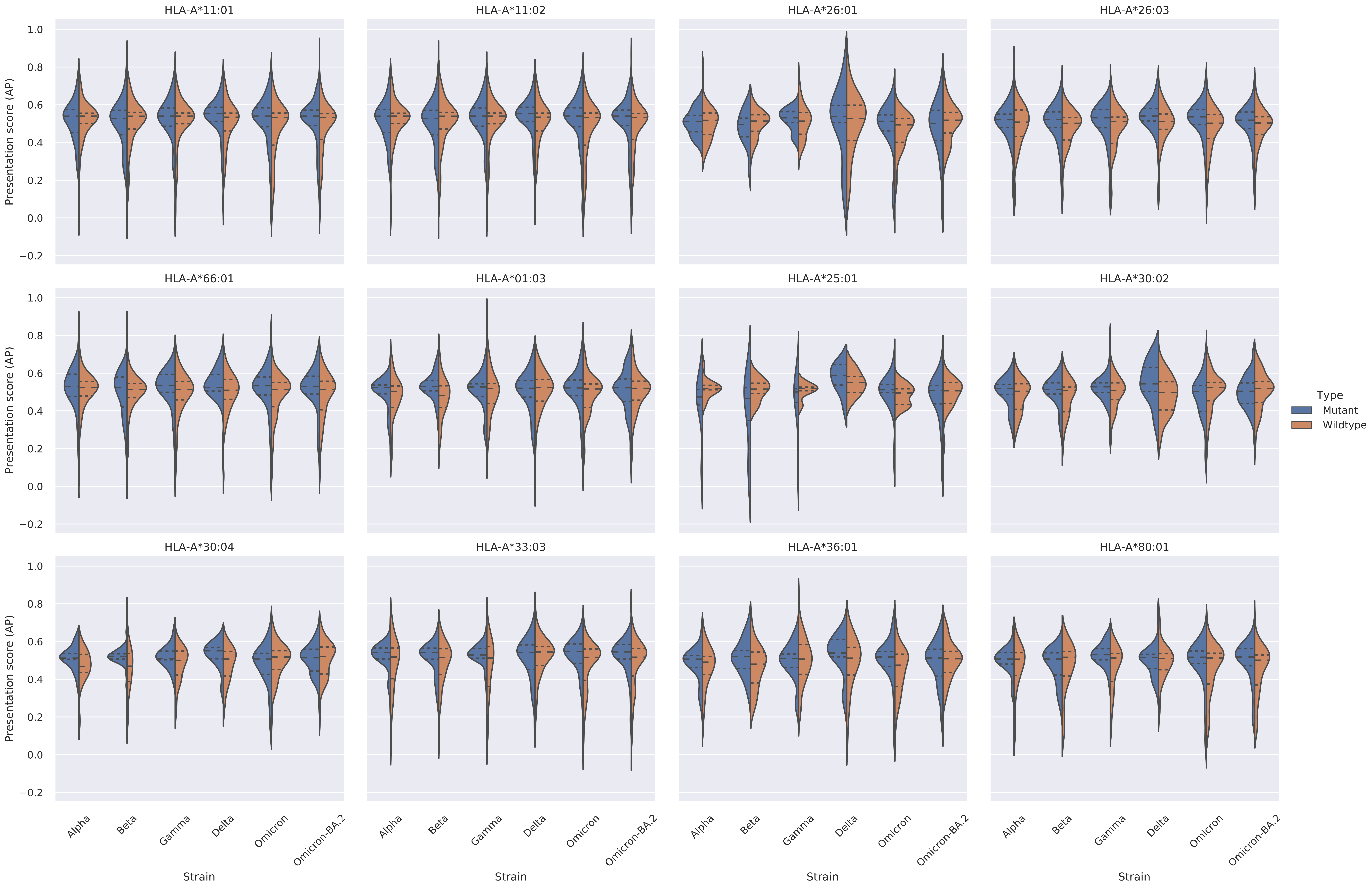

Supplementary Figure 3: Violin plots of the distribution of presentation scores (AP) for mutant and wildtype epitopes per allele for each VOC and Omicron-BA.2. The dashed lines within each violin represent the quartiles. Only entries with an AP score above 0.5 in either the mutant or wildtype were considered for this plot.

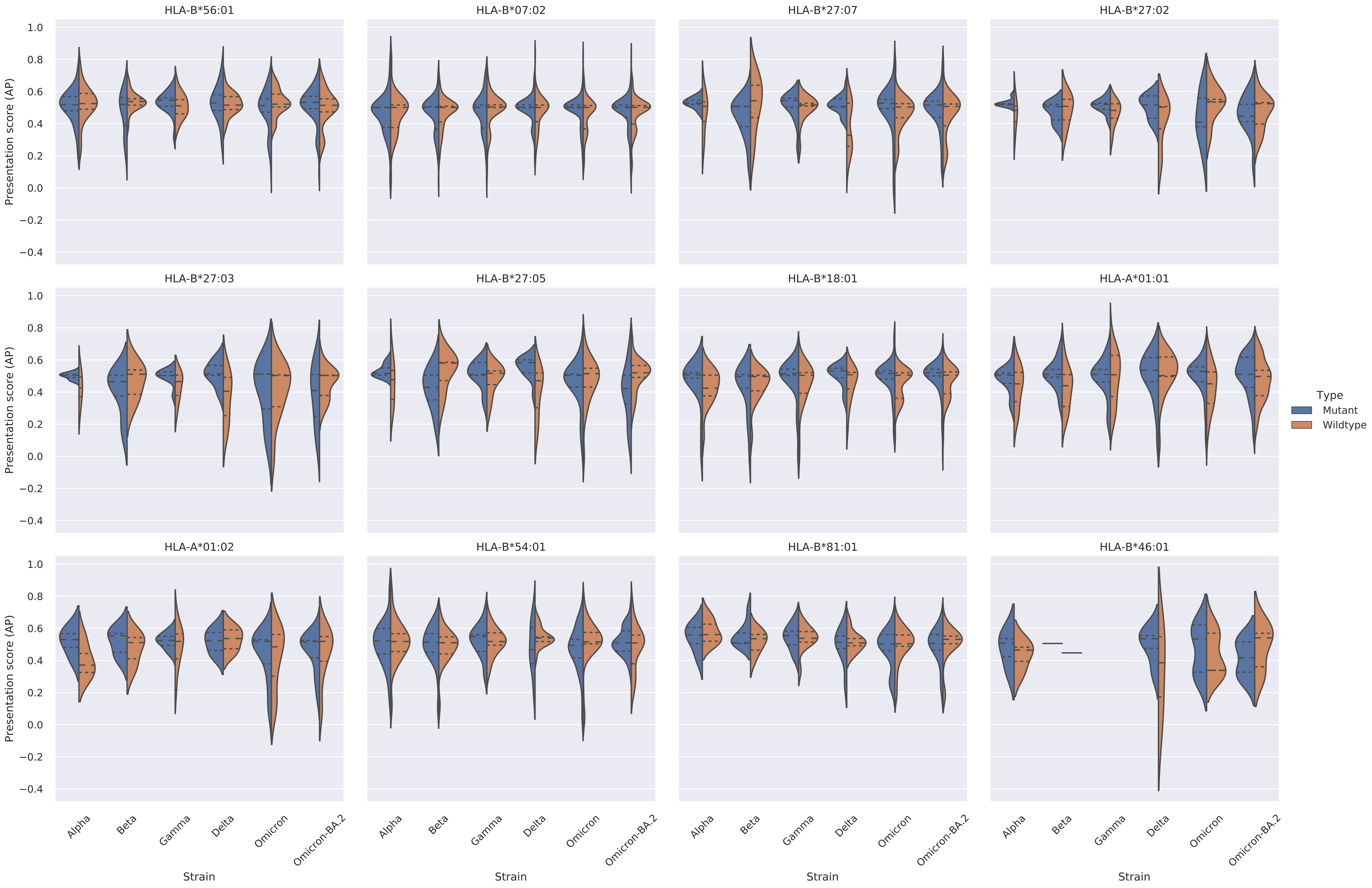

Supplementary Figure 4: Violin plots of the distribution of presentation scores (AP) for mutant and wildtype epitopes per allele for each VOC and Omicron-BA.2. The dashed lines within each violin represent the quartiles.

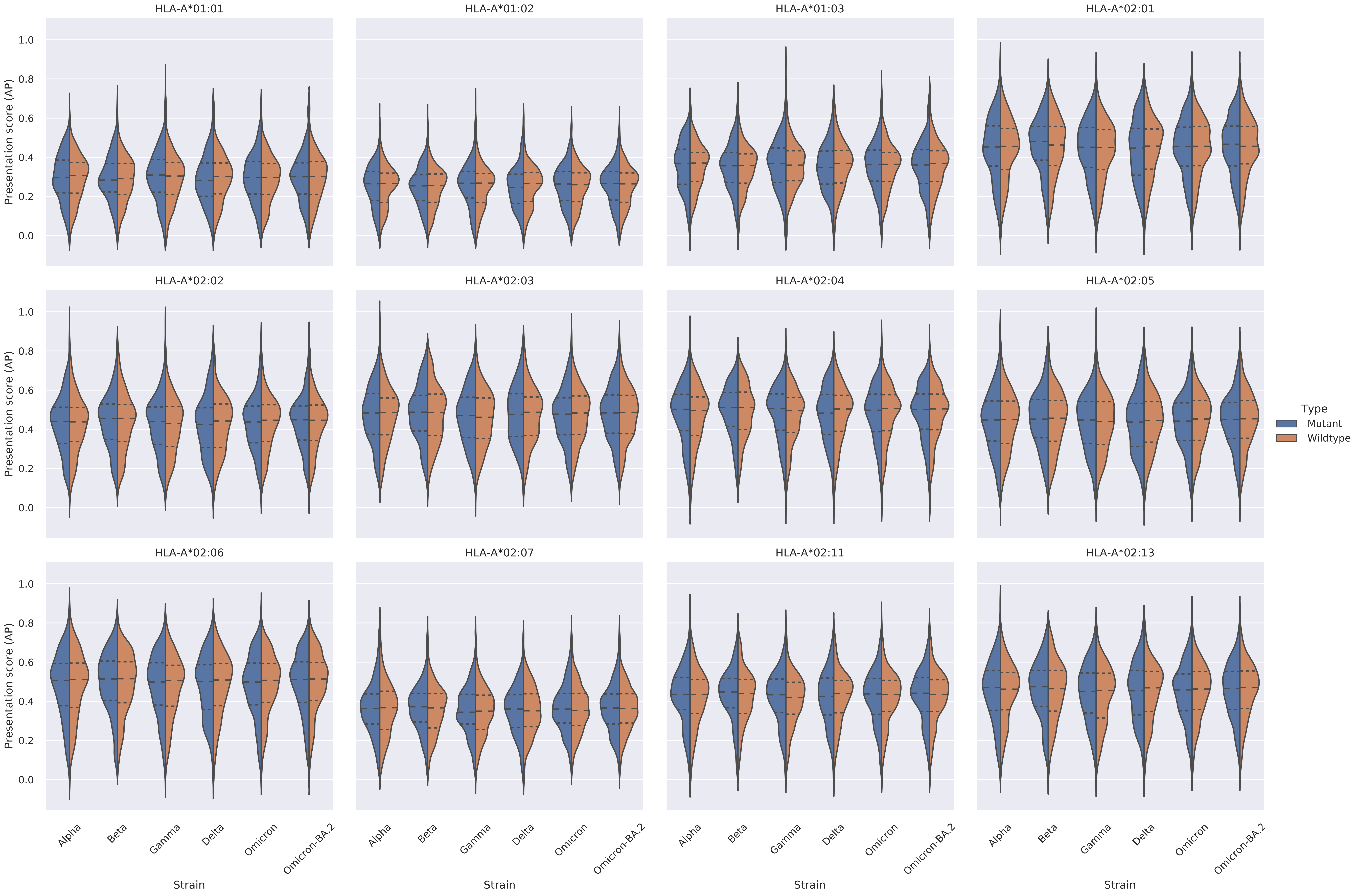

Supplementary Figure 4: Violin plots of the distribution of presentation scores (AP) for mutant and wildtype epitopes per allele for each VOC and Omicron-BA.2. The dashed lines within each violin represent the quartiles.

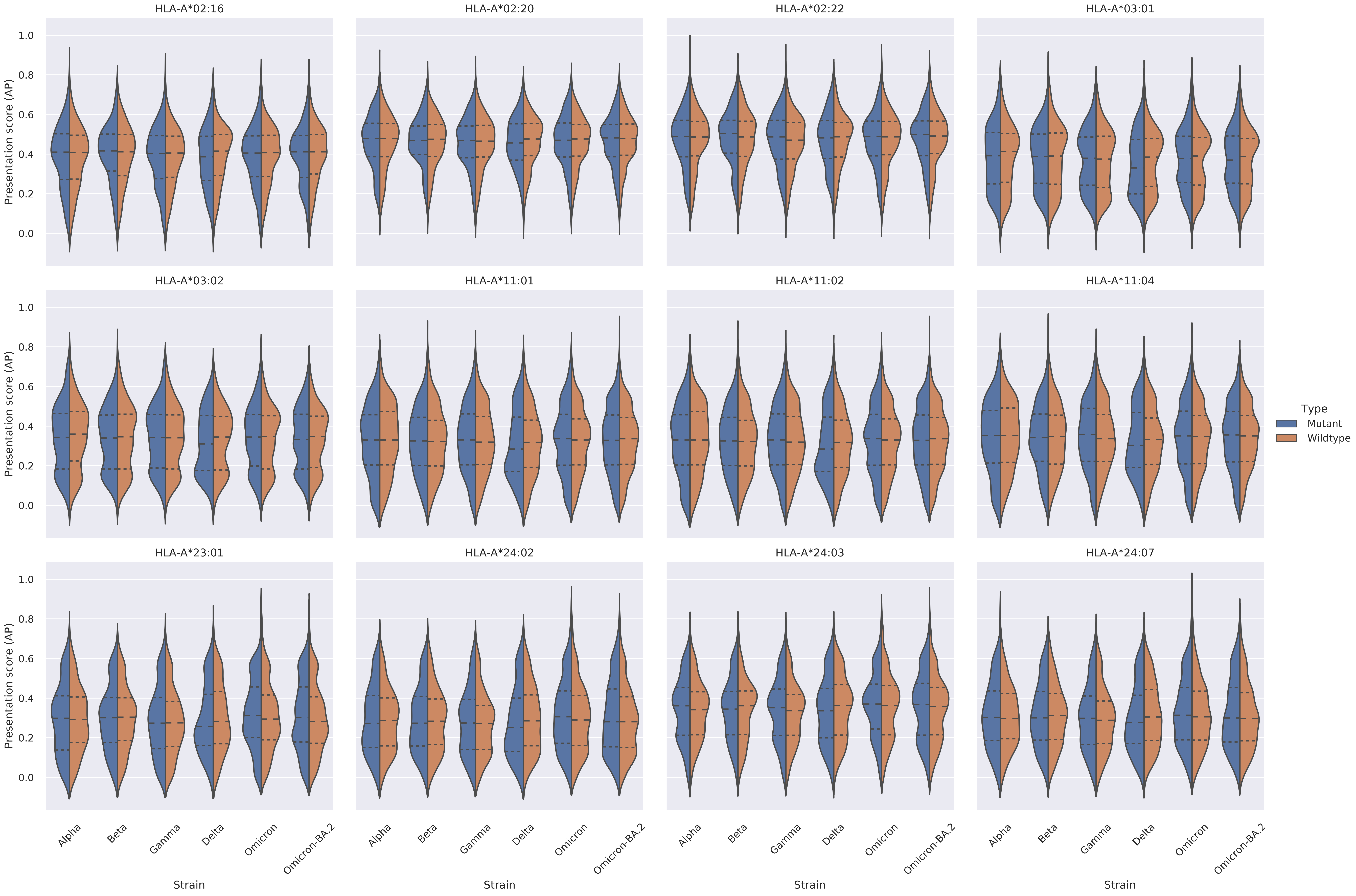

Supplementary Figure 4: Violin plots of the distribution of presentation scores (AP) for mutant and wildtype epitopes per allele for each VOC and Omicron-BA.2. The dashed lines within each violin represent the quartiles.

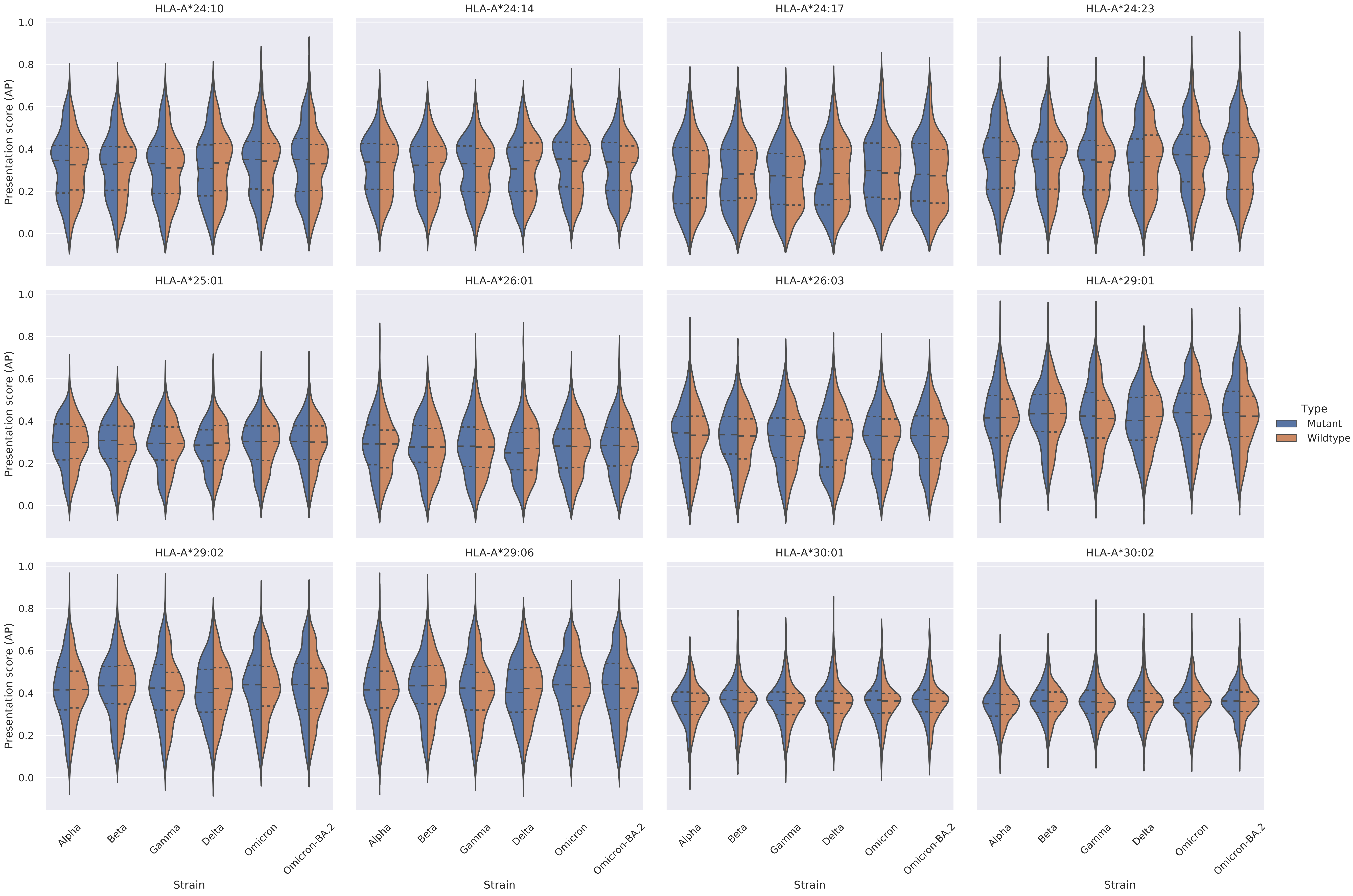

Supplementary Figure 4: Violin plots of the distribution of presentation scores (AP) for mutant and wildtype epitopes per allele for each VOC and Omicron-BA.2. The dashed lines within each violin represent the quartiles.

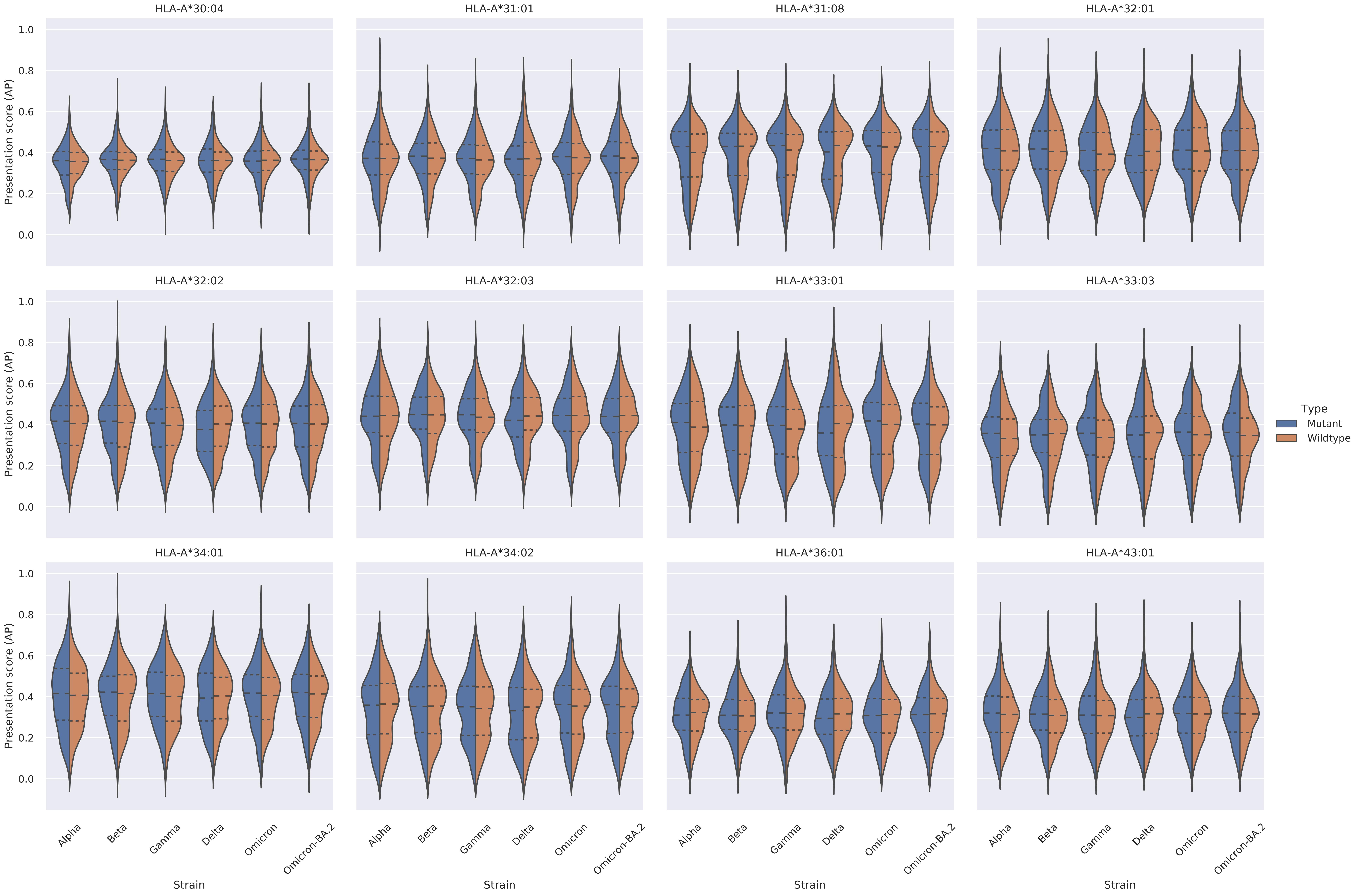

Supplementary Figure 4: Violin plots of the distribution of presentation scores (AP) for mutant and wildtype epitopes per allele for each VOC and Omicron-BA.2. The dashed lines within each violin represent the quartiles.

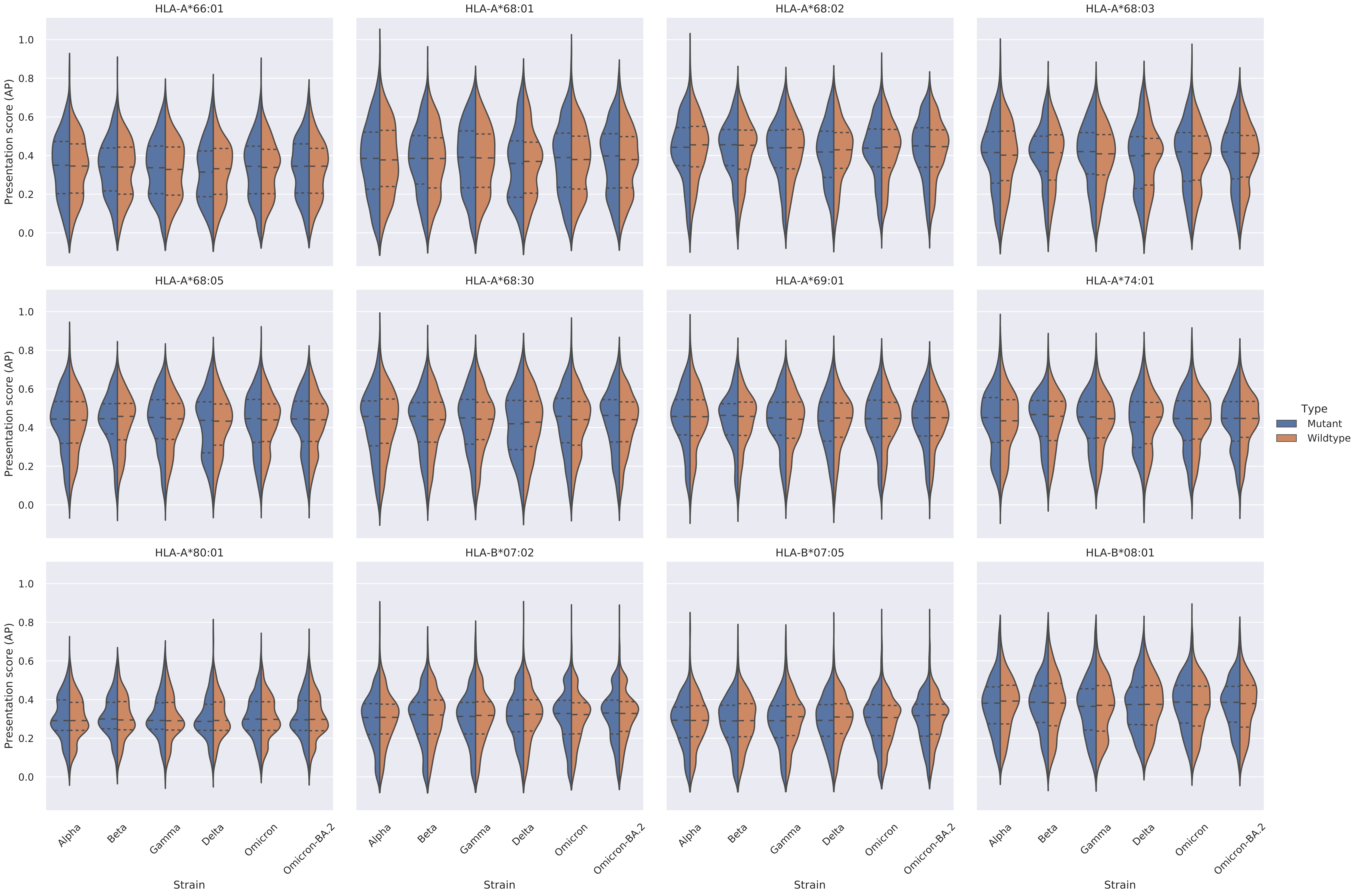

Supplementary Figure 4: Violin plots of the distribution of presentation scores (AP) for mutant and wildtype epitopes per allele for each VOC and Omicron-BA.2. The dashed lines within each violin represent the quartiles.

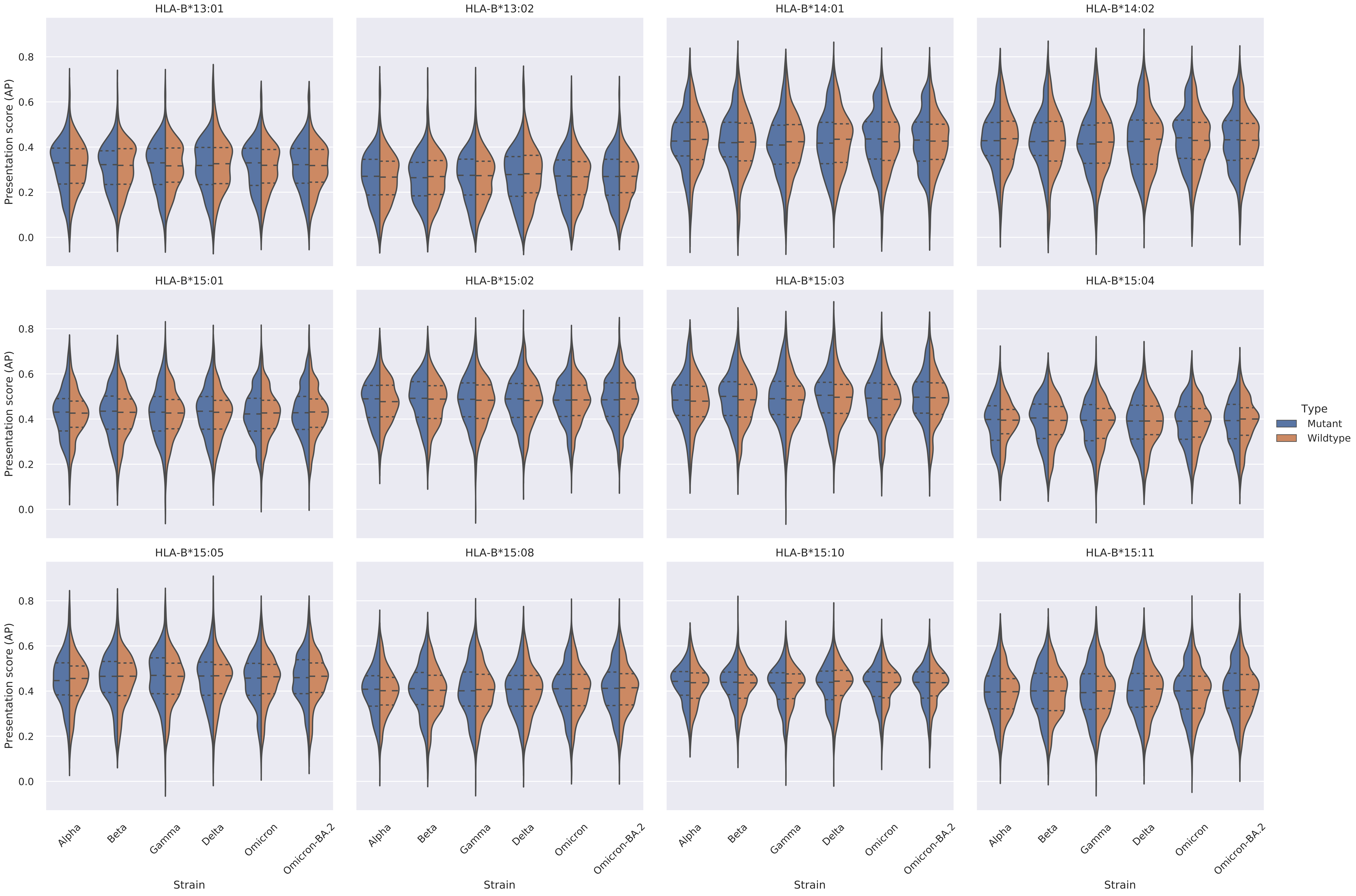

Supplementary Figure 4: Violin plots of the distribution of presentation scores (AP) for mutant and wildtype epitopes per allele for each VOC and Omicron-BA.2. The dashed lines within each violin represent the quartiles.

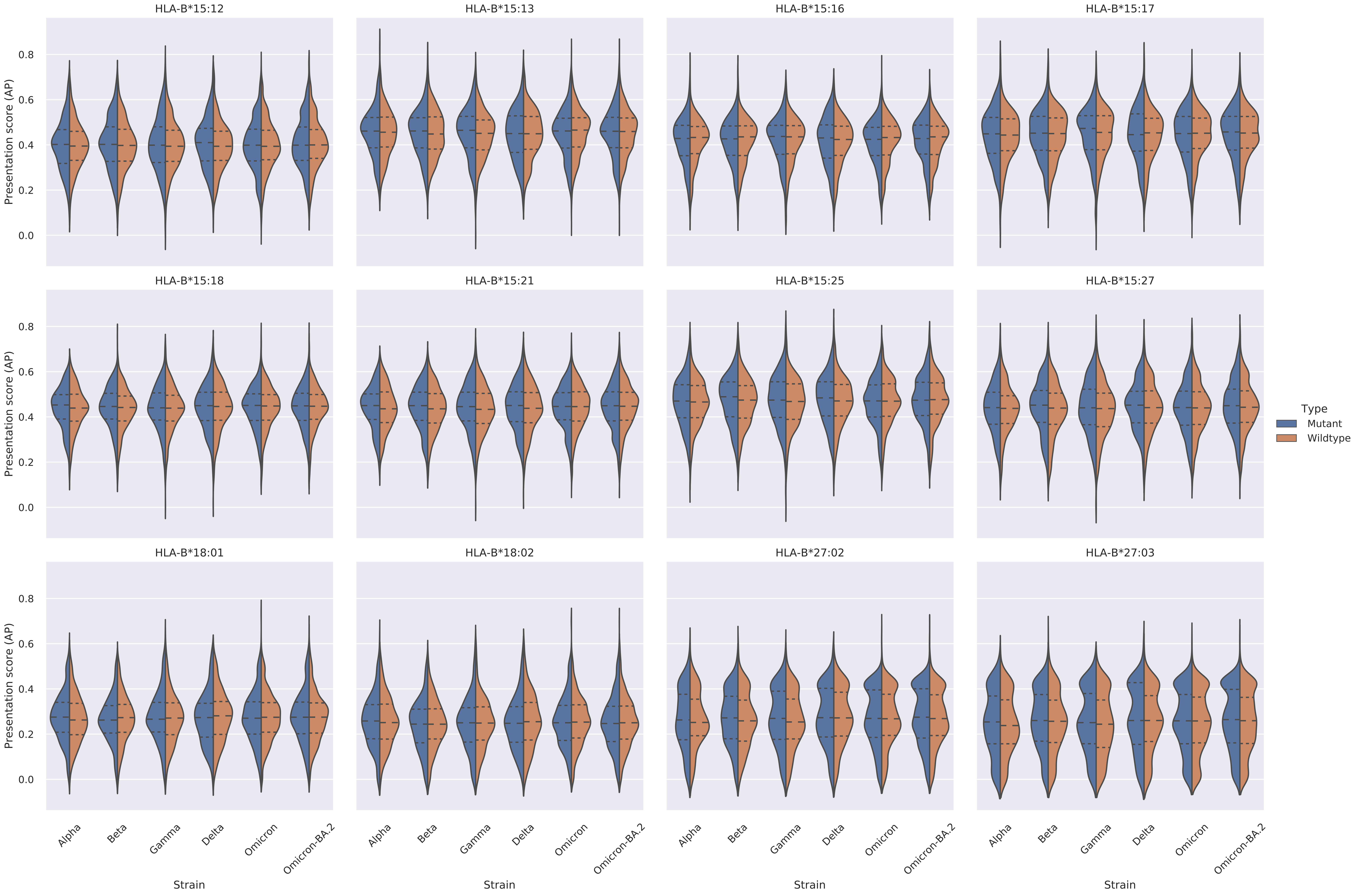

Supplementary Figure 4: Violin plots of the distribution of presentation scores (AP) for mutant and wildtype epitopes per allele for each VOC and Omicron-BA.2. The dashed lines within each violin represent the quartiles.

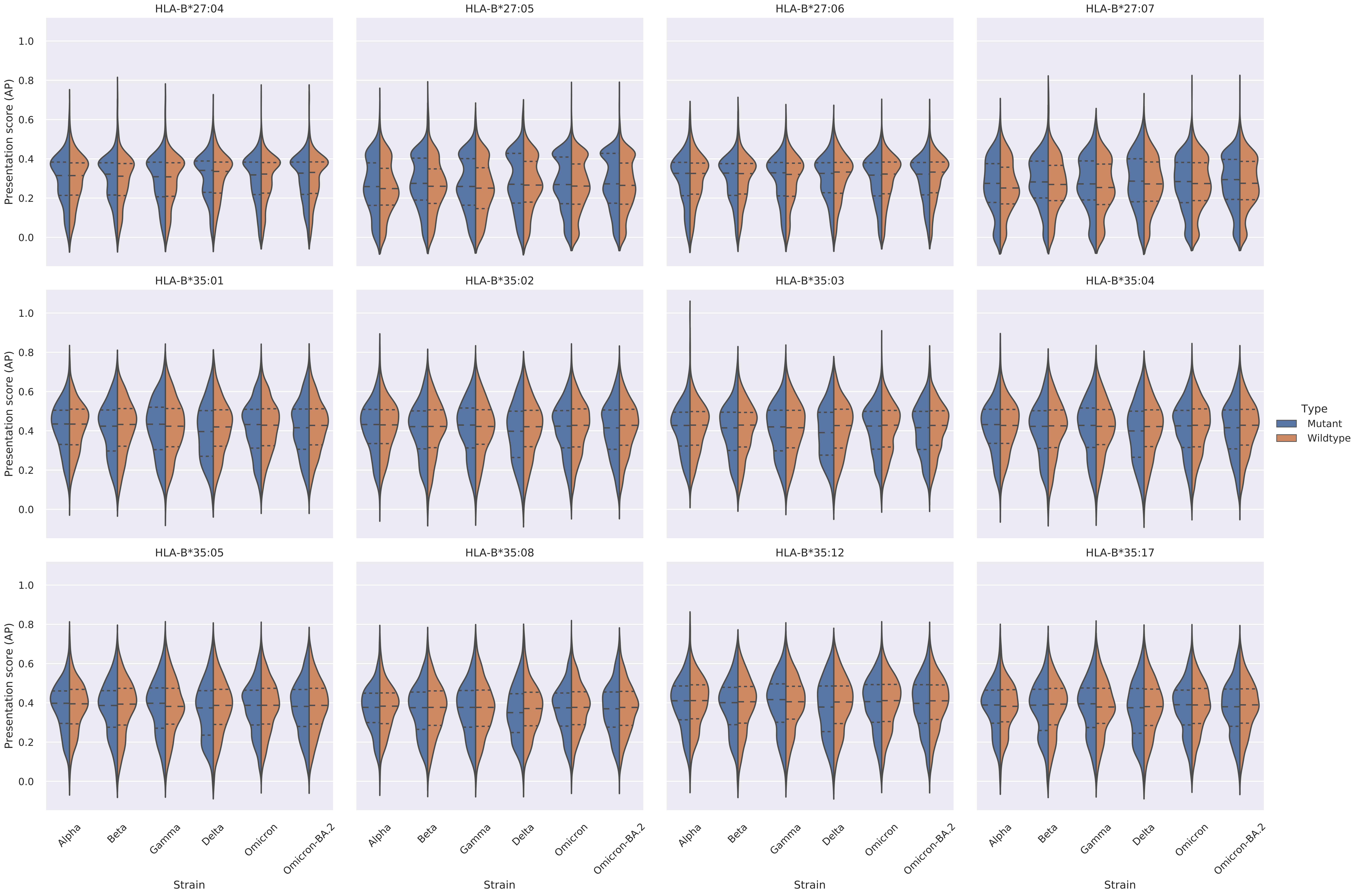

Supplementary Figure 4: Violin plots of the distribution of presentation scores (AP) for mutant and wildtype epitopes per allele for each VOC and Omicron-BA.2. The dashed lines within each violin represent the quartiles.

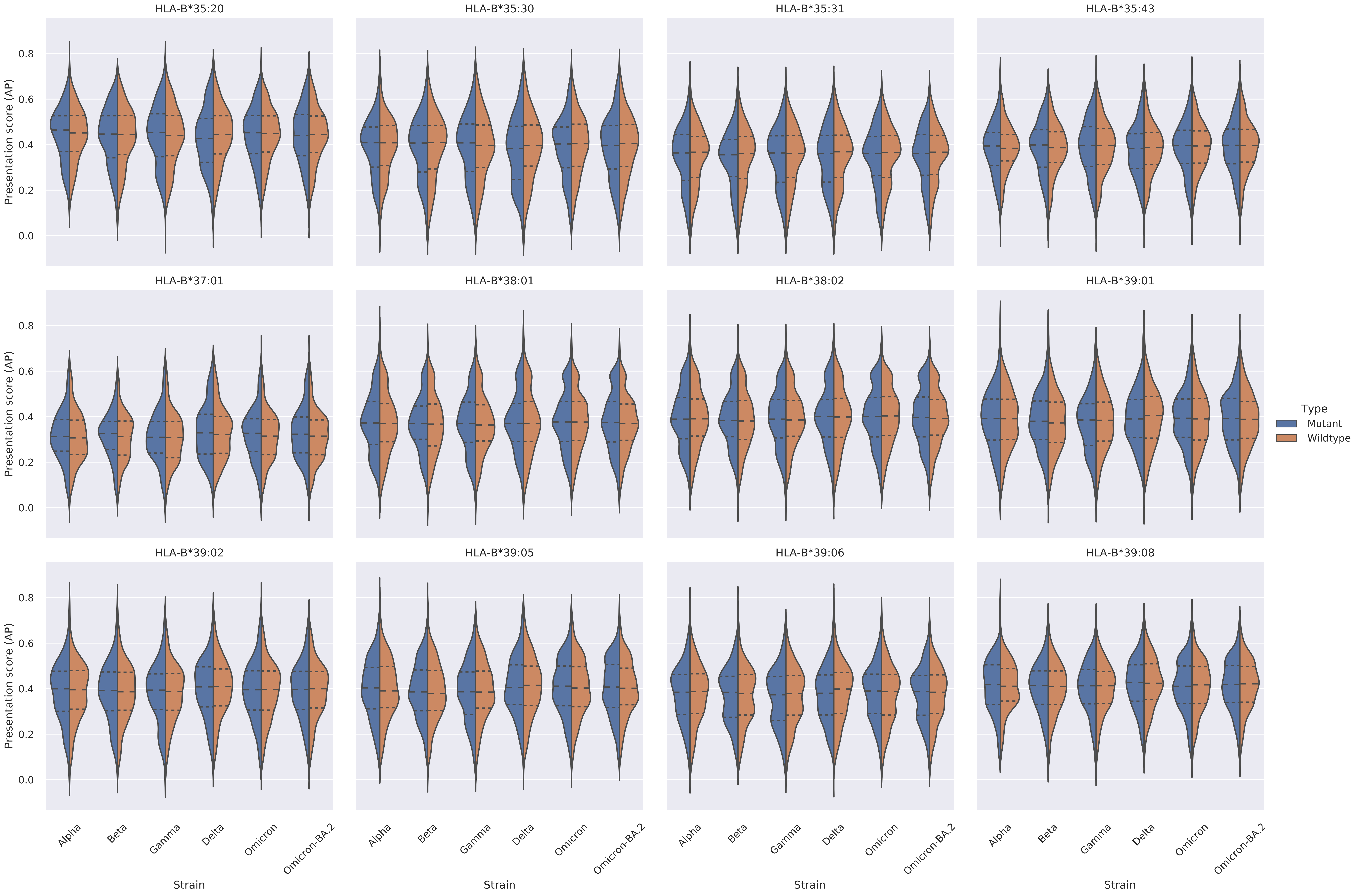

Supplementary Figure 4: Violin plots of the distribution of presentation scores (AP) for mutant and wildtype epitopes per allele for each VOC and Omicron-BA.2. The dashed lines within each violin represent the quartiles.

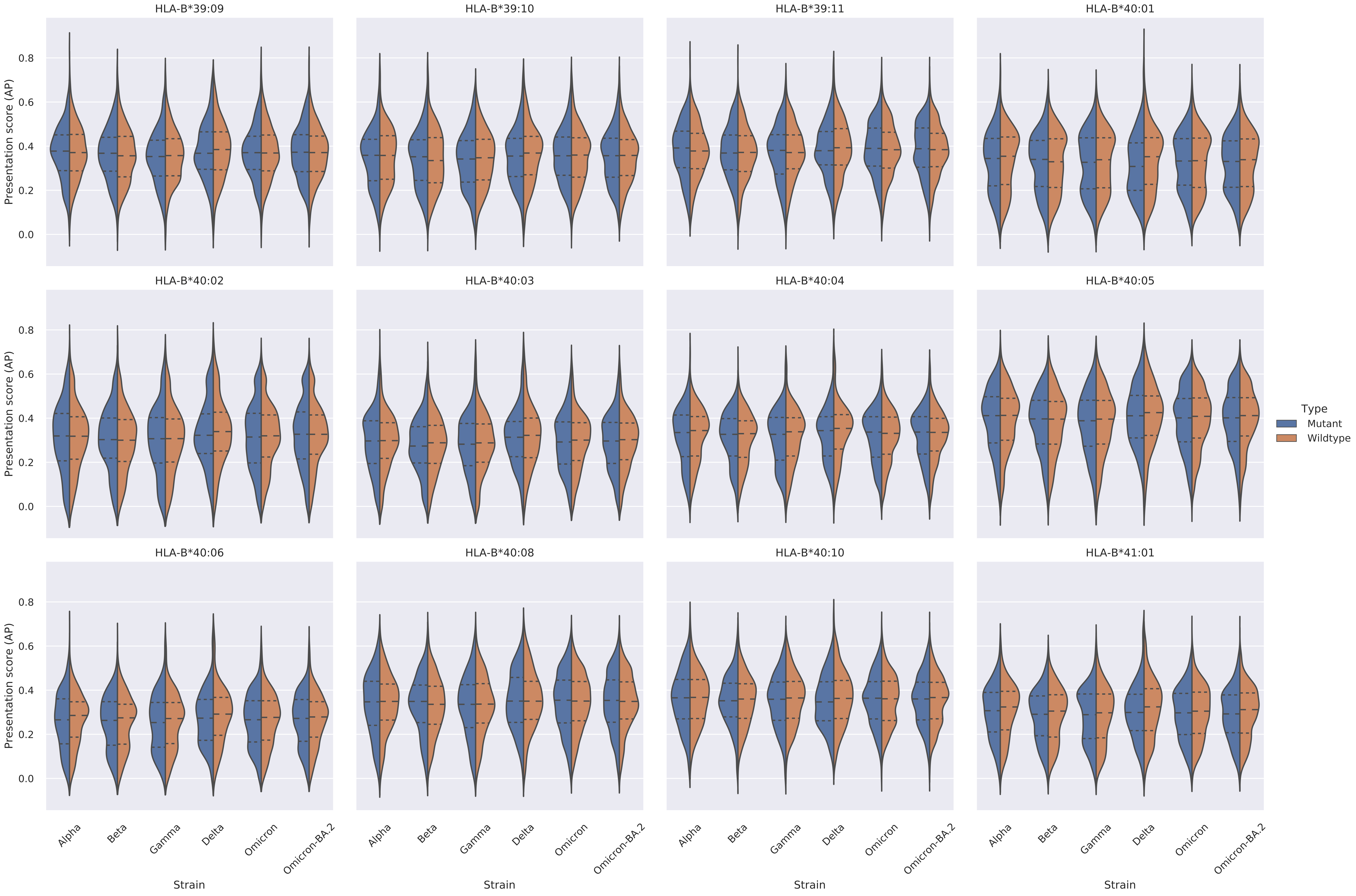

Supplementary Figure 4: Violin plots of the distribution of presentation scores (AP) for mutant and wildtype epitopes per allele for each VOC and Omicron-BA.2. The dashed lines within each violin represent the quartiles.

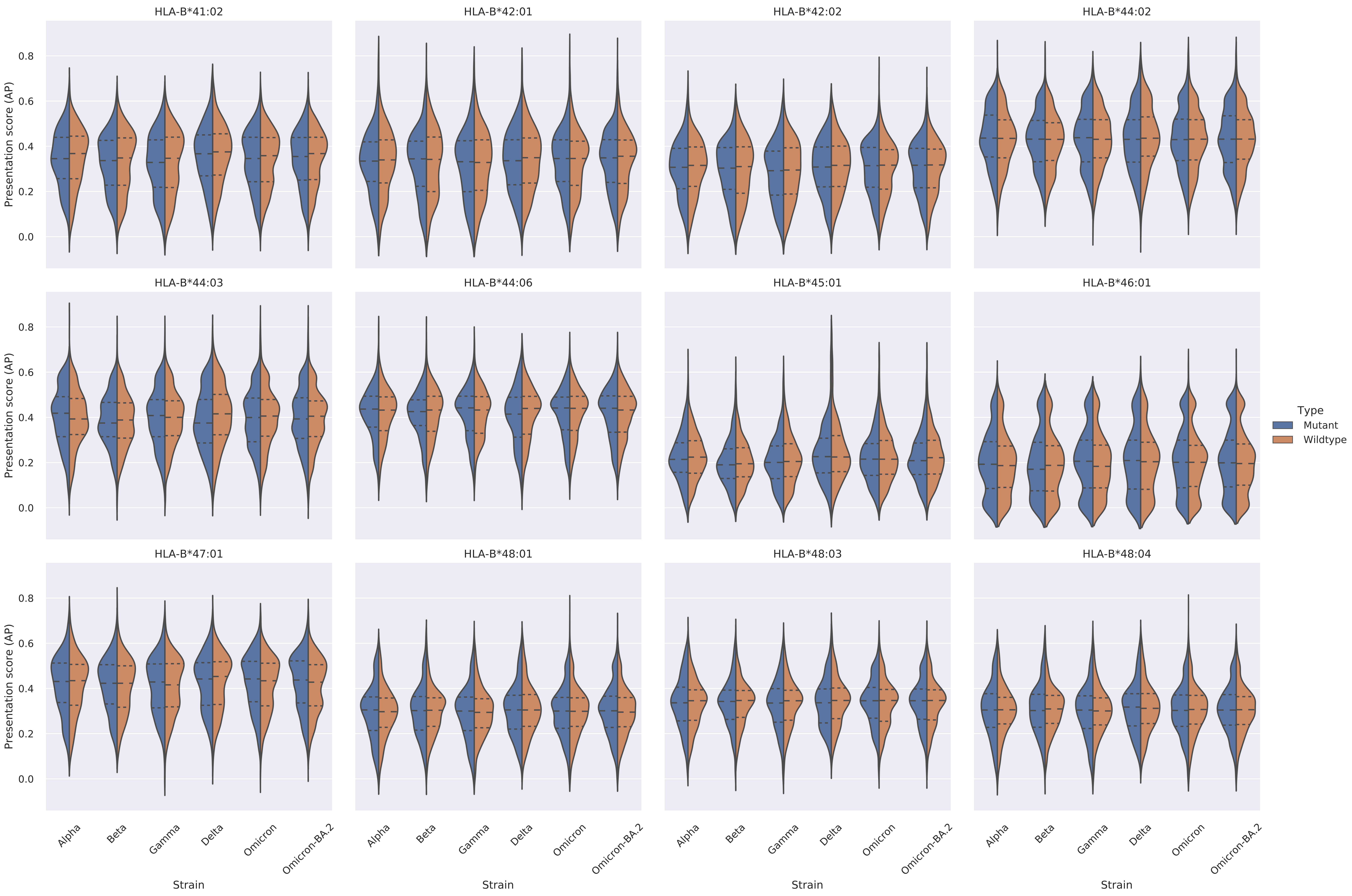

Supplementary Figure 4: Violin plots of the distribution of presentation scores (AP) for mutant and wildtype epitopes per allele for each VOC and Omicron-BA.2. The dashed lines within each violin represent the quartiles.

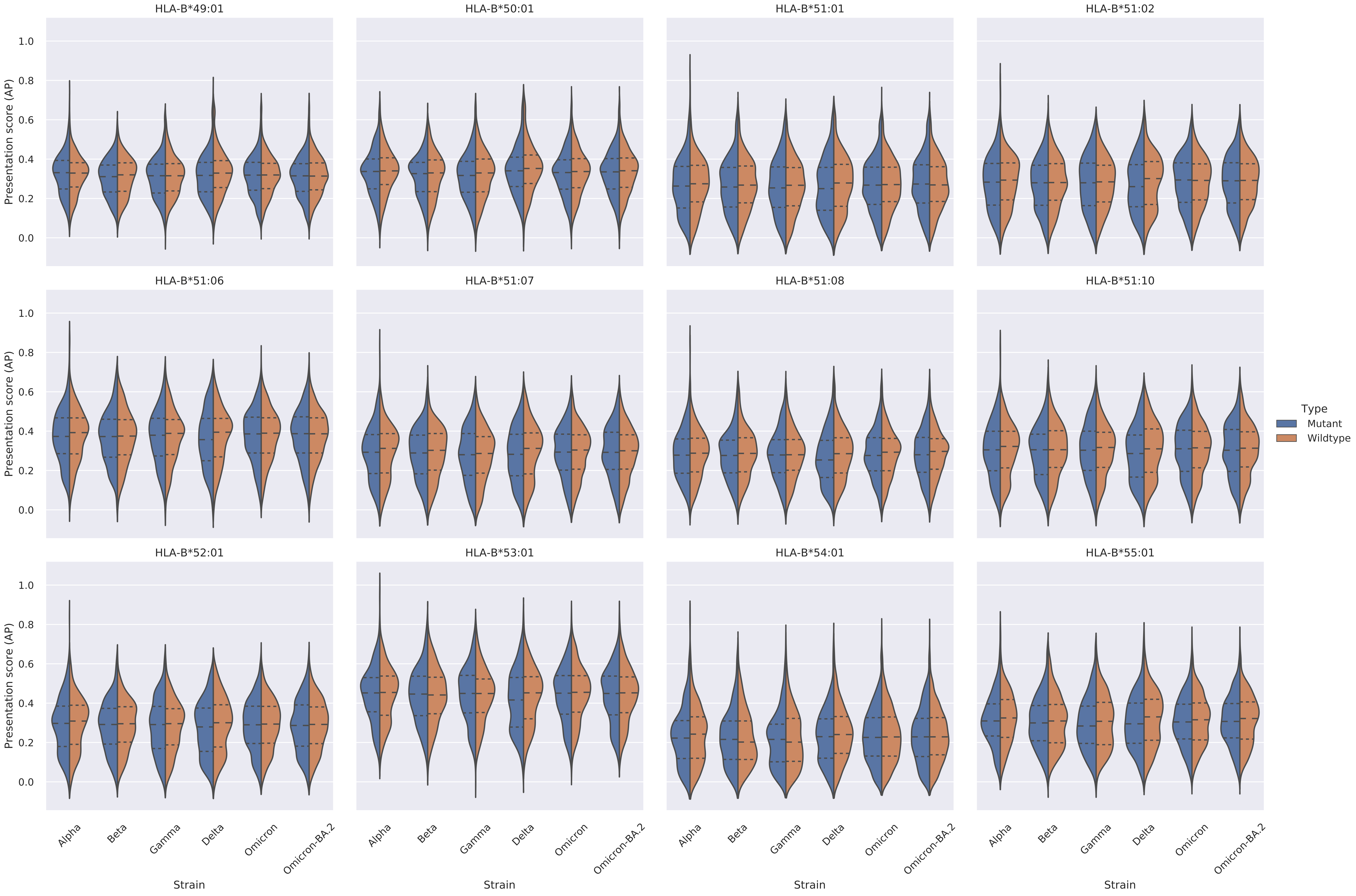

Supplementary Figure 4: Violin plots of the distribution of presentation scores (AP) for mutant and wildtype epitopes per allele for each VOC and Omicron-BA.2. The dashed lines within each violin represent the quartiles.

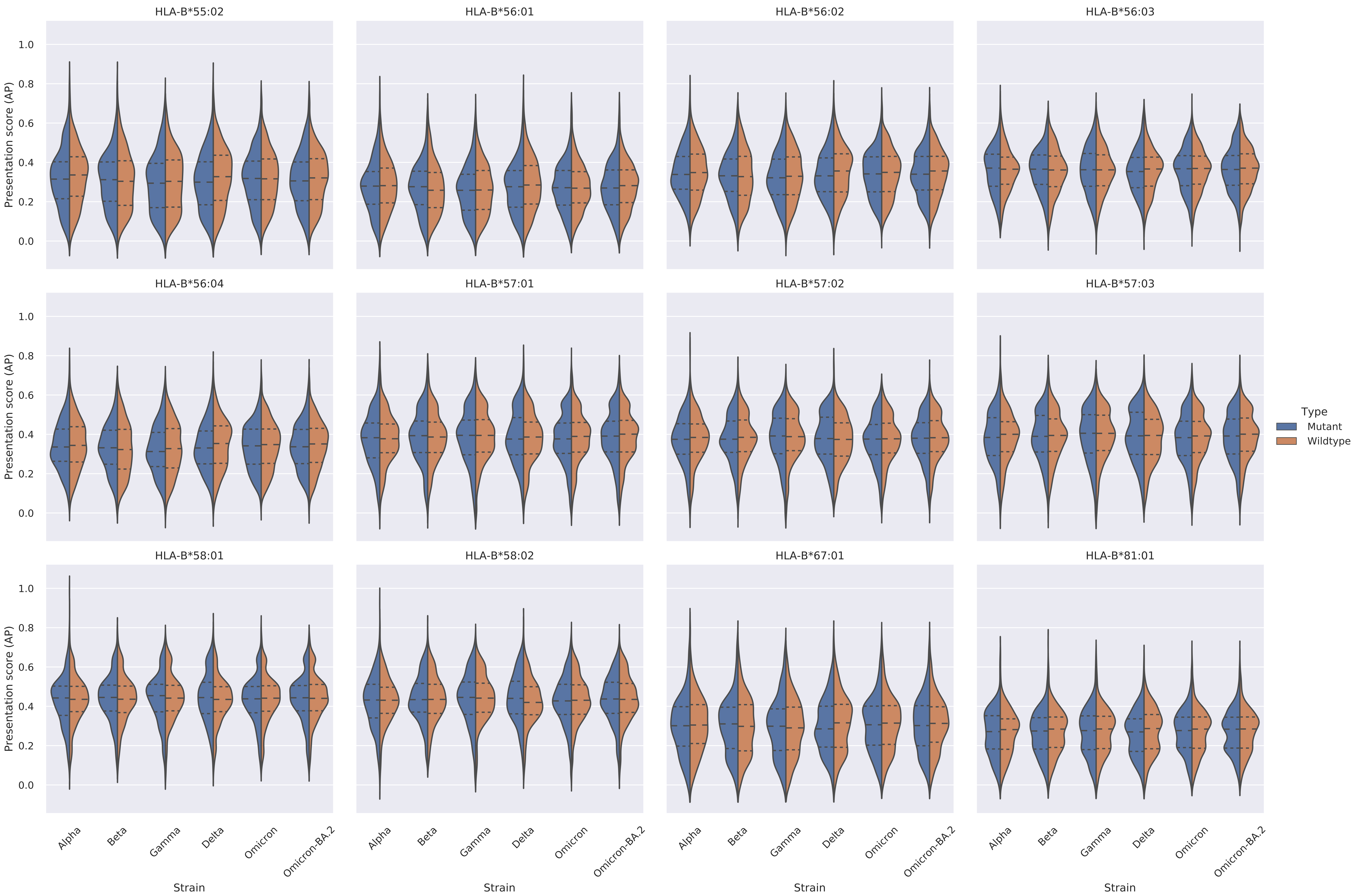

Supplementary Figure 5: Density of average AP scores > 0.5 at each mutation for a mutant (foreground) and wildtype (background) − protein E

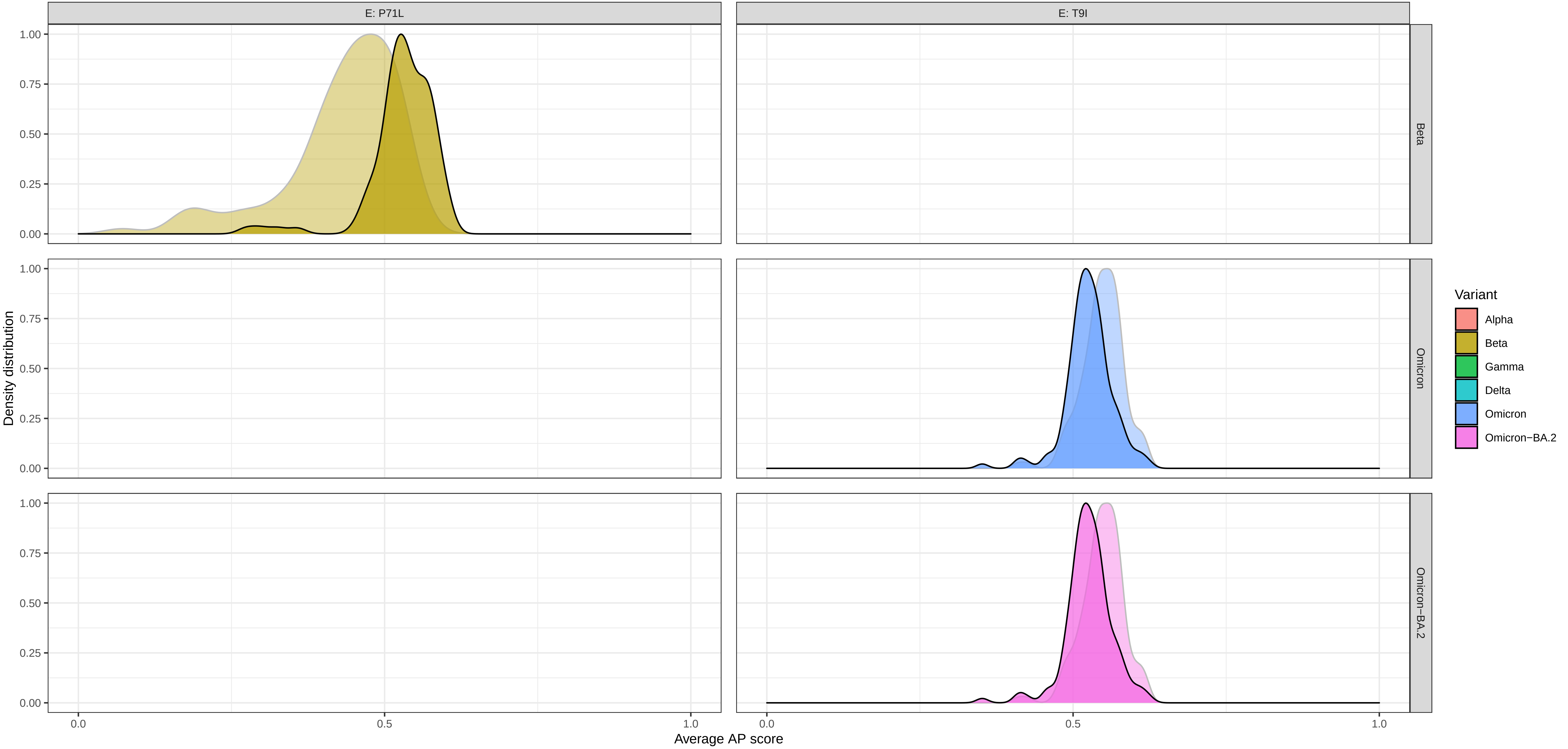

### Supplementary Figure 5: Density of average AP scores > 0.5 at each mutation for a mutant (foreground) and wildtype (background) − protein M

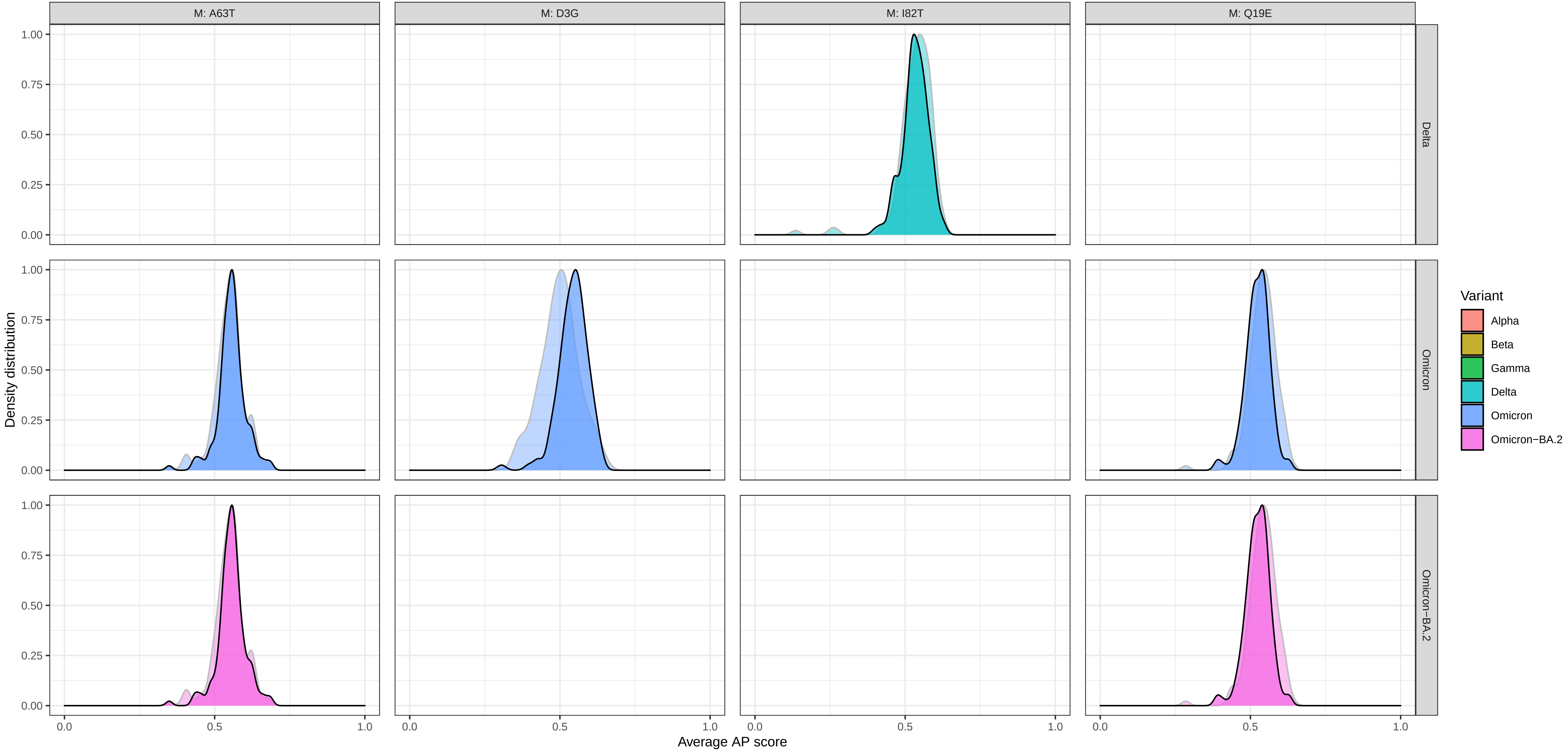

Supplementary Figure 5: Density of average AP scores > 0.5 at each mutation for a mutant (foreground) and wildtype (background) − protein N

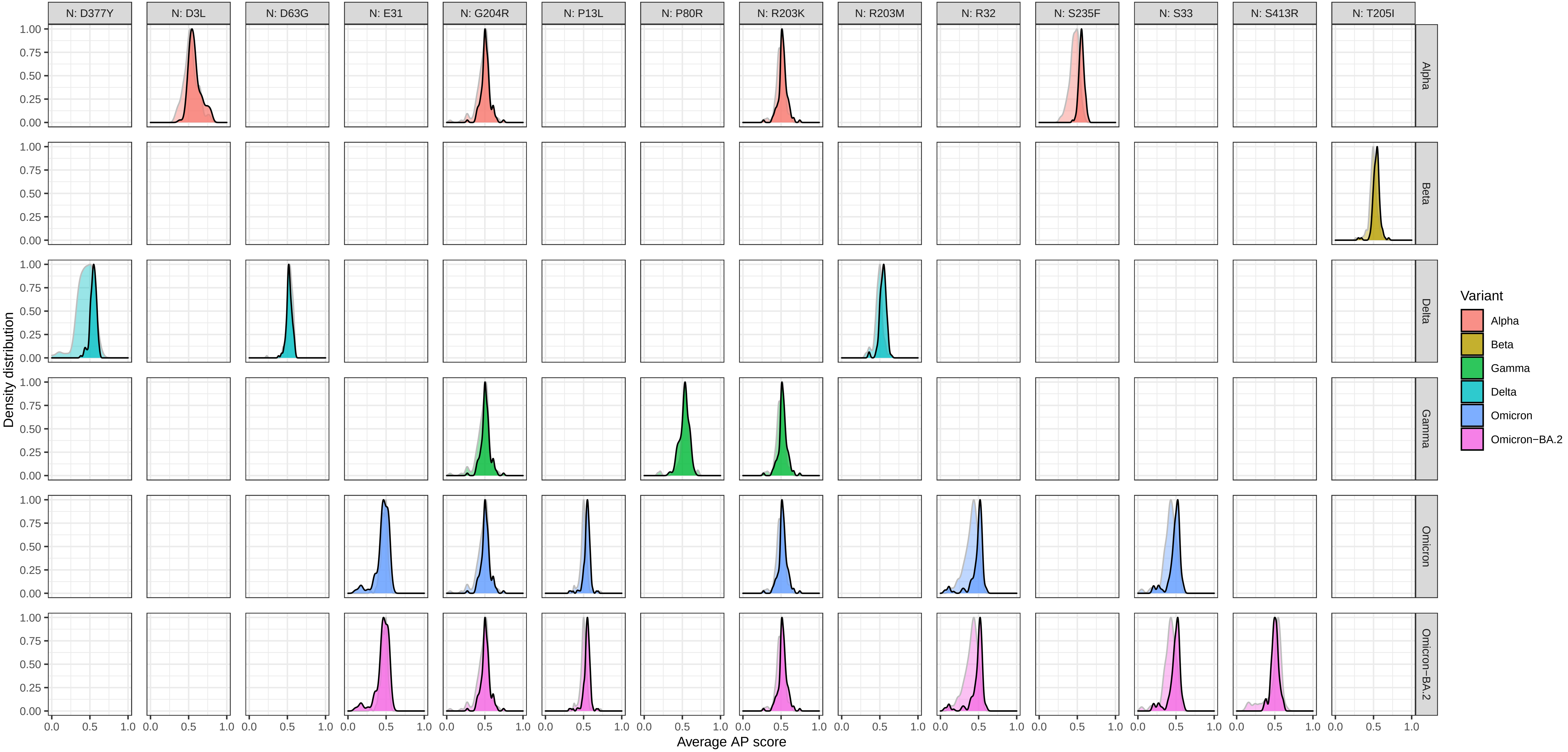

Supplementary Figure 5: Density of average AP scores > 0.5 at each mutation for a mutant (foreground) and wildtype (background) − protein ORF1ab

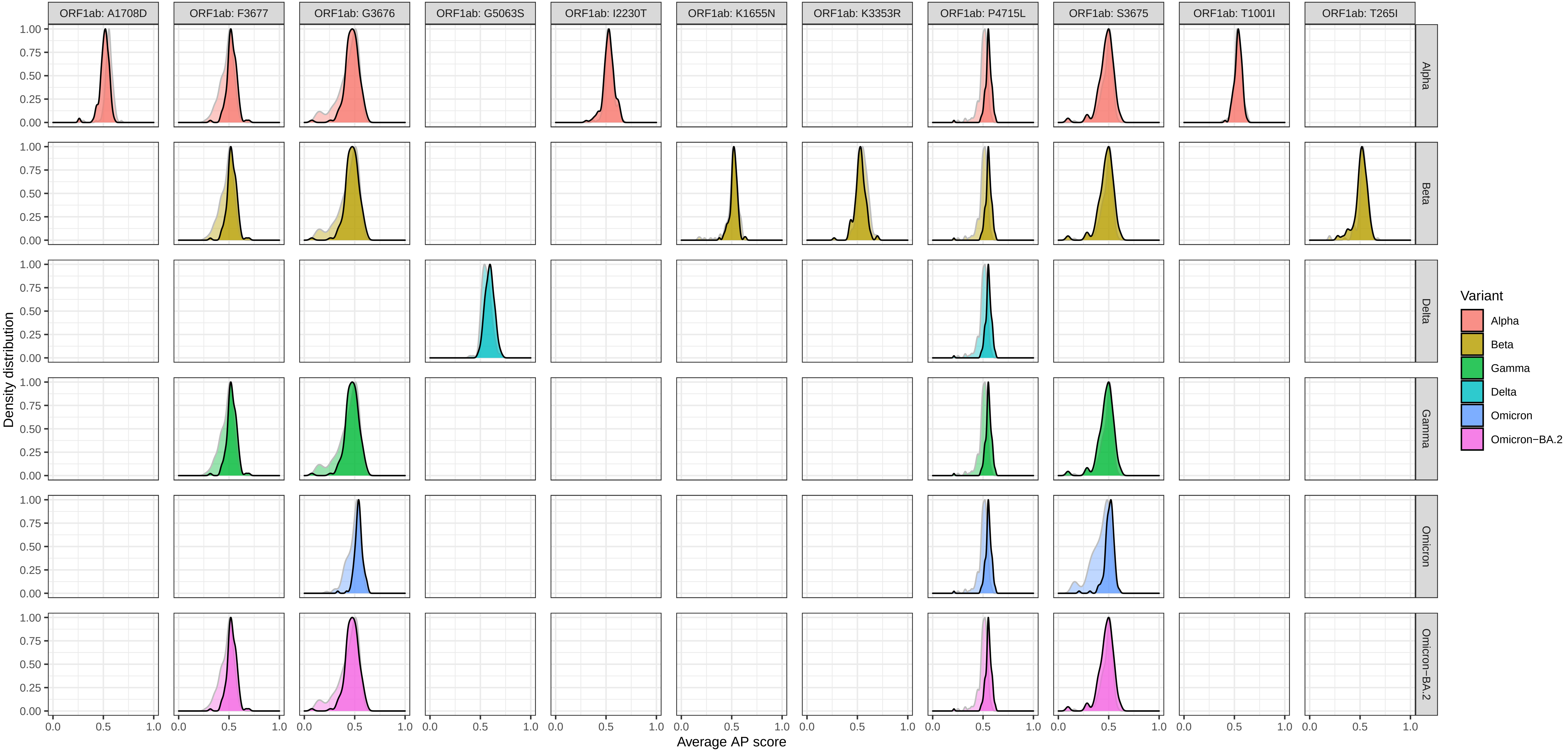

### Supplementary Figure 5: Density of average AP scores > 0.5 at each mutation for a mutant (foreground) and wildtype (background) − protein ORF1ab

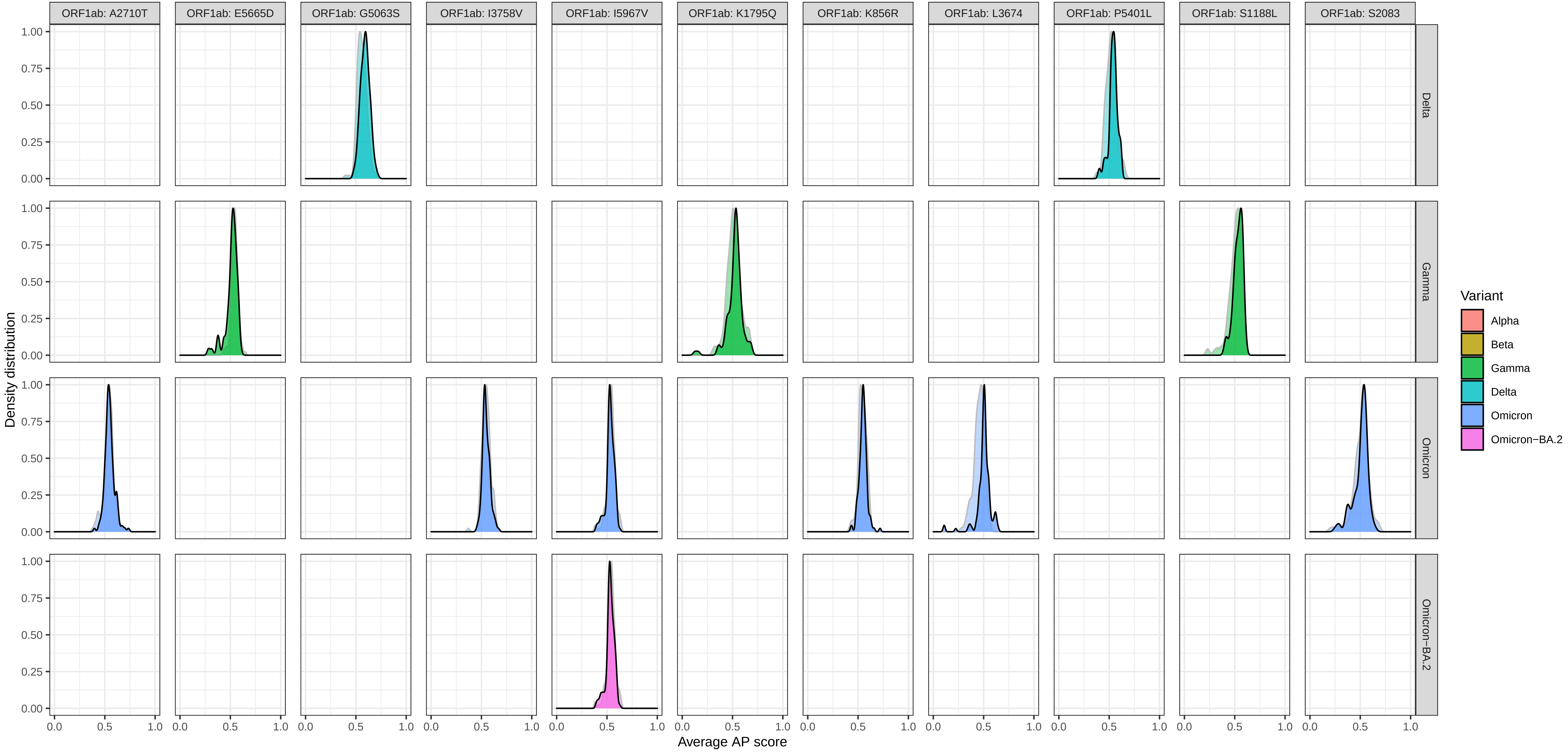

Supplementary Figure 5: Density of average AP scores > 0.5 at each mutation for a mutant (foreground) and wildtype (background) − protein ORF1ab

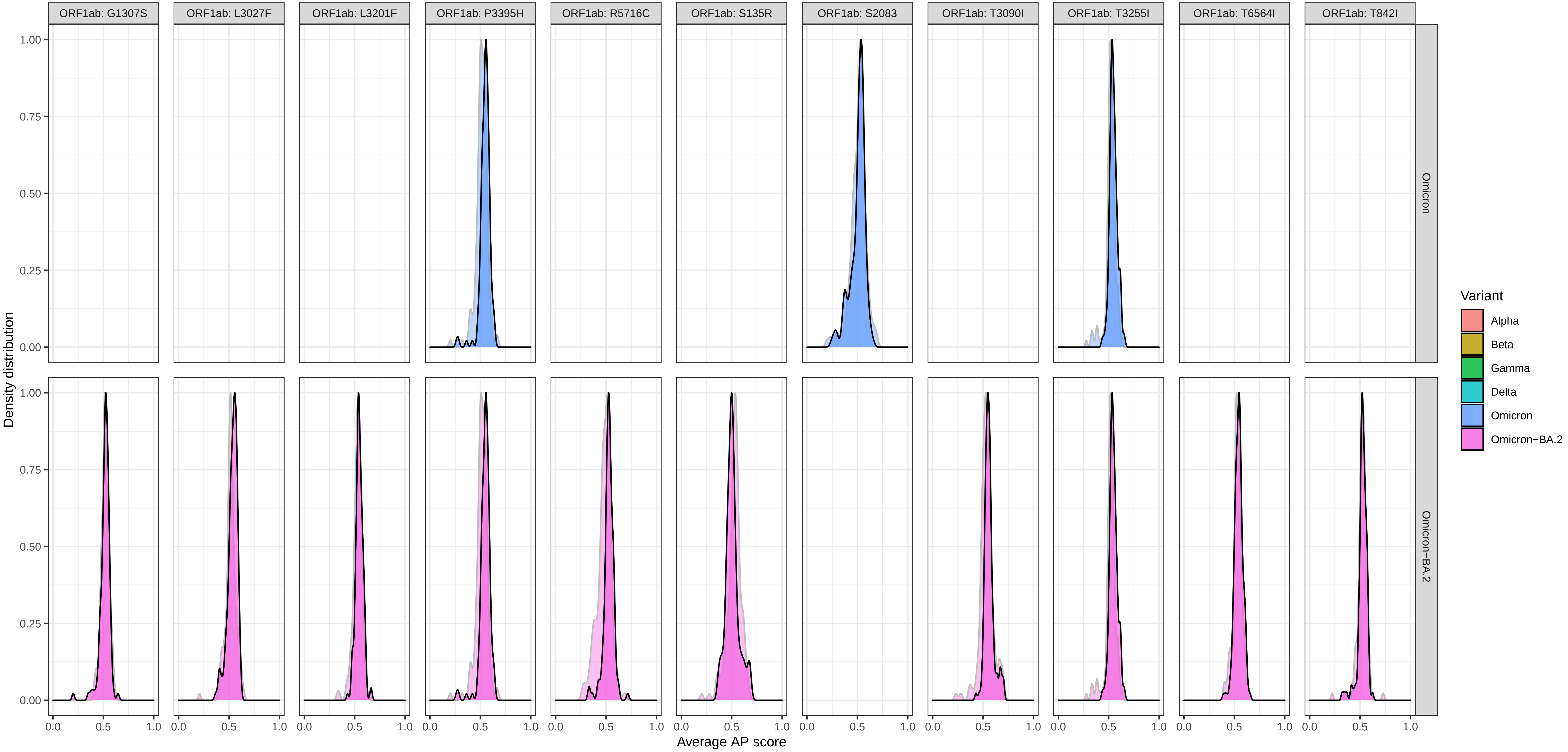

### Supplementary Figure 5: Density of average AP scores > 0.5 at each mutation for a mutant (foreground) and wildtype (background) − protein ORF3a

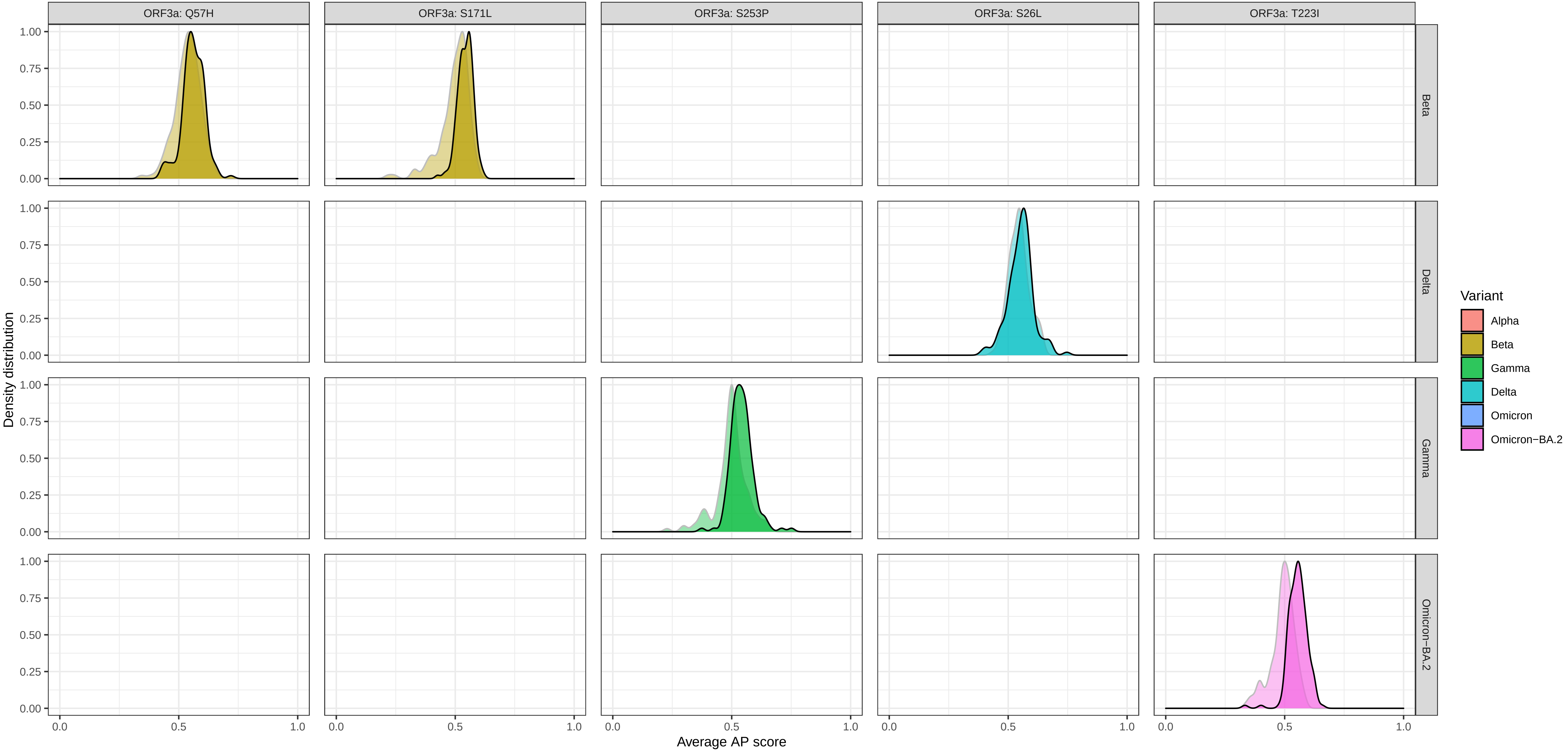

### Supplementary Figure 5: Density of average AP scores > 0.5 at each mutation for a mutant (foreground) and wildtype (background) − protein ORF6

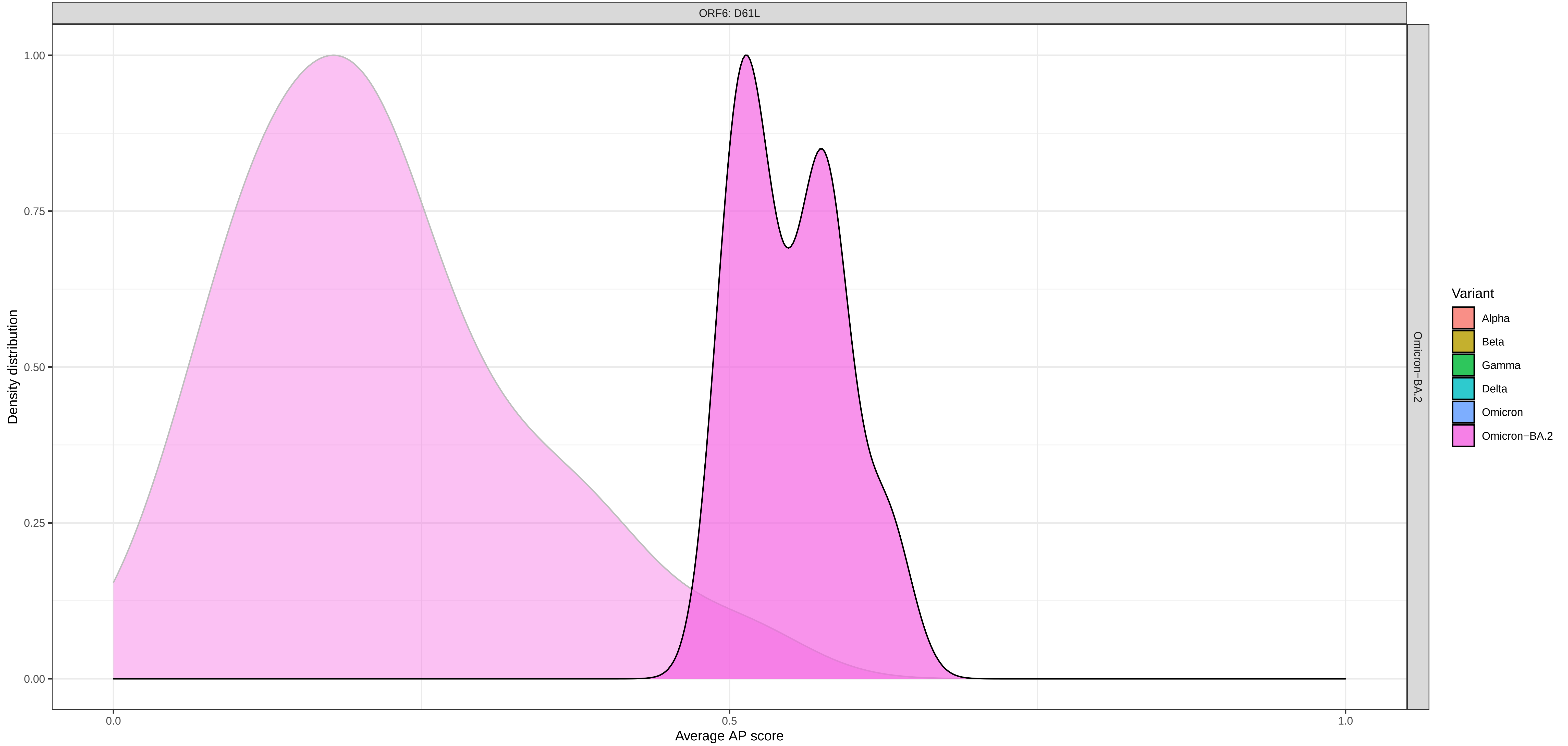

Supplementary Figure 5: Density of average AP scores > 0.5 at each mutation for a mutant (foreground) and wildtype (background) − protein ORF7a

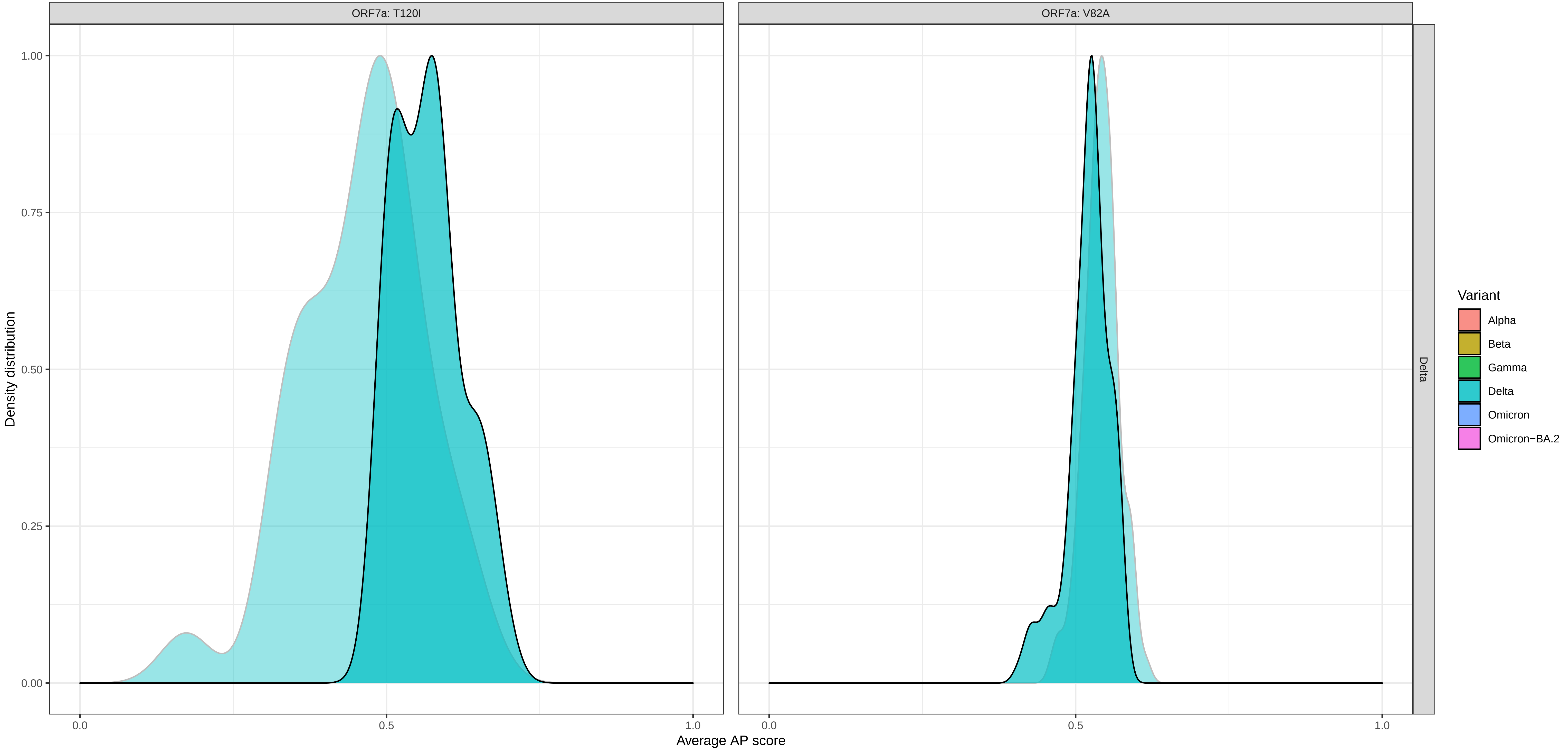

Supplementary Figure 5: Density of average AP scores > 0.5 at each mutation for a mutant (foreground) and wildtype (background) − protein ORF8

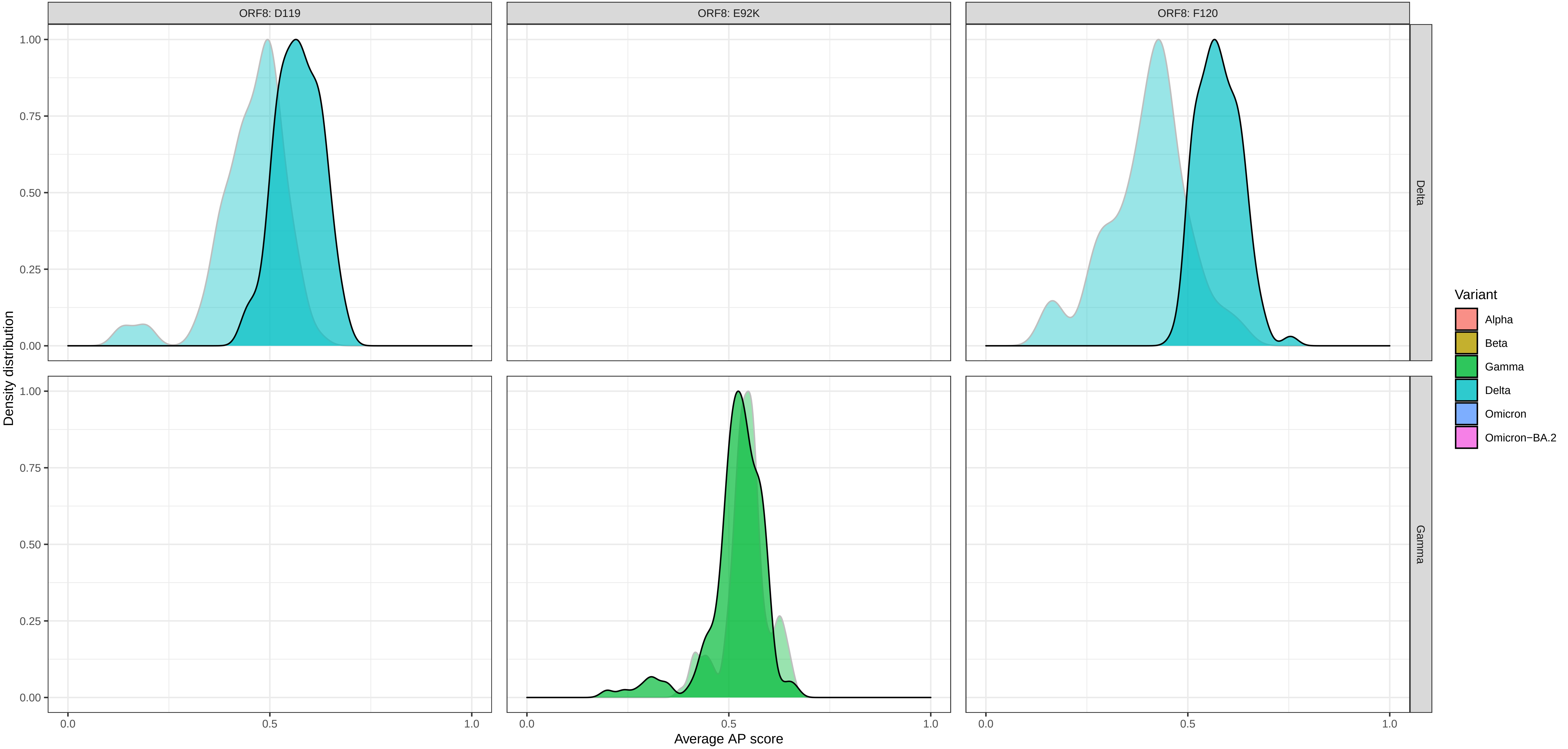

### Supplementary Figure 5: Density of average AP scores > 0.5 at each mutation for a mutant (foreground) and wildtype (background) − protein S

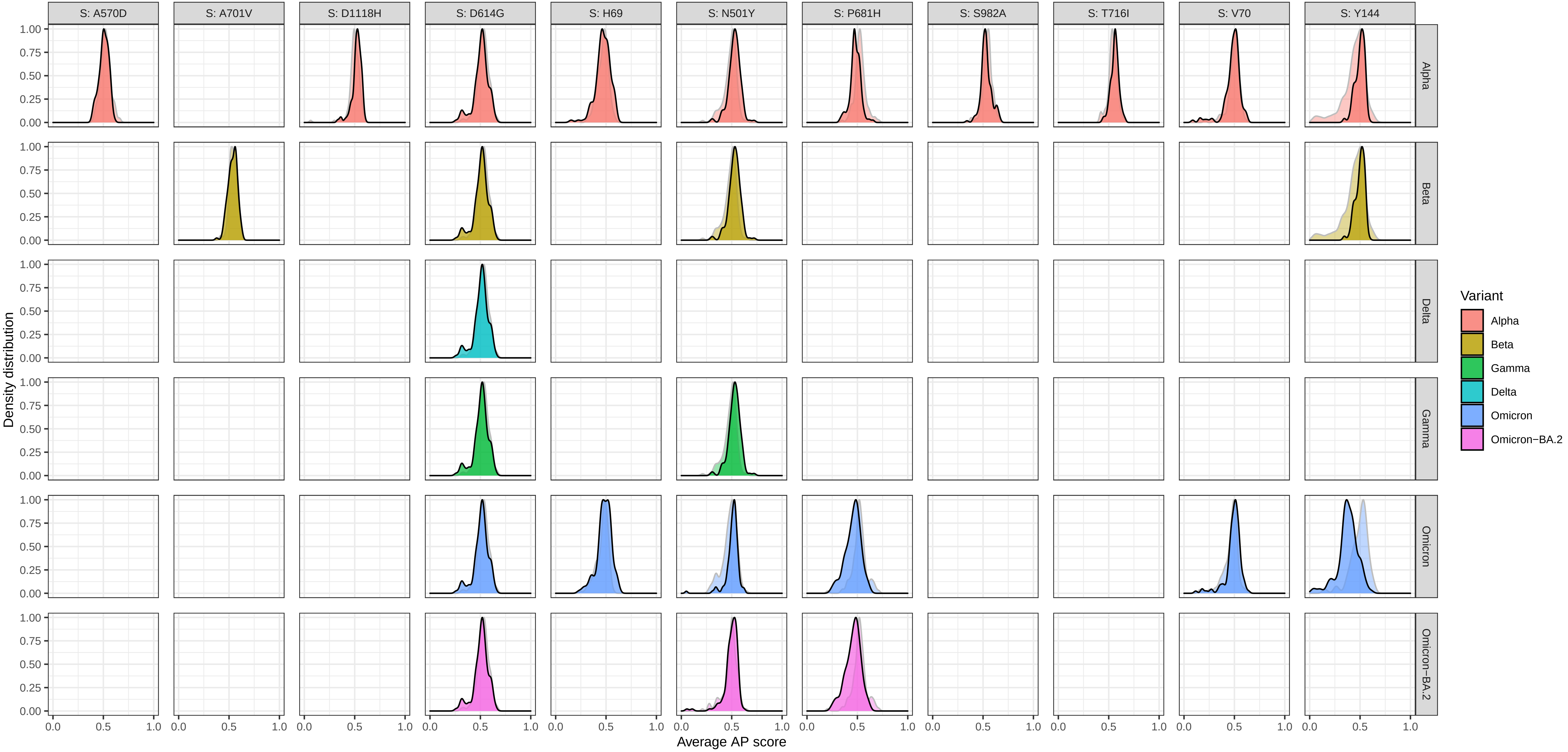

### Supplementary Figure 5: Density of average AP scores > 0.5 at each mutation for a mutant (foreground) and wildtype (background) − protein S

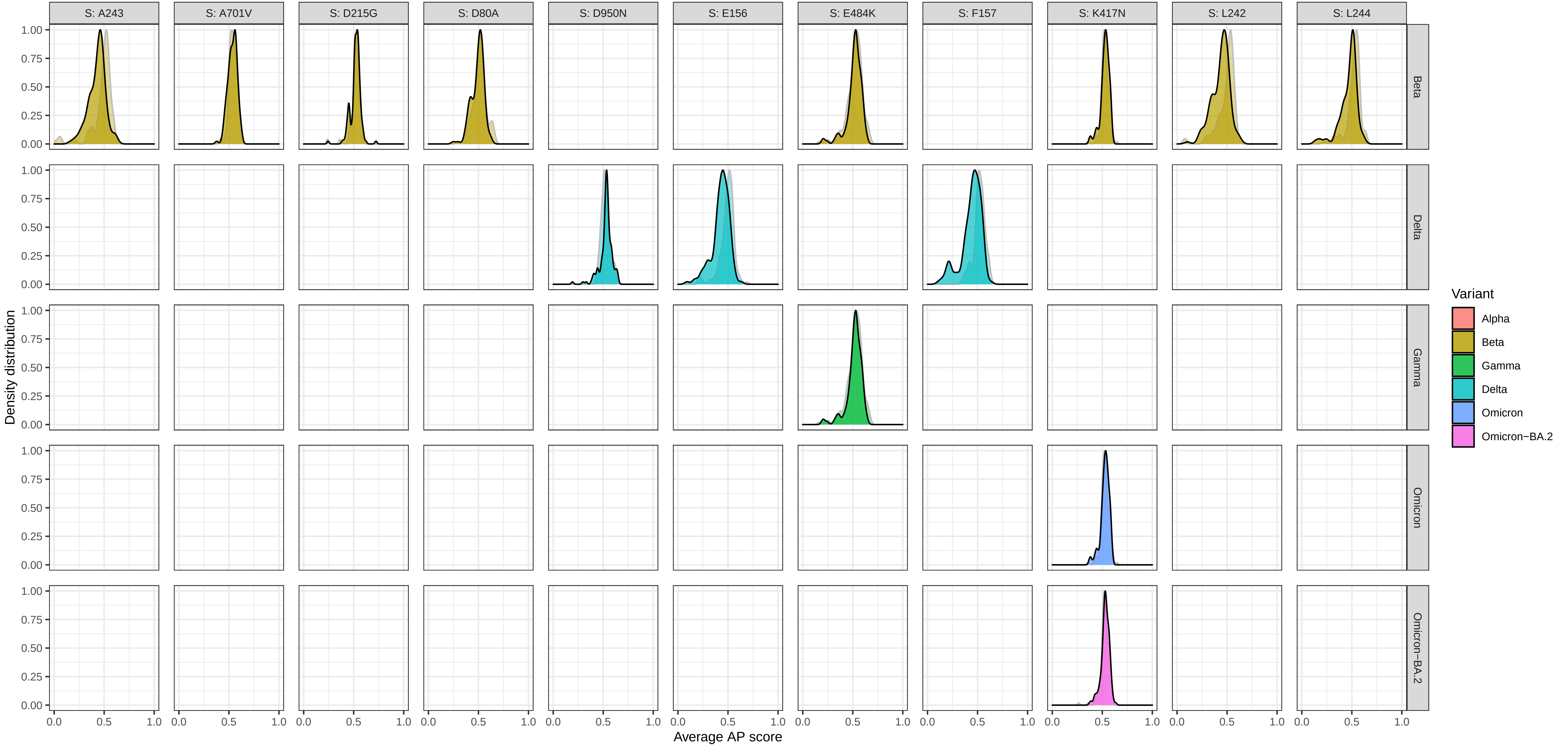

Supplementary Figure 5: Density of average AP scores > 0.5 at each mutation for a mutant (foreground) and wildtype (background) − protein S

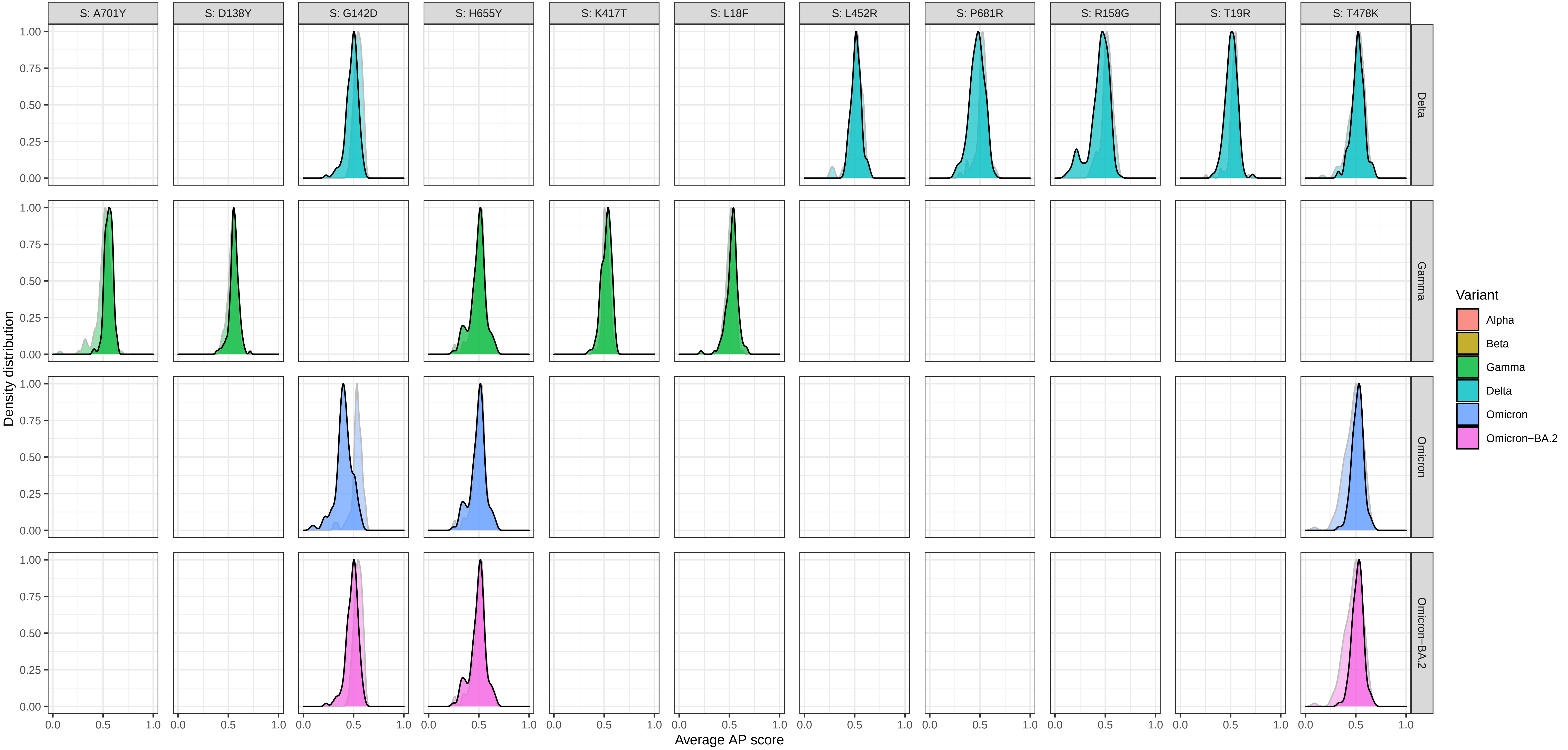

### Supplementary Figure 5: Density of average AP scores > 0.5 at each mutation for a mutant (foreground) and wildtype (background) − protein S

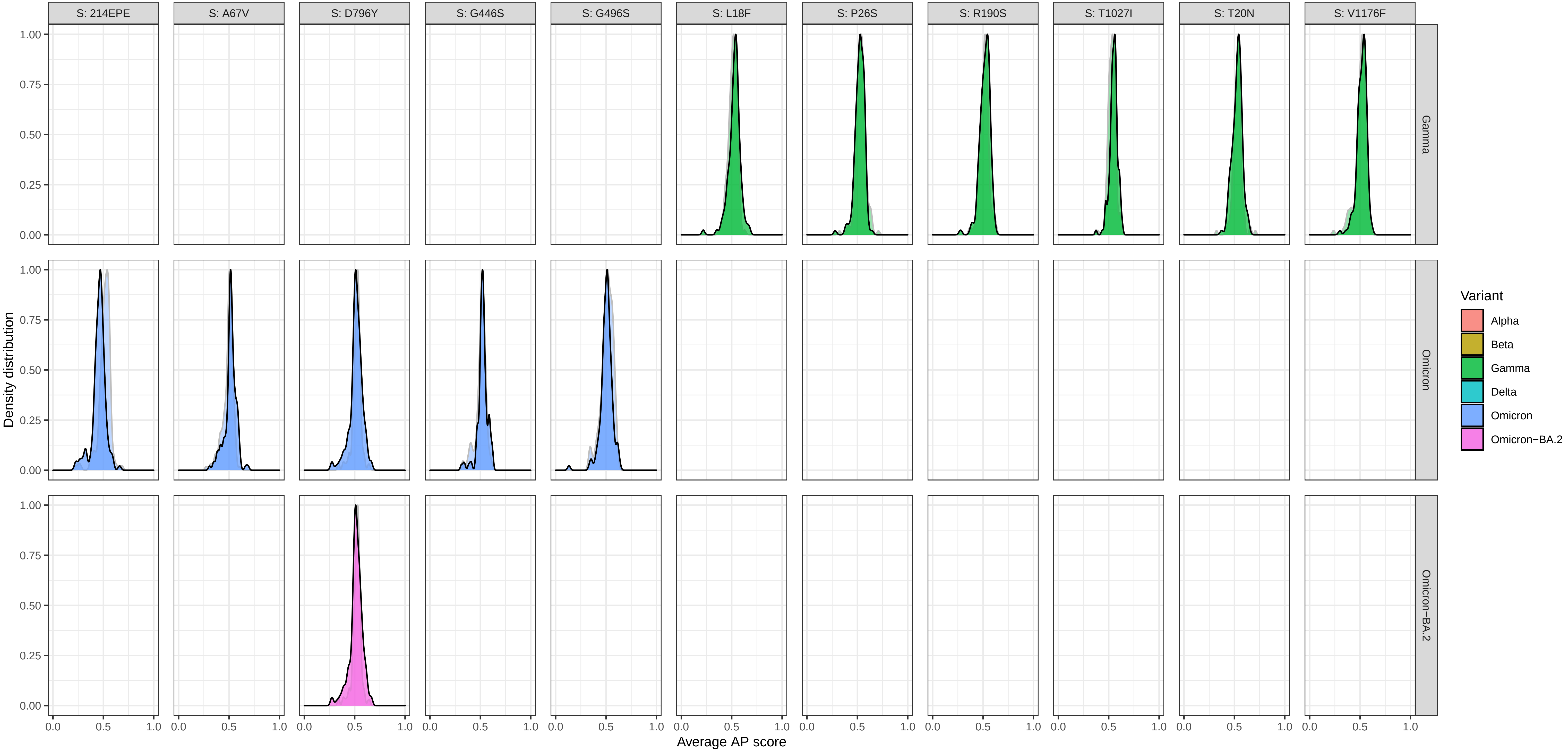

Supplementary Figure 5: Density of average AP scores > 0.5 at each mutation for a mutant (foreground) and wildtype (background) − protein S

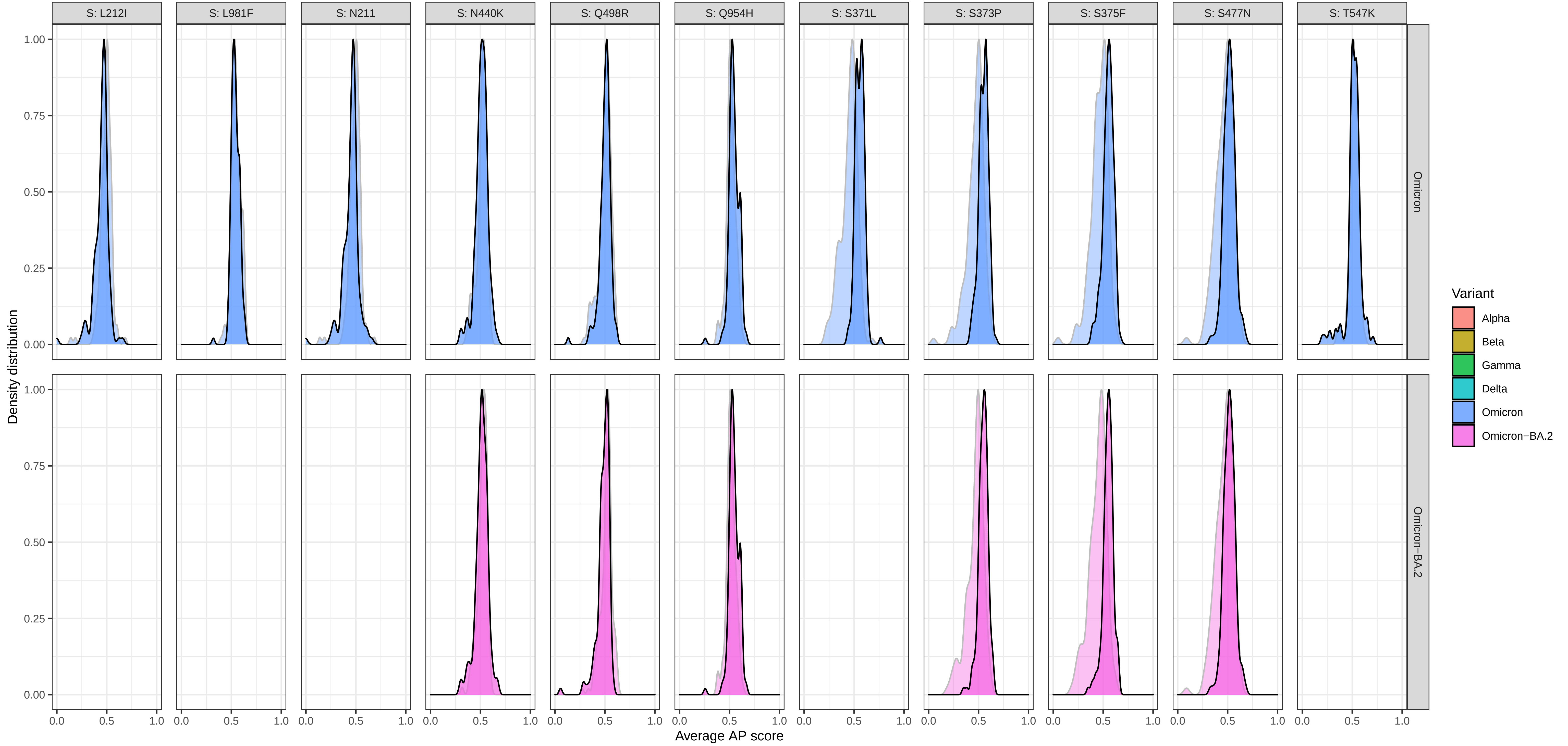

Supplementary Figure 5: Density of average AP scores > 0.5 at each mutation for a mutant (foreground) and wildtype (background) − protein S

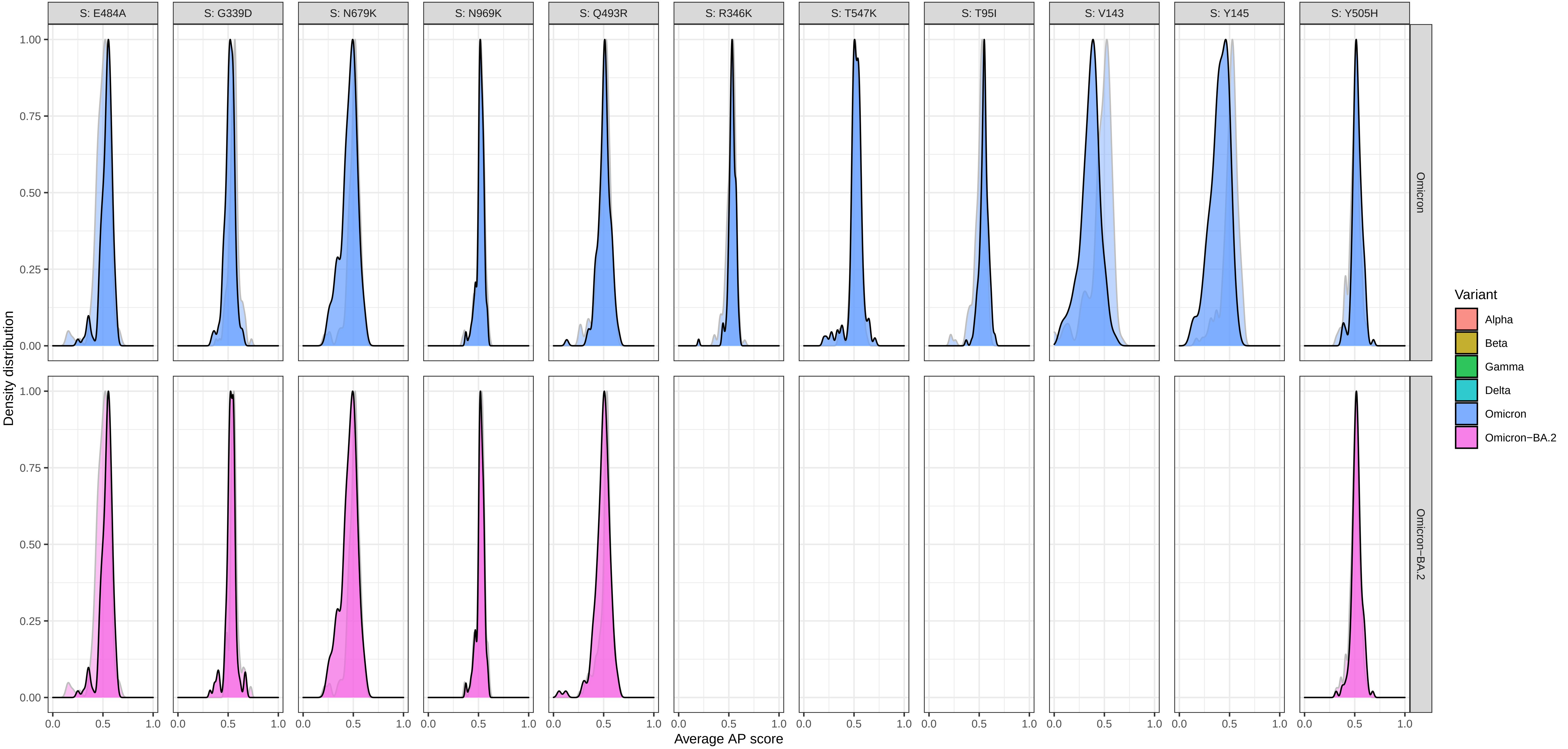

Supplementary Figure 5: Density of average AP scores > 0.5 at each mutation for a mutant (foreground) and wildtype (background) − protein S

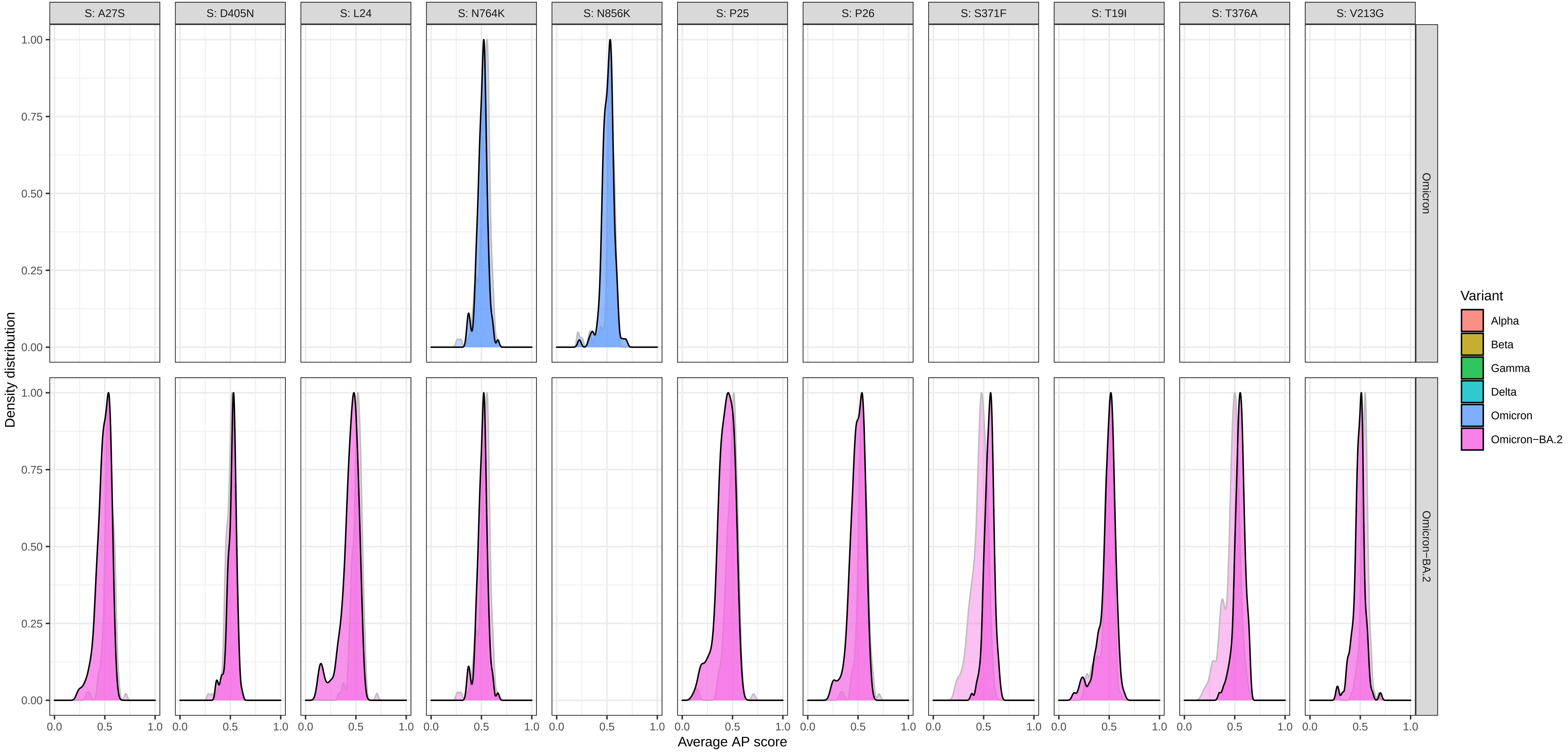

### Supplementary Figure 6: Density of average AP scores at each mutation for a mutant (foreground) and wildtype (background) − protein E

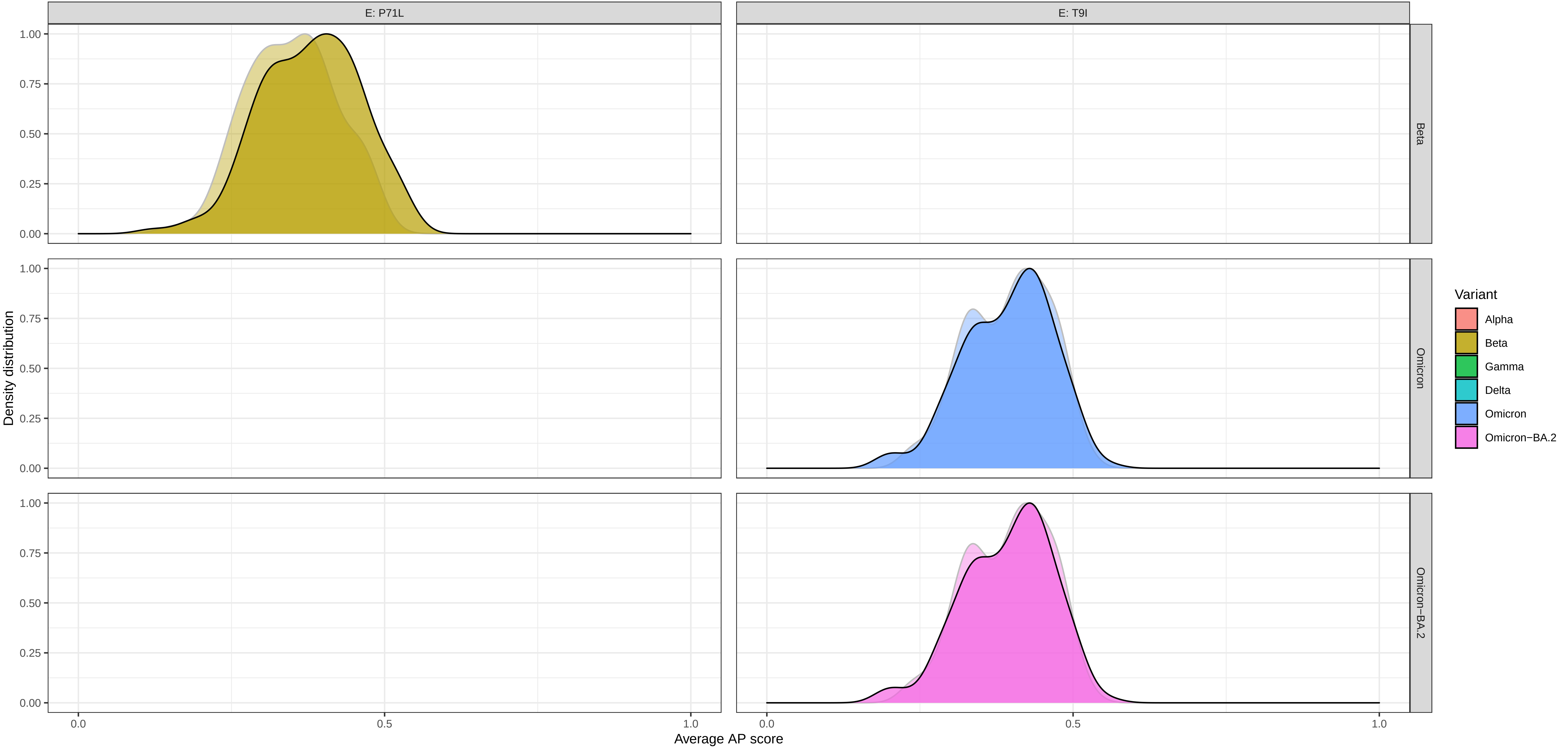

Supplementary Figure 6: Density of average AP scores at each mutation for a mutant (foreground) and wildtype (background) − protein M

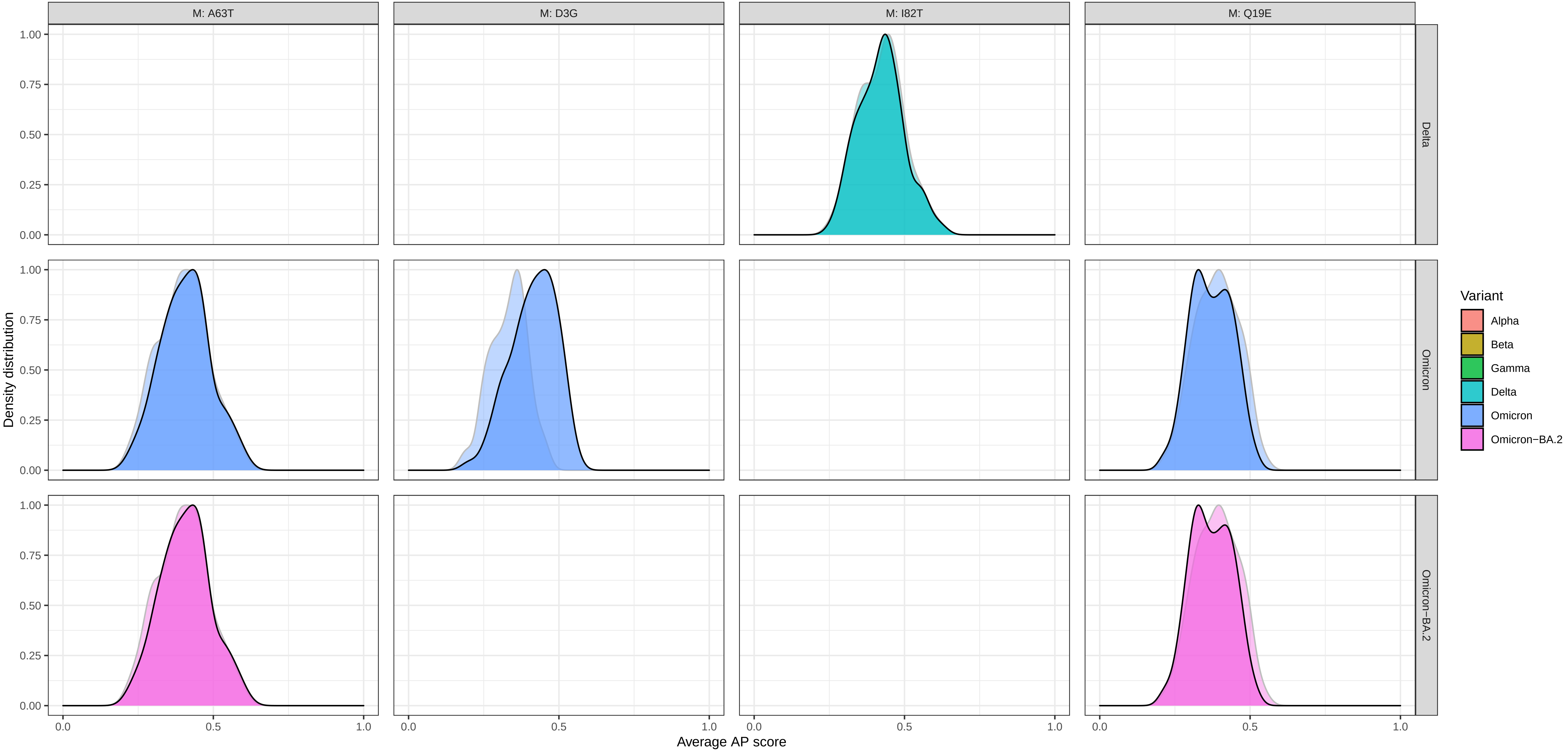

Supplementary Figure 6: Density of average AP scores at each mutation for a mutant (foreground) and wildtype (background) − protein N

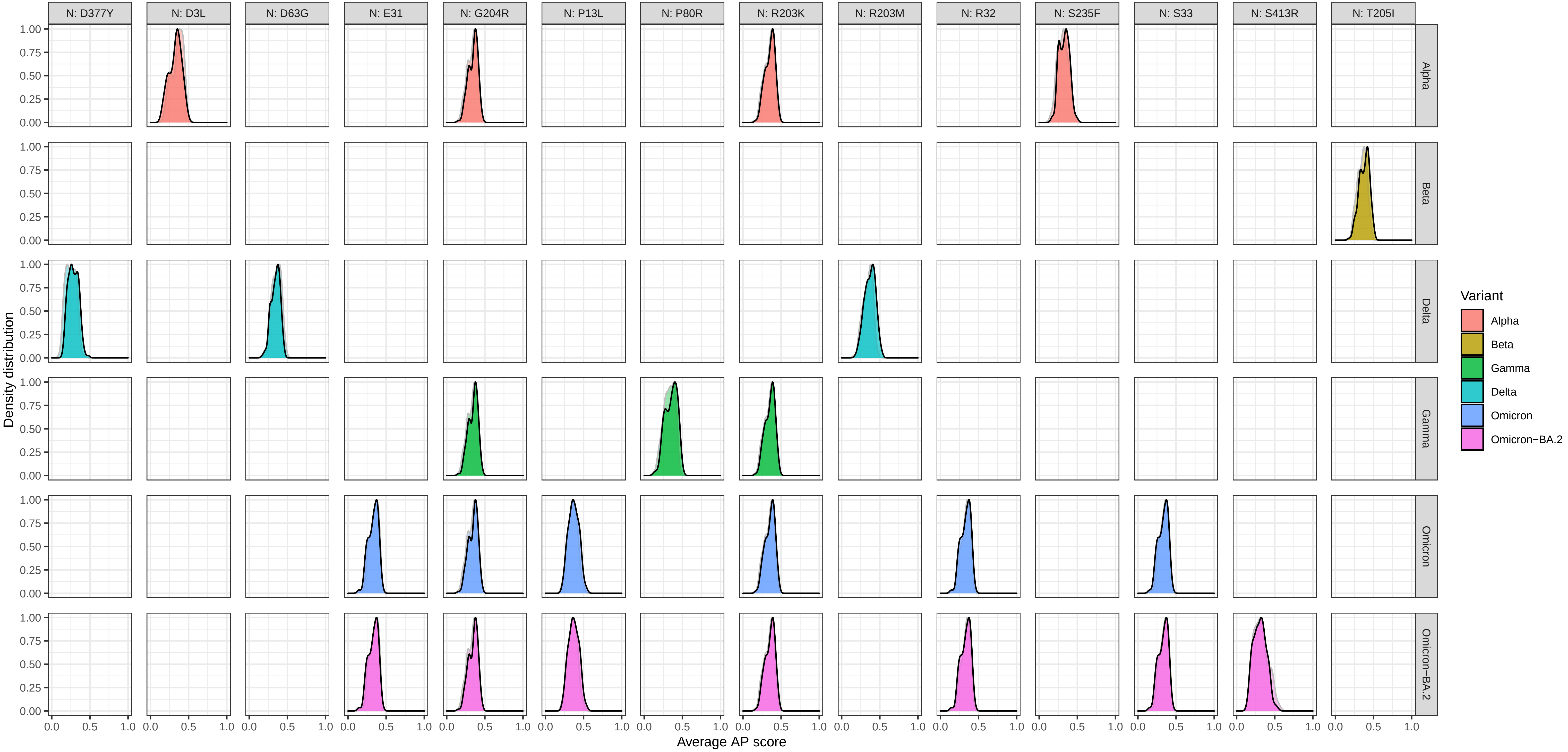

### Supplementary Figure 6: Density of average AP scores at each mutation for a mutant (foreground) and wildtype (background) − protein ORF1ab

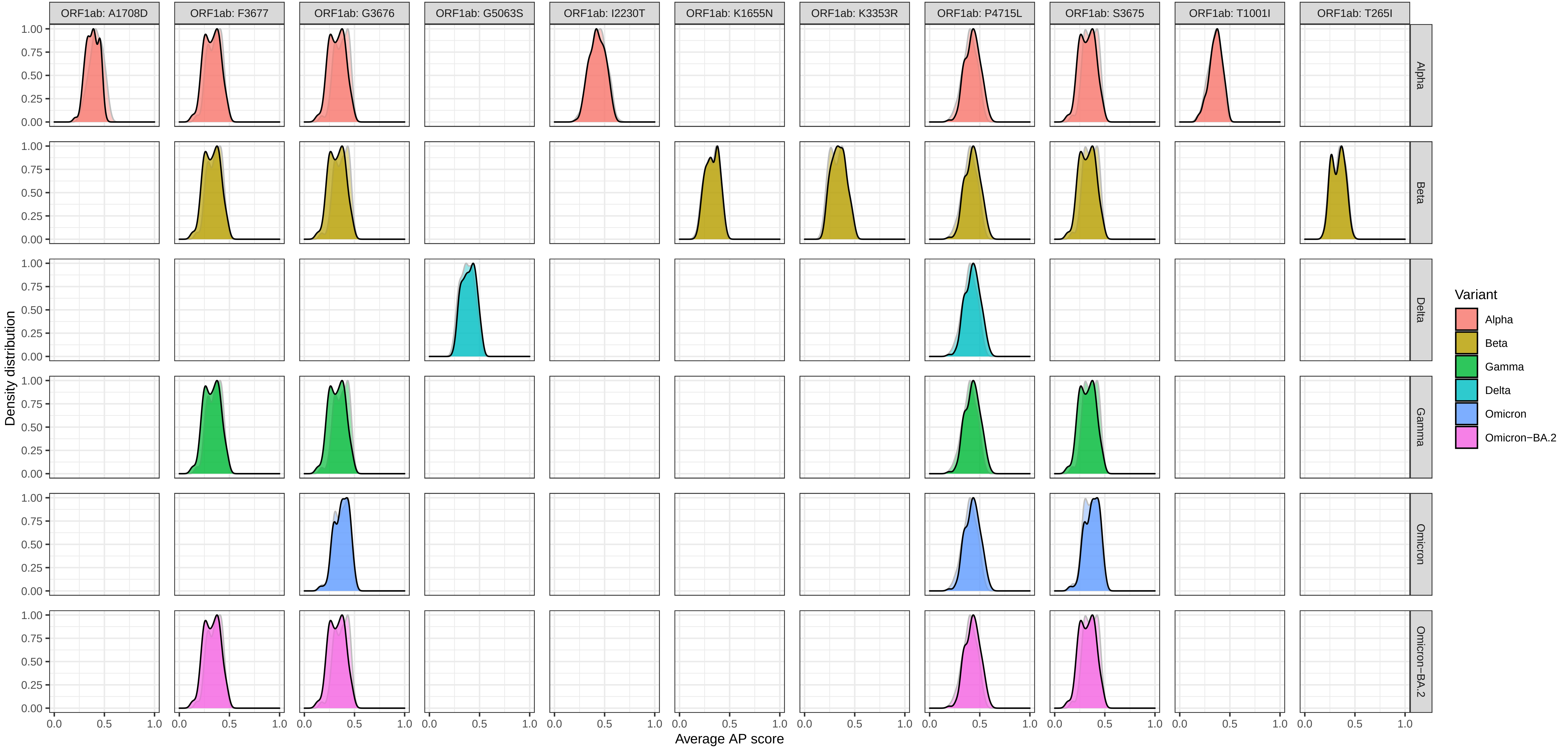

Supplementary Figure 6: Density of average AP scores at each mutation for a mutant (foreground) and wildtype (background) − protein ORF1ab

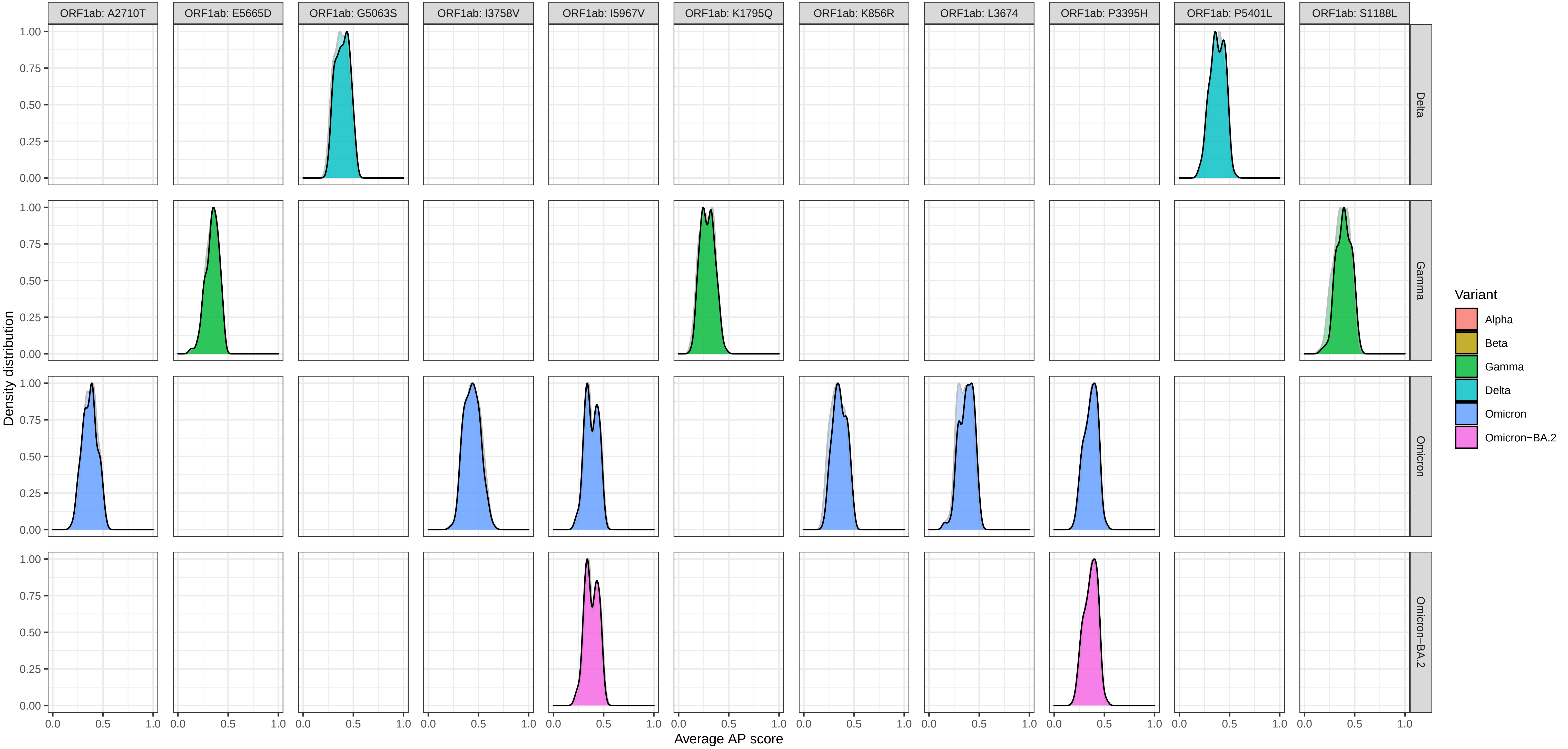

### Supplementary Figure 6: Density of average AP scores at each mutation for a mutant (foreground) and wildtype (background) − protein ORF1ab

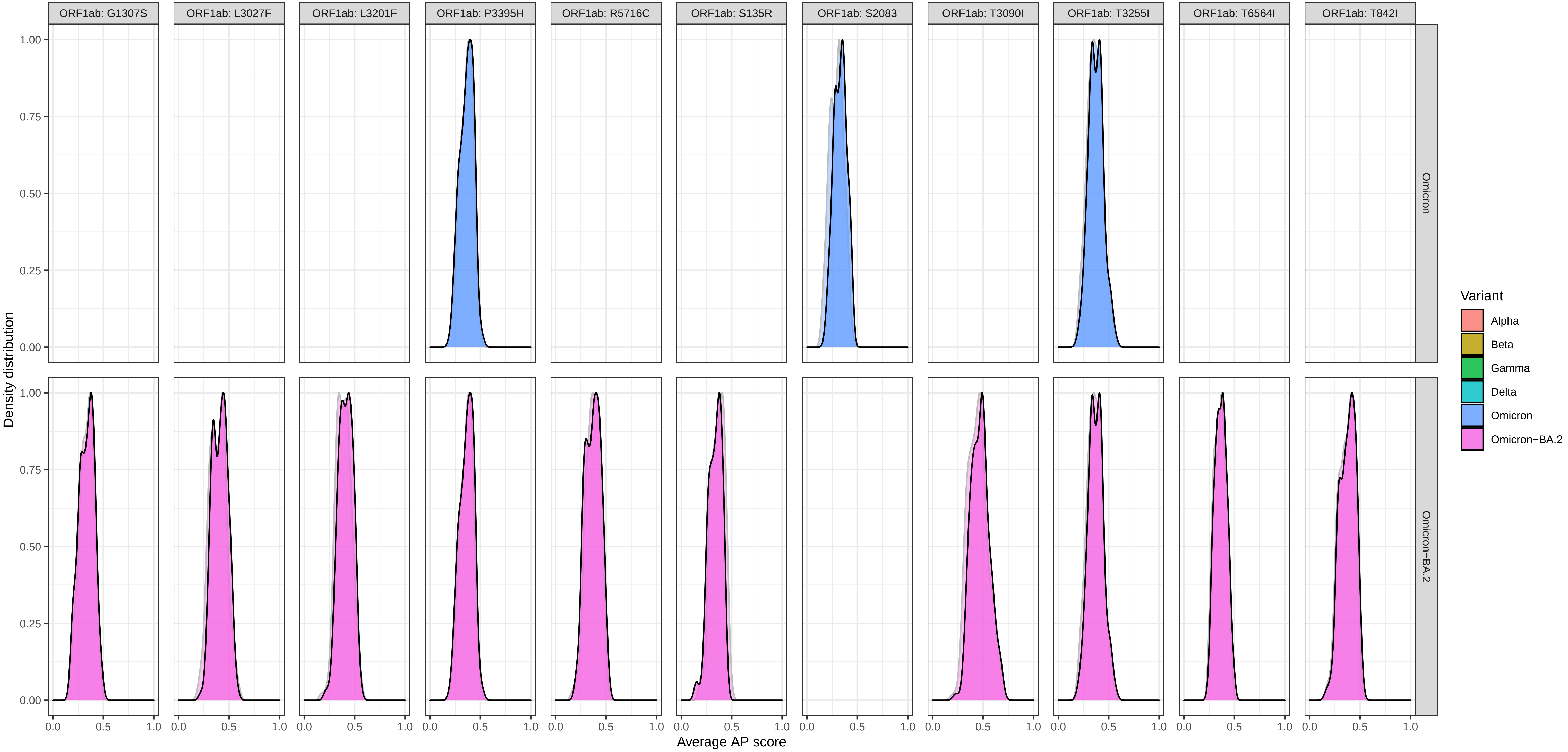

### Supplementary Figure 6: Density of average AP scores at each mutation for a mutant (foreground) and wildtype (background) − protein ORF3a

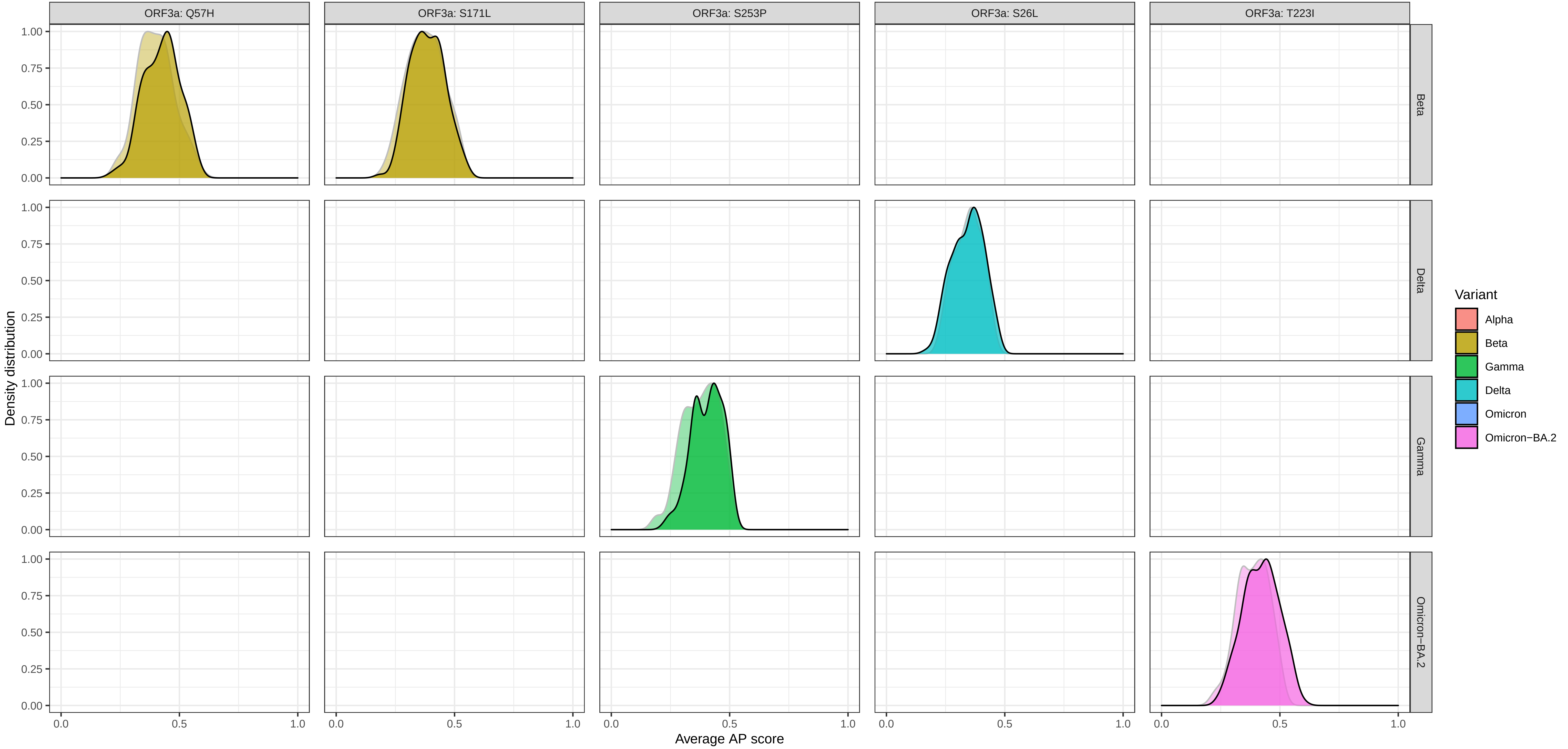

Supplementary Figure 6: Density of average AP scores at each mutation for a mutant (foreground) and wildtype (background) − protein ORF6

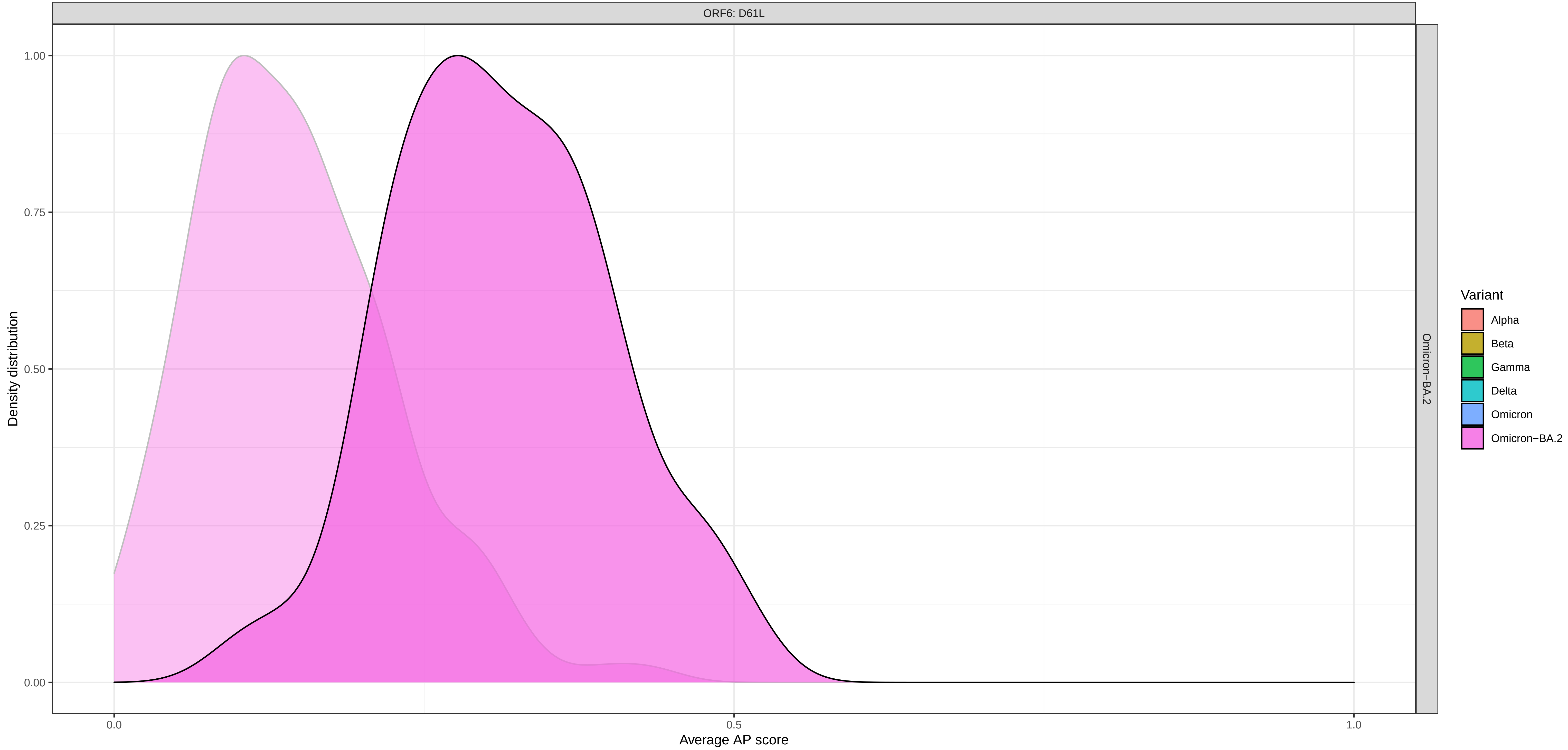

### Supplementary Figure 6: Density of average AP scores at each mutation for a mutant (foreground) and wildtype (background) − protein ORF7a

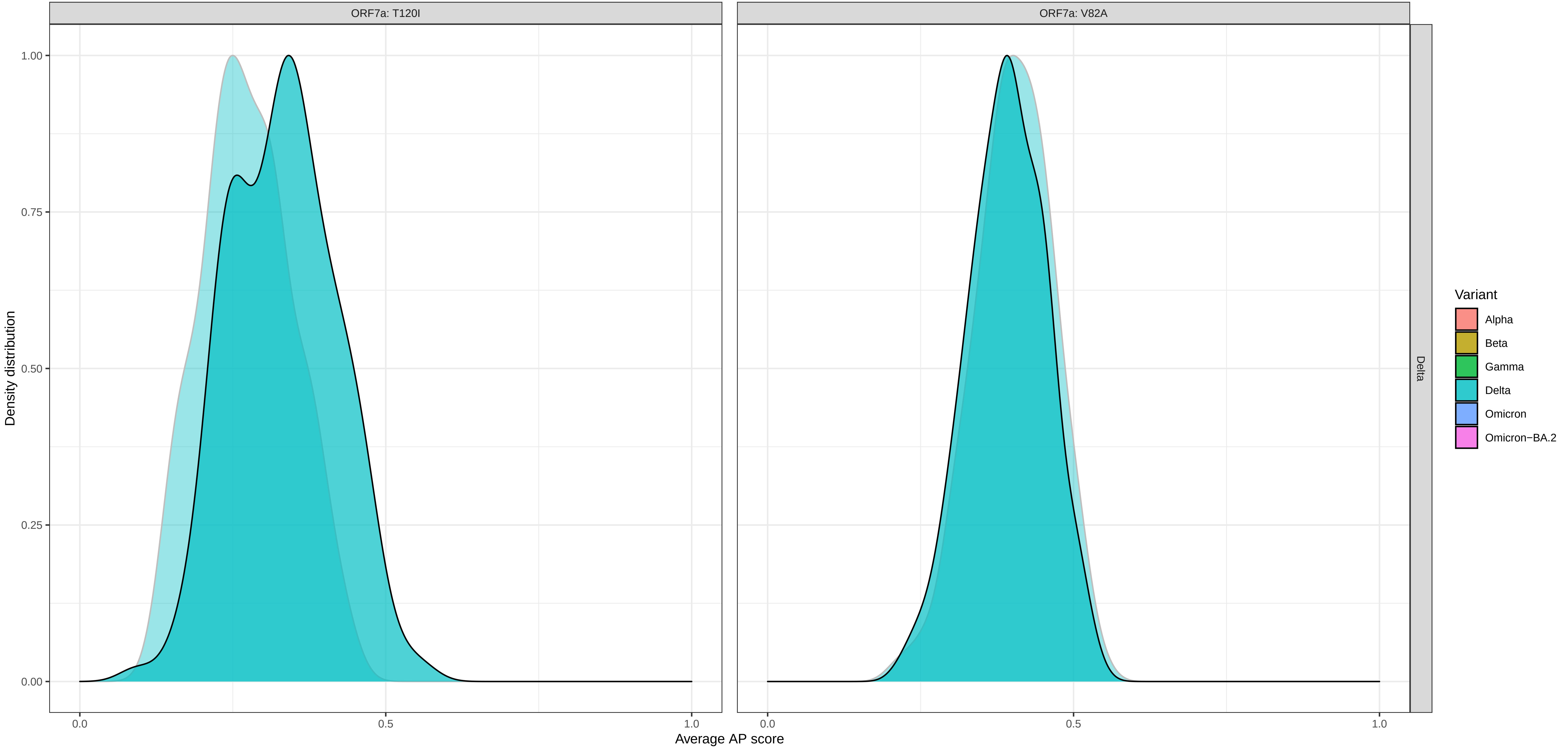

### Supplementary Figure 6: Density of average AP scores at each mutation for a mutant (foreground) and wildtype (background) − protein ORF8

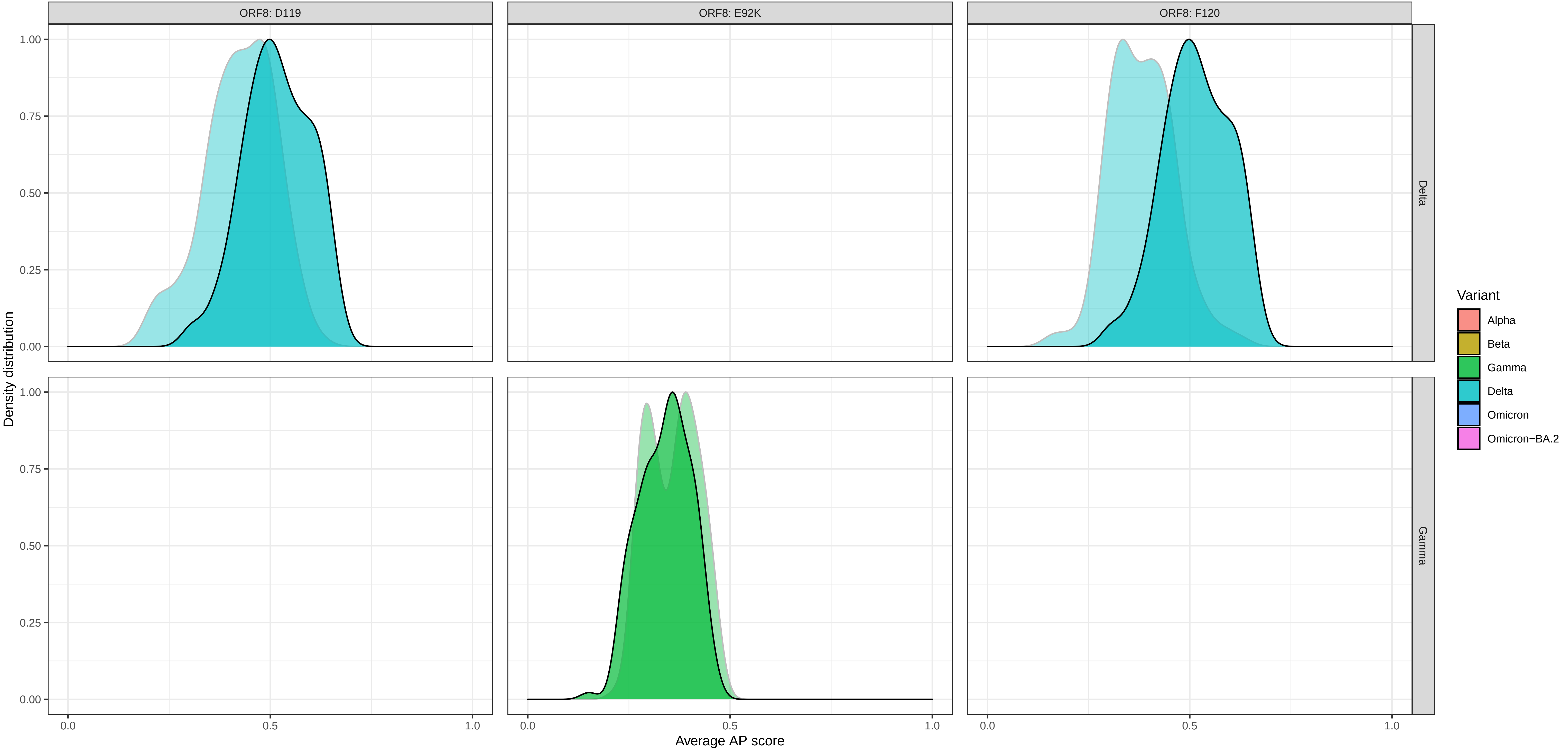

Supplementary Figure 6: Density of average AP scores at each mutation for a mutant (foreground) and wildtype (background) − protein S

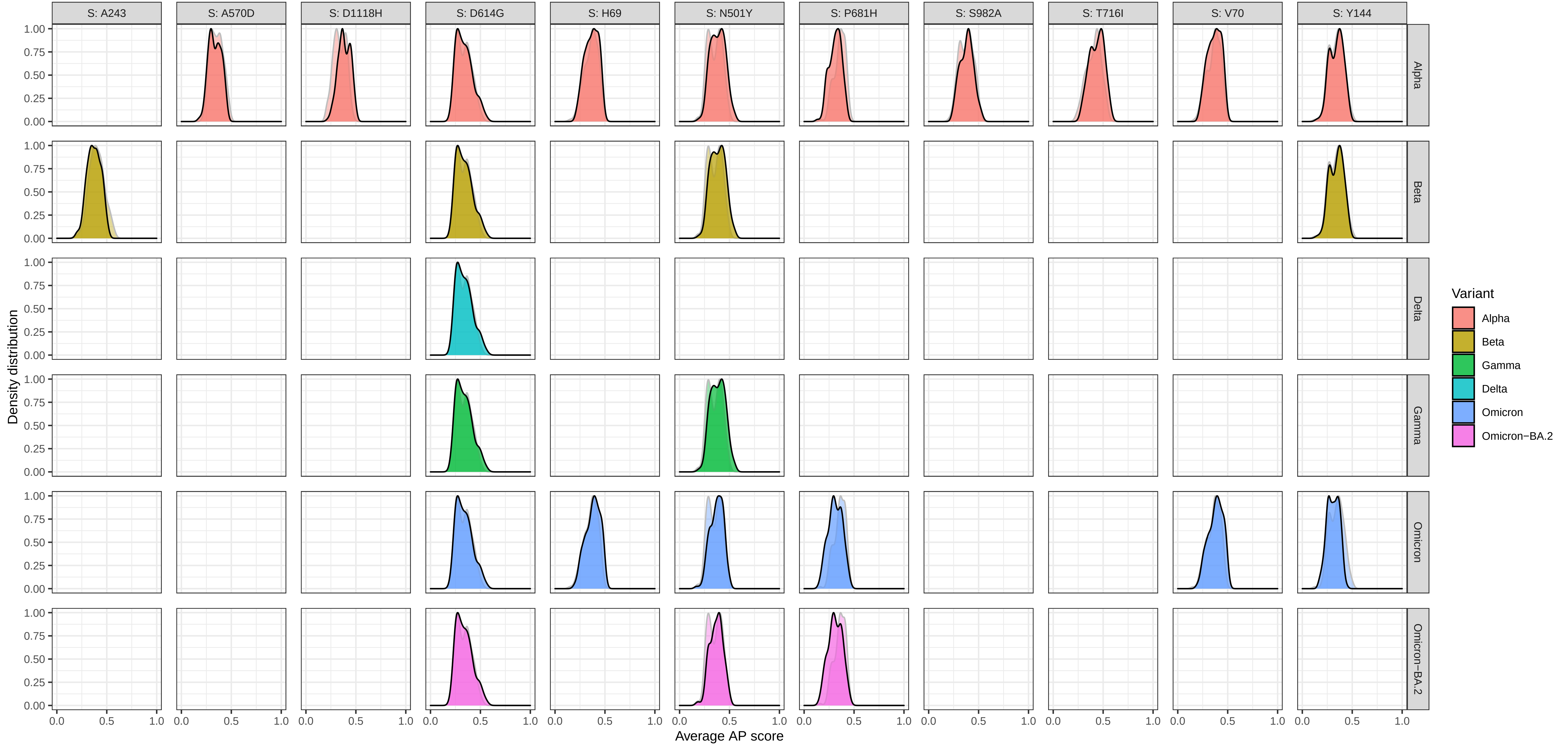

Supplementary Figure 6: Density of average AP scores at each mutation for a mutant (foreground) and wildtype (background) − protein S

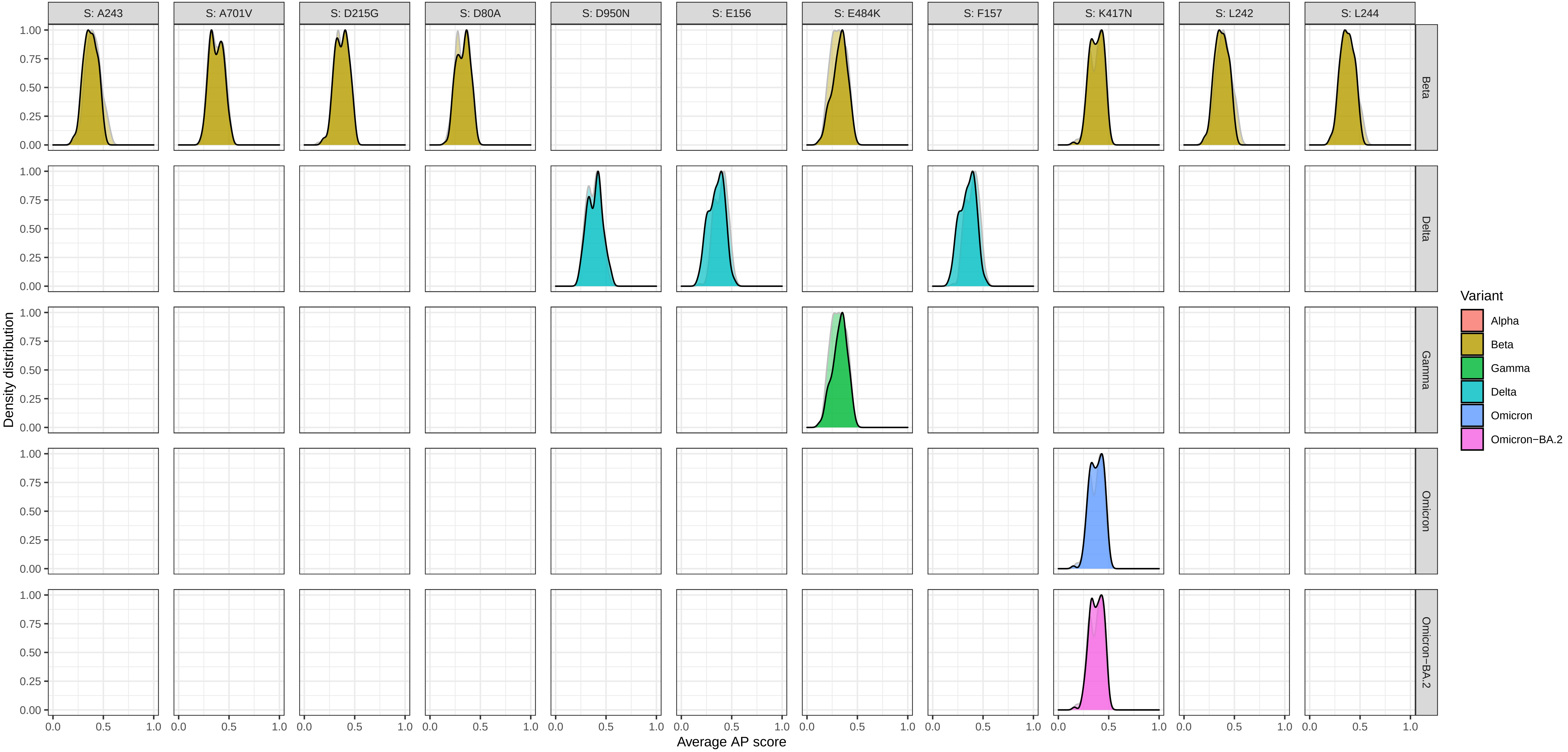

Supplementary Figure 6: Density of average AP scores at each mutation for a mutant (foreground) and wildtype (background) − protein S

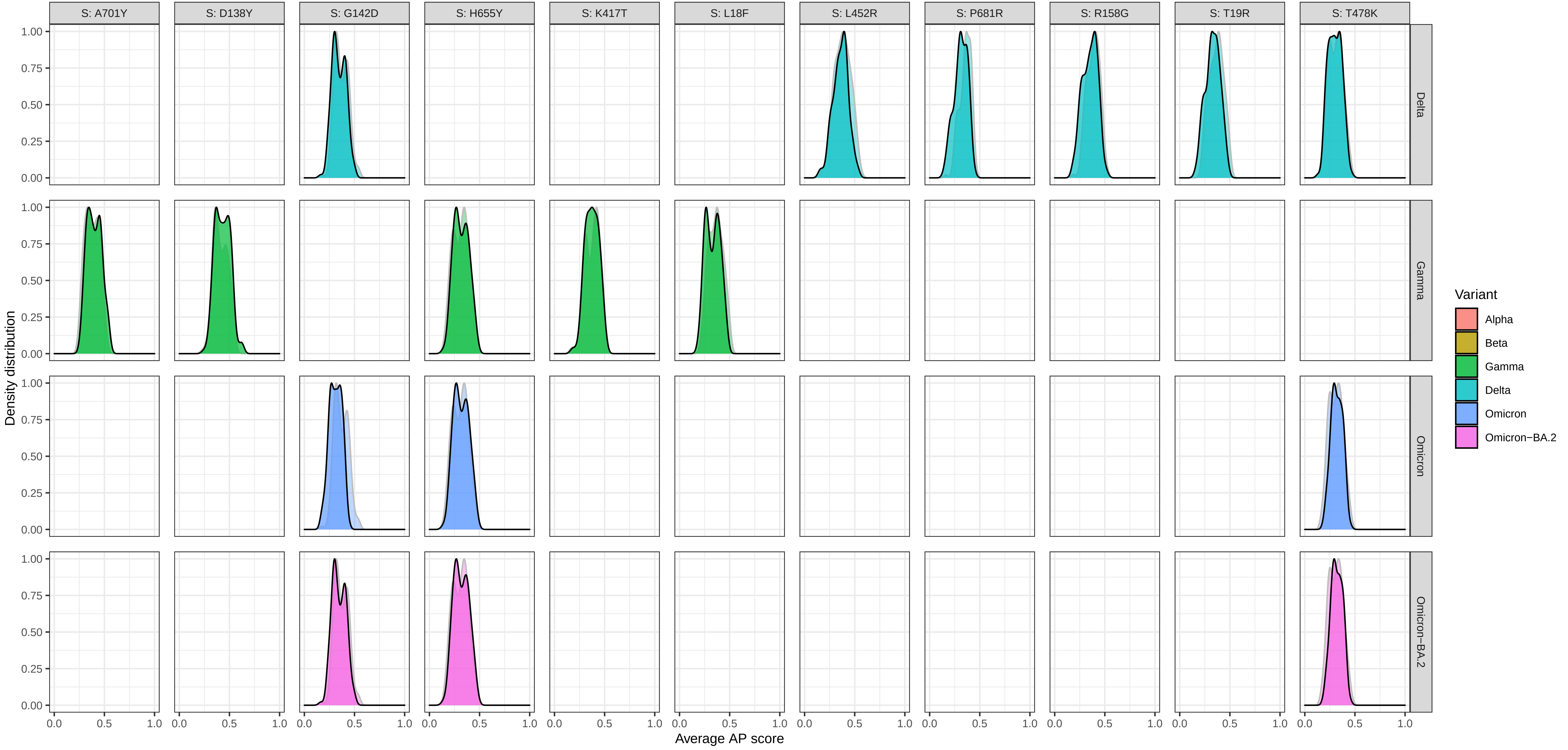

### Supplementary Figure 6: Density of average AP scores at each mutation for a mutant (foreground) and wildtype (background) − protein S

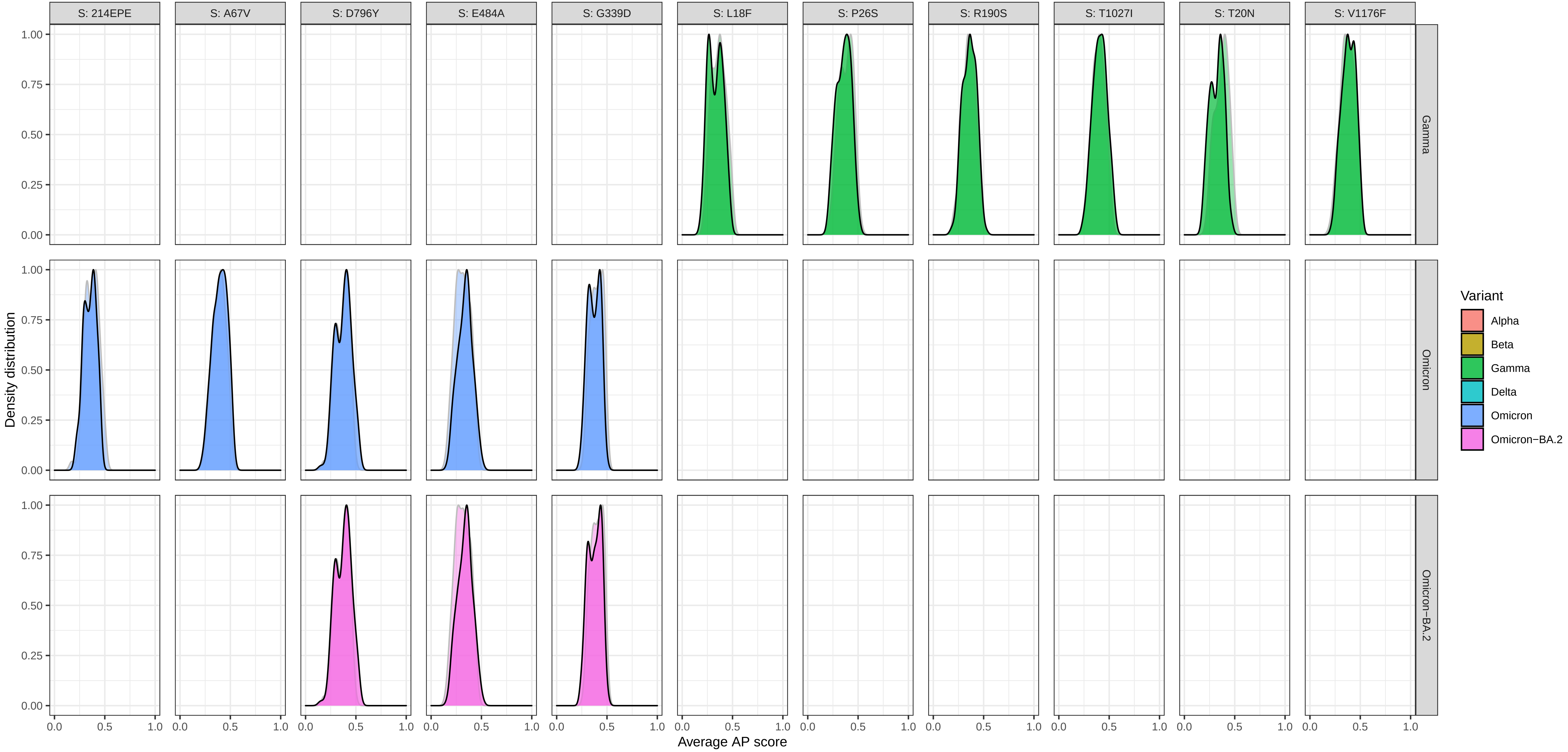

Supplementary Figure 6: Density of average AP scores at each mutation for a mutant (foreground) and wildtype (background) − protein S

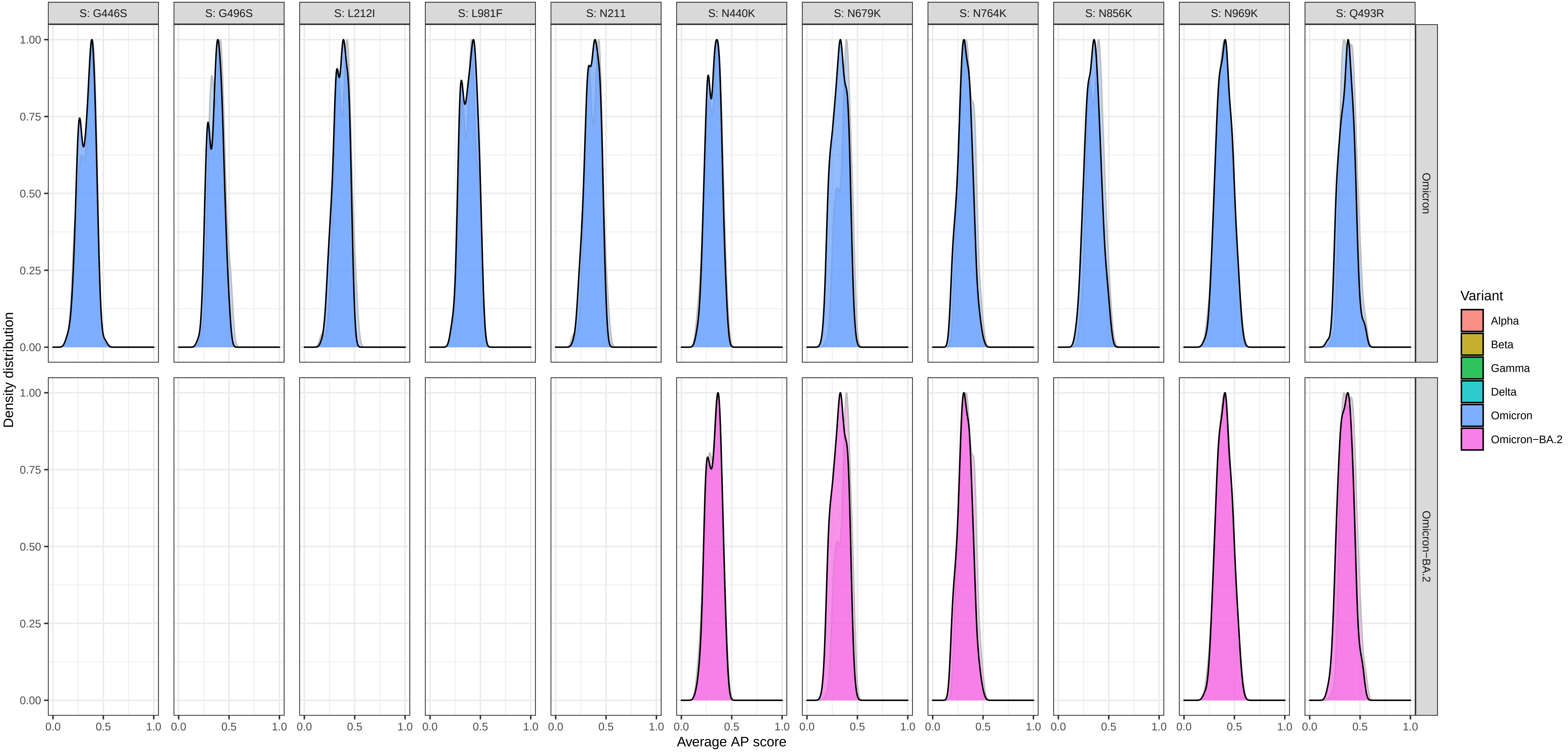

### Supplementary Figure 6: Density of average AP scores at each mutation for a mutant (foreground) and wildtype (background) − protein S

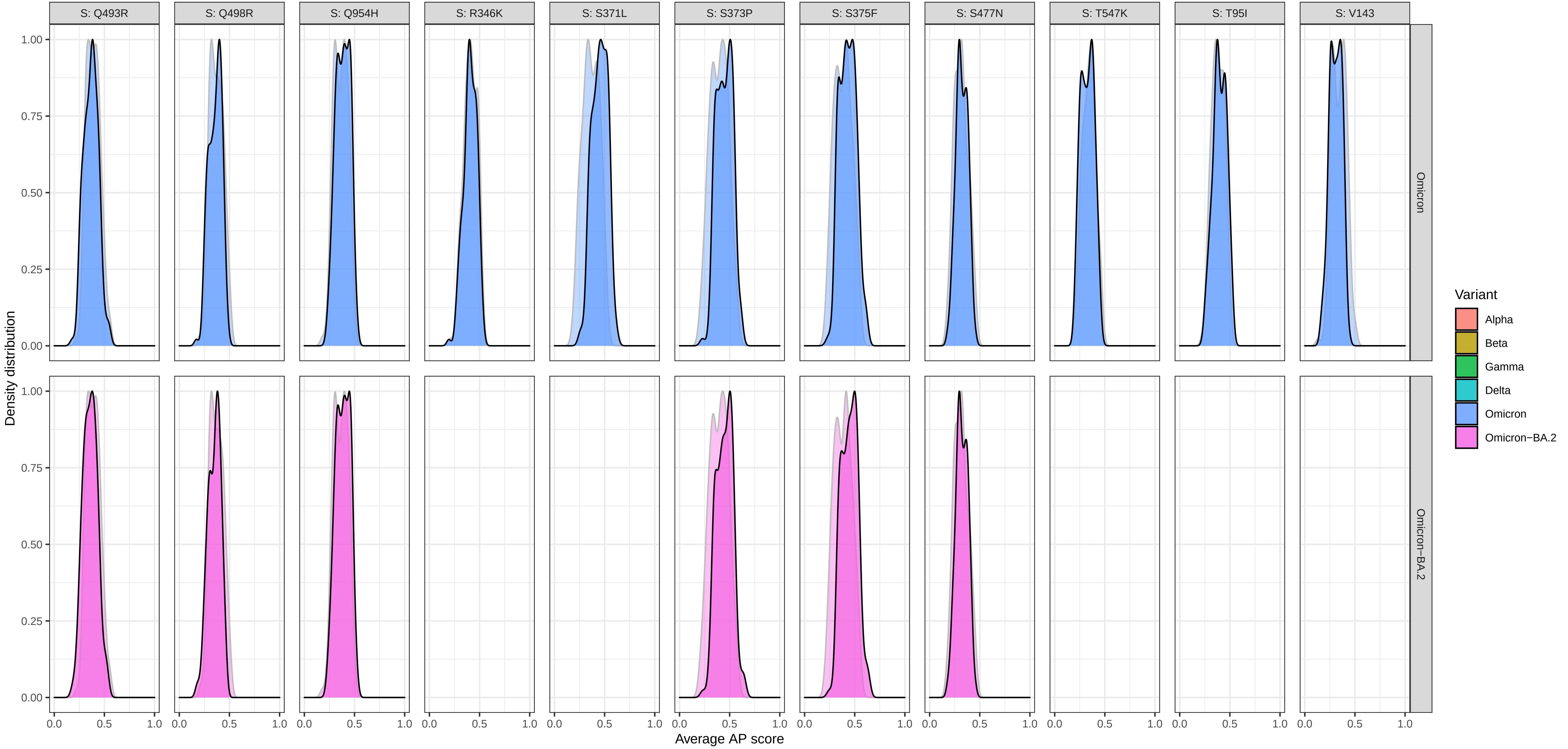

Supplementary Figure 6: Density of average AP scores at each mutation for a mutant (foreground) and wildtype (background) − protein S

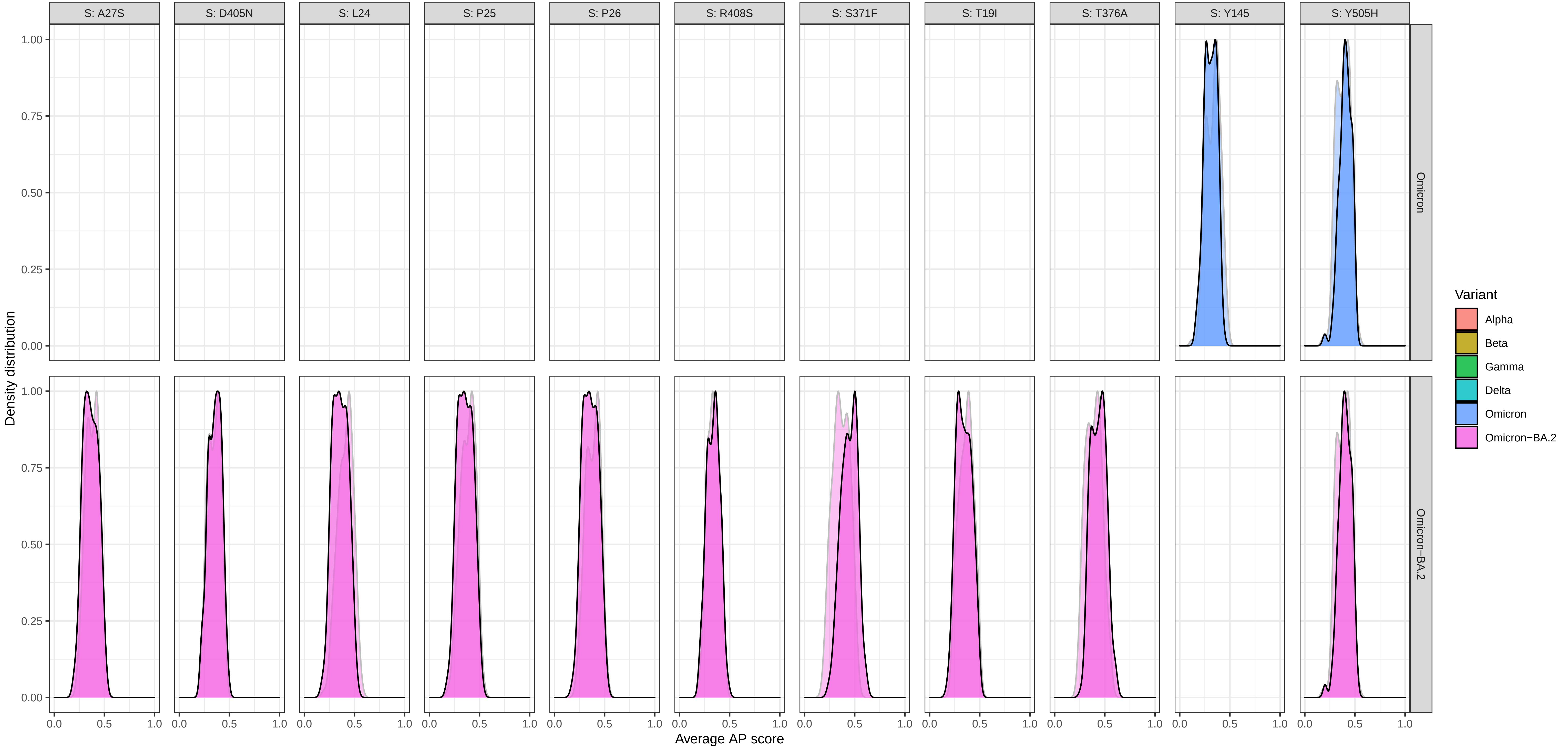

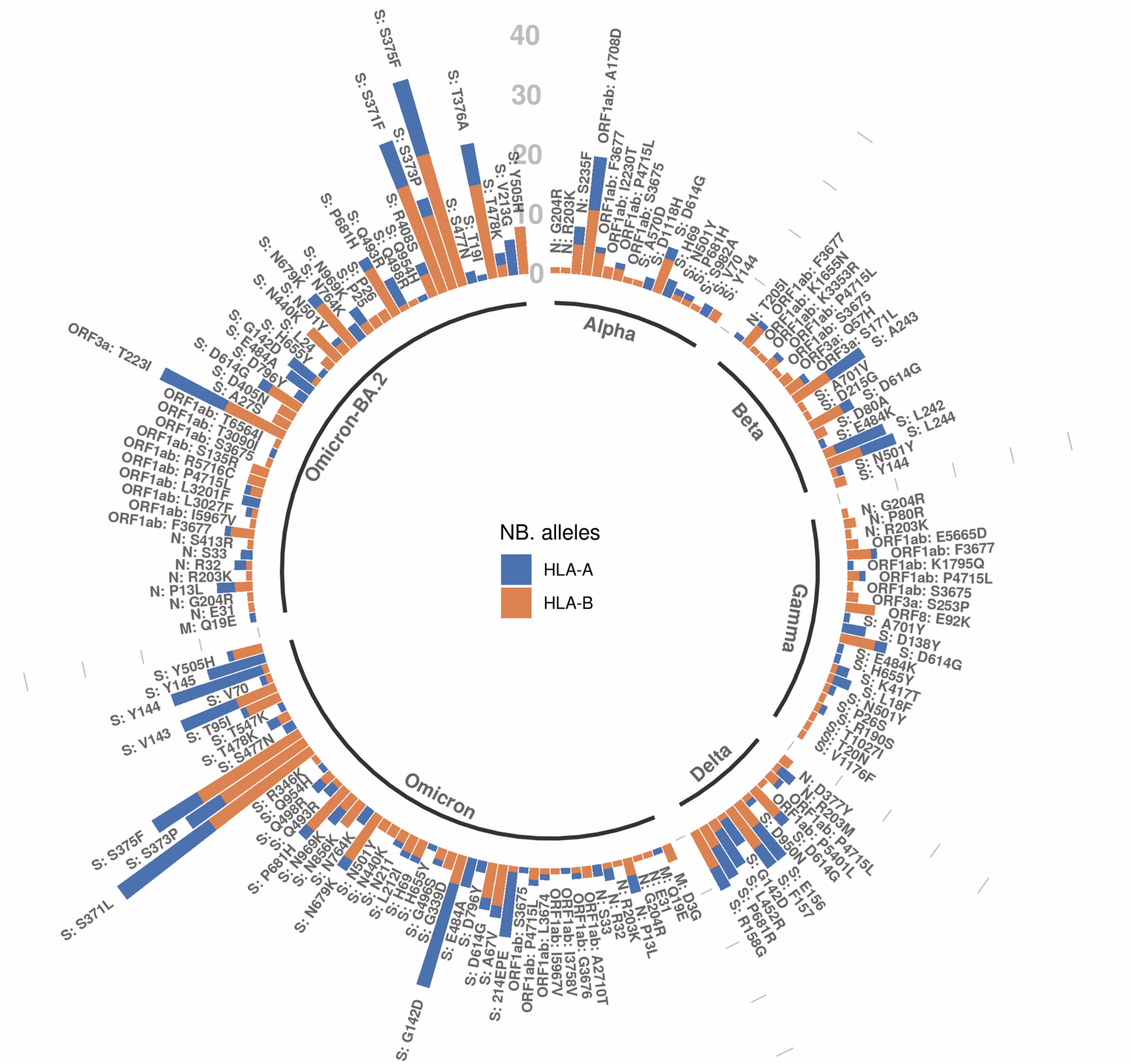


**Supplementary Figure 7:** This circular plot shows the number of HLA alleles per mutation for which there was a significant difference (p-value < 0.05) from the two-sided Kolmogorov-Smirnov (KS) test applied on the the variant and the wildtype AP score distribution. Only entries with an AP score above 0.5 in either the mutant or wildtype were considered for this plot. The label of each bar contains the protein name and the mutation details separated by colon. Each variant has its own group and the number of HLA-A alleles and HLA-B alleles is displayed separately for each mutation. The plot also includes the newly emerged Omicron-BA2 variant.

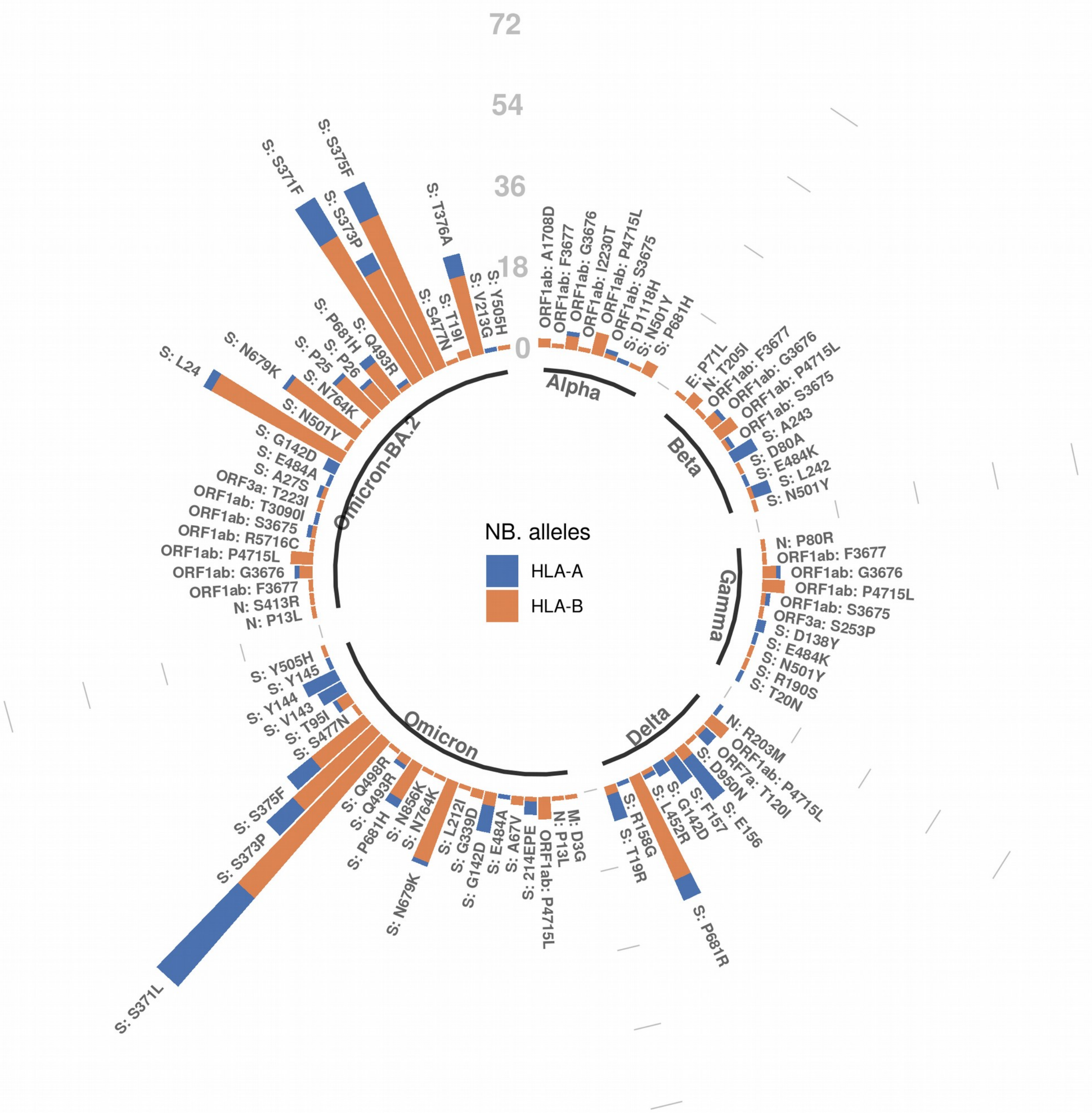


**Supplementary Figure 8:** This circular plot shows the number of HLA alleles per mutation for which there was a significant difference (p-value < 0.05) from the two-sided Kolmogorov-Smirnov (KS) test applied on the the variant and the wildtype AP score distribution. All AP scores (i.e., no prior filtering) were considered for this plot. The label of each bar contains the protein name and the mutation details separated by colon. Each variant has its own group and the number of HLA-A alleles and HLA-B alleles is displayed separately for each mutation. The plot also includes the newly emerged Omicron-BA2 variant.

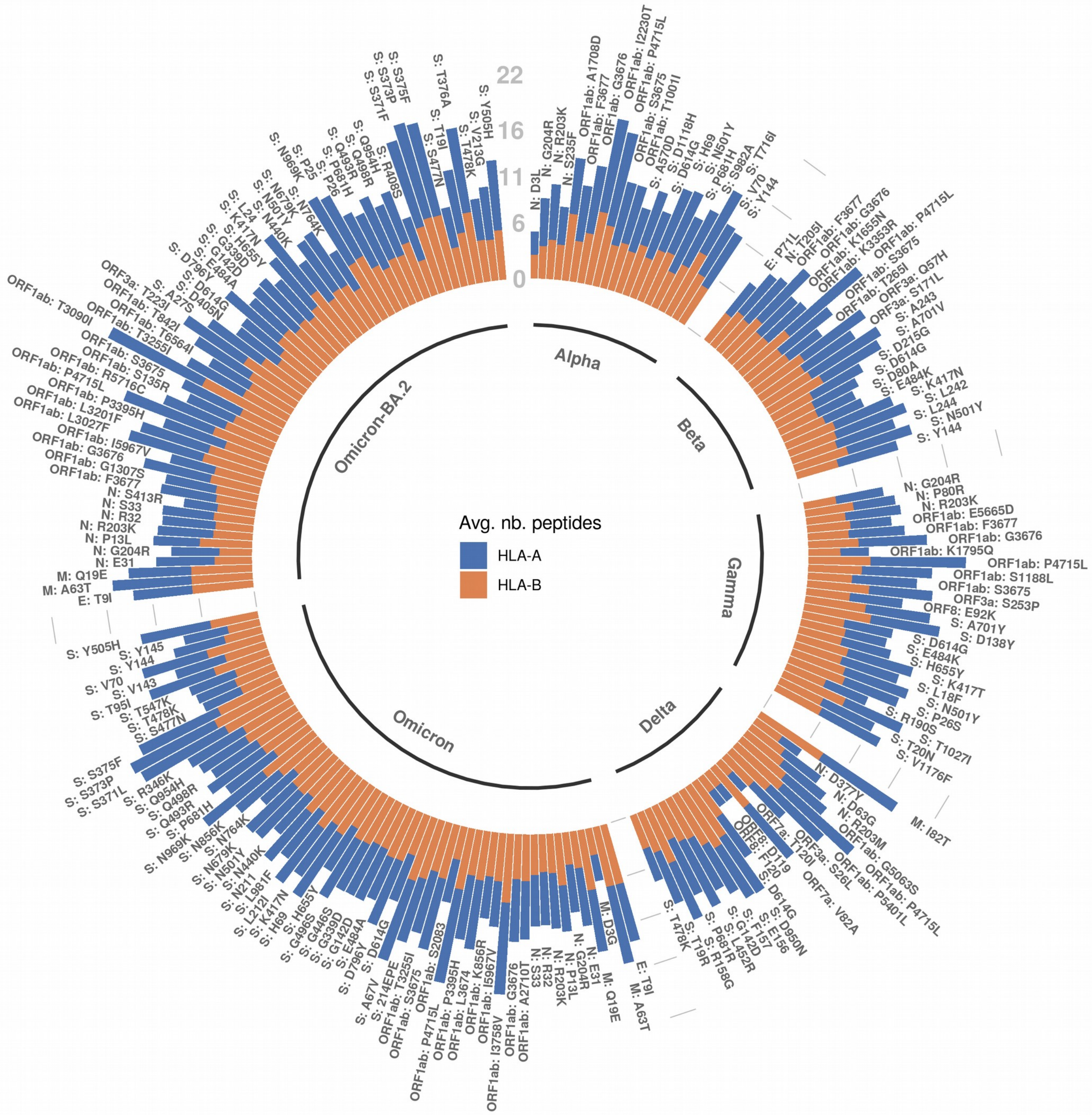


**Supplementary Figure 9:** This circular plot shows the average number of peptides across all HLA alleles for each of the mutations occurring in each variant. Only entries with an AP score above 0.5 in either the mutant or wildtype were considered for this plot. The label of each bar contains the protein name and the mutation details separated by colon. Each variant has its own group and the average across HLA-A alleles and HLA-B alleles is displayed separately for each mutation. This plot can give an indication about the resolution used for the KS two sided significance test. The plot also includes the newly emerged Omicron-BA2 variant.

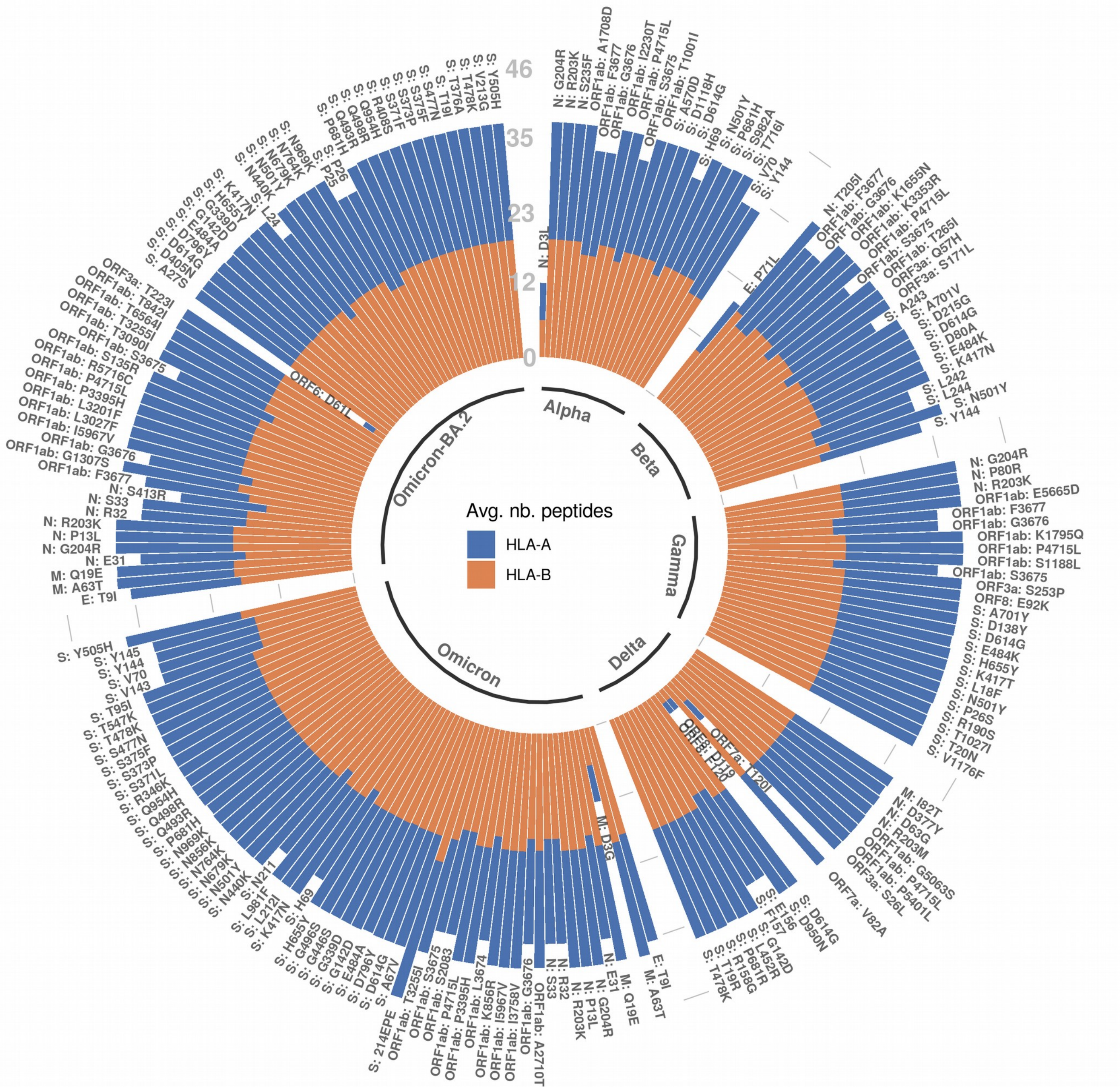


**Supplementary Figure 10:** This circular plot shows the average number of peptides across all HLA alleles for each of the mutations occurring in each variant. All AP scores (i.e., no prior filtering) were considered for this plot. The label of each bar contains the protein name and the mutation details separated by colon. Each variant has its own group and the average across HLA-A alleles and HLA-B alleles is displayed separately for each mutation. This plot can give an indication about the resolution used for the KS two sided significance test. The plot also includes the newly emerged Omicron-BA2 variant.
